# Distribution of methionine sulfoxide reductases in fungi and conservation of the free-methionine-*R*-sulfoxide reductase in multicellular eukaryotes

**DOI:** 10.1101/2021.02.26.433065

**Authors:** Hayat Hage, Marie-Noëlle Rosso, Lionel Tarrago

## Abstract

Methionine, either as a free amino acid or included in proteins, can be oxidized into methionine sulfoxide (MetO), which exists as *R* and *S* diastereomers. Almost all characterized organisms possess thiol-oxidoreductases named methionine sulfoxide reductase (Msr) enzymes to reduce MetO back to Met. MsrA and MsrB reduce the *S* and *R* diastereomers of MetO, respectively, with strict stereospecificity and are found in almost all organisms. Another type of thiol-oxidoreductase, the free-methionine-*R*-sulfoxide reductase (fRMsr), identified so far in prokaryotes and a few unicellular eukaryotes, reduces the *R* MetO diastereomer of the free amino acid. Moreover, some bacteria possess molybdenum-containing enzymes that reduce MetO, either in the free or protein-bound forms. All these Msrs play important roles in the protection of organisms against oxidative stress. Fungi are heterotrophic eukaryotes that colonize all niches on Earth and play fundamental functions, in organic matter recycling, as symbionts, or as pathogens of numerous organisms. However, our knowledge on fungal Msrs is still limited. Here, we performed a survey of *msr* genes in almost 700 genomes across the fungal kingdom. We show that most fungi possess one gene coding for each type of methionine sulfoxide reductase: MsrA, MsrB, and fRMsr. However, several fungi living in anaerobic environments or as obligate intracellular parasites were devoid of *msr* genes. Sequence inspection and phylogenetic analyses allowed us to identify non-canonical sequences with potentially novel enzymatic properties. Finaly, we identified several ocurences of *msr* horizontal gene transfer from bacteria to fungi.

**Highlights:**

- Free and protein-bound methionine can be oxidized into methionine sulfoxide (MetO).
- Methionine sulfoxide reductases (Msr) reduce MetO in most organisms.
- Sequence characterization and phylogenomics revealed strong conservation of Msr in fungi.
- fRMsr is widely conserved in unicellular and multicellular fungi.
- Some *msr* genes were acquired from bacteria via horizontal gene transfers.

## 1. Introduction

Life in the presence of dioxygen necessarily exposes the biological systems to oxidant molecules. Due to its high reactivity, dioxygen can be converted to reactive oxygen species (ROS), which play key roles in physiological and pathological contexts through redox modifications of macromolecules [1]. Met, either as a free amino acid or as a residue included in a protein, can be oxidized into Met sulfoxide (MetO) by the addition of an oxygen atom to the sulfur of the lateral chain. MetO exists as diastereomer *R* or *S* (Met-*R*-O and Met-*S*-O, respectively) [2], and can be reduced back to Met by the action of oxidoreductases called methionine sulfoxide reductases (Msr). The two main types are the (seleno)thiol-containing MsrA and MsrB, which display strict stereoselectivities toward the *S*- and the *R*-diastereomer of MetO, respectively [3–5]. Whereas MsrA reduces free MetO and protein-bound MetO with similar catalytic efficiencies, MsrB generally only reduces efficiently the protein-bound MetO [6]. Another enzyme, fRMsr (free methionine-(R)-sulfoxide reductase) specifically reduces the free form of Met-*R*-O [7–9]. Some bacteria also possess several molybdoenzymes that reduce exclusively the free MetO [10, 11] or both the free and protein-bound MetO [12–15]. Despite the lack of sequence and structure similarities, MsrA, MsrB and fRMsr generally catalyze the reduction of MetO using a similar 3-steps-mechanism [9, 16] : i) a *‘catalytic’* Cys (or, less frequently, a selenocysteine, Sec) reduces the target MetO and is converted into a sulfenic (or selenic) acid [17], ii) an internal *‘resolving’* Cys reduces it through the formation of an intramolecular disulfide bond, and finally iii) the oxidized Msr is regenerated through disulfide exchange with a thioredoxin [18]. Variations of this mechanism exist, with some Msrs devoid of any resolving Cys, in which the sulfenic acid is directly reduced by an external reducer [19–21]. Alternatively, in other Msrs, a disulfide exchange occurs with a second internal resolving Cys before regeneration by the thioredoxin [7,22,23]. Genome mining analyses indicated that *msrA* and *msrB* genes originated from early prokaryotes and are now present in all organisms across the tree of life, with few exceptions [24–26]. Most organisms have few *msr* genes, generally one of each type [24–26], but this number is generally higher in plants (e.g. up to 5 *msrA* and 9 *msrB* genes in *Arabidopsis thaliana* [25]). Moreover, some bacteria encode a bifunctional MsrA/MsrB fusion and some others lack a MsrB, but no organism of any kind was so far described with only a MsrB [24, 26]. Finally, only very few organisms do not possess any *msr* gene at all, such as a few endosymbiotic or obligatory parasitic bacteria and some archaea [24, 26]. fRMsr were so far only reported in bacteria and a few unicellular eukaryotes [7–9]. Most interestingly, the only eukaryote for which the absence of *msr* gene was observed is the fungus *Encephalitozoon cuniculi*, a microsporidium having a remarkably reduced genome (∼2.9 Mb) and living as intracellular parasite of mammals [24, 27]. The conservation of typical MsrAs and MsrBs in almost all known organisms argues for the critical role of MetO reduction in cellular metabolism, and numerous studies showed that Msrs are involved in the protection against oxidative stress and the regulation of protein functions. Schematically, Msr protective roles against oxidative injuries occur through two main functions: i) the repair of oxidized proteins, and ii) an antioxidant function through ROS scavenging by cyclic Met oxidation and reduction. Moreover, the reversible Met oxidation was shown to act as a post-translational modification responsible for the activation of enzymes and transcription factors or the regulation of protein-protein interactions [28–30]. The role of fRMsr has been far less studied, but it is suspected to have an antioxidant function by reducing free MetO and maintaining the pool of Met available for protein synthesis and for the production of sulfur-containing metabolites [8, 31]. Overall, these functions were well established in animals, bacteria and plants, and have recently been reviewed [32–36].

Fungi are heterotrophic eukaryotes that colonize virtually all niches on Earth and exist as unicellular or multicellular organisms. They can have numerous lifestyles, either as free living organisms playing key roles in organic and inorganic matter cycling, or as symbionts or pathogens with crucial impacts on plant and animal health [37]. As any other organism living in aerobic conditions and exposed to environmental constraints, fungi are exposed to oxidative constraints and protein oxidation [38]. However, the effects of Met oxidation and the roles of Msrs were largely overlooked in these eukaryotes. The Msr system was mainly characterized in the yeast *Saccharomyces cerevisiae*, which possesses one Msr of each type [6, 8]. In *S. cerevisiae*, MsrA is located in the cytosol, MsrB is found in mitochondria and in the cytosol and fRMsr is located both in the cytosol and the nucleus [39, 40]. Genetic manipulations have shown that these Msrs are involved in the protection against oxidative stress and in maintaining the yeast lifespan [6,8,41,42]. Consistently, the overexpression of *msrA* in the basidiomycete *Pleurotus ostreatus* and of *msrB* in the yeast *Schizosaccharomyces pombe* increased the resistance to oxidative constraints [43, 44]. Moreover, the filamentous fungus *Aspergillus nidulans* possesses one *msrA* and one *msrB* genes, and the deletion of one or both genes increased the sensitivity of the fungus to oxidative treatments [45]. Few proteins with oxidized Met residues have been characterized in fungi, but remarkably interesting findings were obtained. For instance, in *S. cerevisiae*, the reversible oxidation of Met regulates the oligomerization state of the ataxin-2 protein and the activity of the co-chaperone Fes1 [40, 46]. Moreover, Met oxidation enhanced the activity of an α-galactosidase in *Trichoderma reesei* [47]. Finally, in *A. nidulans*, it was demonstrated that the nuclear localization of the nitrate-responsive transcription factor NirA was actively regulated through cyclic Met oxidation [48]. These data indicate that, as for other organisms, fungal Msrs certainly play key roles under many conditions of oxidative stress, such as biotic interactions or abiotic constraints.

In this study, we searched for *msr* genes in about 700 fungal genomes. We show that the great majority of fungi have one *msrA* and one *msrB* genes. Moreover, we identified *fRmsr* genes in almost all the fungi we analyzed and thereby demonstrate that the enzyme is conserved in these multicellular eukaryotes. Finally, using a phylogenetic analysis and a close inspection of the sequence features, we identified fungal Msrs with unusual sequence characteristics and uncovered horizontal gene transfers from bacteria to fungi.

## 2. Material and methods

### 2.1. Search for Msr homologs in fungi

The protein sequence of MsrA (Uniprot accession # C8Z745), MsrB (Uniprot accession # P25566) and fRMsr (Uniprot accession # P36088) from *S. cerevisiae*, MsrP (Uniprot accession # P76342) and BisC (Uniprot accession # P20099) from *Escherichia coli*, TorZ (Uniprot accession # P44798) from *Haemophilus influenzae*, and DorA (Uniprot accession # Q57366) from *Rhodobacter sphaeroides* were used as BLASTP and TBLASTN [49] queries to identify *msr* genes in 683 genomes available in the MycoCosm database [50] (https://mycocosm.jgi.doe.gov/mycocosm/home).

### 2.2. List of all explored fungal genomes

*Aaosphaeria arxii CBS 175.79 v1.0; Amniculicola lignicola CBS 123094 v1.0; Ampelomyces quisqualis HMLAC05119 v1.0; Aplosporella prunicola CBS 121.167 v1.0; Aulographum hederae v2.0; Bimuria novae-zelandiae CBS 107.79 v1.0; Byssothecium circinans CBS 675.92 v1.0; Cercospora zeae-maydis v1.0; Clathrospora elynae CBS 161.51 v1.0; Cucurbitaria berberidis CBS 394.84 v1.0; Decorospora gaudefroyi v1.0; Delitschia confertaspora ATCC 74209 v1.0; Delphinella strobiligena CBS 735.71 v1.0; Didymella exigua CBS 183.55 v1.0; Dissoconium aciculare v1.0; Dothidotthia symphoricarpi v1.0; Elsinoe ampelina CECT 20119 v1.0; Eremomyces bilateralis CBS 781.70 v1.0; Hortaea acidophila CBS 113389 v1.0; Karstenula rhodostoma CBS 690.94 v1.0; Lentithecium fluviatile v1.0; Lindgomyces ingoldianus ATCC 200398 v1.0; Lineolata rhizophorae ATCC 16933 v1.0; Lizonia empirigonia CBS 542.76 v1.0; Lophiostoma macrostomum v1.0; Lophiotrema nucula CBS 627.86 v1.0; Lophium mytilinum CBS 269.34 v1.0; Macroventuria anomochaeta CBS 525.71 v1.0; Massarina eburnea CBS 473.64 v1.0; Massariosphaeria phaeospora CBS 611.86 v1.0; Melanomma pulvis-pyrius v1.0; Microthyrium microscopicum CBS 115976 v1.0; Myriangium duriaei CBS 260.36 v1.0; Mytilinidion resinicola CBS 304.34 v1.0; Ophiobolus disseminans CBS 113818 v1.0; Patellaria atrata v1.0; Phoma tracheiphila IPT5 v1.0; Piedraia hortae CBS 480.64 v1.1; Pleomassaria siparia v1.0; Polychaeton citri v1.0; Polyplosphaeria fusca CBS 125425 v1.0; Pseudovirgaria hyperparasitica CBS 121739 v1.0; Rhizodiscina lignyota CBS 133067 v1.0; Saccharata proteae CBS 121410 v1.0; Setomelanomma holmii CBS 110217 v1.0; Sporormia fimetaria v1.0; Teratosphaeria nubilosa CBS 116005 v1.0; Tothia fuscella CBS 130266 v1.0; Trematosphaeria pertusa CBS 122368 v1.0; Trichodelitschia bisporula CBS 262.69 v1.0; Verruculina enalia CBS 304.66 v1.0; Viridothelium virens v1.0; Westerdykella ornata CBS 379.55 v1.0; Zasmidium cellare ATCC 36951 v1.0; Zopfia rhizophila v1.0* [51]*; Acaromyces ingoldii MCA 4198 v1.0; Ceraceosorus guamensis MCA 4658 v1.0; Jaminaea sp. MCA 5214 v1.0; Meira miltonrushii MCA 3882 v1.0; Pseudomicrostroma glucosiphilum gen et sp. nov. MCA 4718 v1.0; Testicularia cyperi MCA 3645 v1.0; Tilletiopsis washingtonensis MCA 4186 v1.0; Violaceomyces palustris SA 807 v1.0* [52]*; Acidomyces richmondensis BFW; Acidomyces richmondensis BFW* [53]*; Acremonium chrysogenum ATCC 11550* [54]*; Agaricus bisporus var bisporus (H97) v2.0; Agaricus bisporus var. burnettii JB137-S8; Gigaspora rosea v1.0; Rhizophagus cerebriforme DAOM 227022 v1.0; Rhizophagus diaphanus v1.0* [55]*; Amanita muscaria Koide v1.0; Gymnopus luxurians v1.0; Hebeloma cylindrosporum h7 v2.0; Hydnomerulius pinastri v2.0; Hypholoma sublateritium v1.0; Laccaria amethystina LaAM-08-1 v2.0; Paxillus adelphus Ve08.2h10 v2.0; Paxillus involutus ATCC 200175 v1.0; Piloderma croceum F 1598 v1.0; Pisolithus microcarpus 441 v1.0; Pisolithus tinctorius Marx 270 v1.0; Plicaturopsis crispa v1.0; Scleroderma citrinum Foug A v1.0; Sebacina vermifera MAFF 305830 v1.0; Sphaerobolus stellatus v1.0; Suillus luteus UH-Slu-Lm8-n1 v3.0; Tulasnella calospora AL13/4D v1.0; Oidiodendron maius Zn v1.0* [56]*; Amanita thiersii Skay4041 v1.0* [57]*; Ambrosiozyma philentoma NRRL Y-7523; Candida boidinii NRRL Y-2332; Citeromyces matritensis NRRL Y-2407; Nakazawaea wickerhamii NRRL Y-2563; Peterozyma xylosa NRRL Y-12939; Saccharomycopsis capsularis NRRL Y-17639; Saturnispora dispora NRRL Y-1447* [58]*; Amorphotheca resinae v1.0; Meliniomyces bicolor E v2.0; Meliniomyces variabilis F v1.0; Rhizoscyphus ericae UAMH 7357 v1.0* [59]*; Anaeromyces robustus v1.0; Neocallimastix californiae G1 v1.0; Piromyces finnis v3.0; Piromyces sp. E2 v1.0* [60]*; Antonospora locustae HM-2013* [61]*; Armillaria cepistipes B5; Armillaria gallica 21-2 v1.0; Armillaria ostoyae C18/9; Armillaria solidipes 28-4 v1.0* [62]*; Armillaria mellea DSM 3731* [63]*; Arthrobotrys oligospora ATCC 24927* [64]*; Ascobolus immersus RN42 v1.0; Choiromyces venosus 120613-1 v1.0; Morchella importuna CCBAS932 v1.0; Terfezia boudieri ATCC MYA-4762 v1.1; Tuber aestivum var. urcinatum v1.0; Tuber borchii Tbo3840 v1.0; Tuber magnatum v1.0* [65]; *Ascochyta rabiei ArDII* [66]; *Ascocoryne sarcoides NRRL50072* [67]*; Ascodesmis nigricans CBS 389.68 v1.0* [68]*; Aspergillus aculeatinus CBS 121060 v1.0; Aspergillus brunneoviolaceus CBS 621.78 v1.0; Aspergillus costaricaensis CBS 115574 v1.0; Aspergillus ellipticus CBS 707.79 v1.0; Aspergillus eucalypticola CBS 122712 v1.0; Aspergillus fijiensis CBS 313.89 v1.0; Aspergillus heteromorphus CBS 117.55 v1.0; Aspergillus homomorphus CBS 101889 v1.0; Aspergillus ibericus CBS 121593 v1.0; Aspergillus indologenus CBS 114.80 v1.0; Aspergillus japonicus CBS 114.51 v1.0; Aspergillus neoniger CBS 115656 v1.0; Aspergillus niger (lacticoffeatus) CBS 101883 v1.0; Aspergillus niger (phoenicis Corda) Thom ATCC 13157 v1.0; Aspergillus niger NRRL3; Aspergillus niger van Tieghem ATCC 13496 v1.0; Aspergillus piperis CBS 112811 v1.0; Aspergillus saccharolyticus JOP 1030-1 v1.0; Aspergillus sclerotiicarbonarius CBS 121057 v1.0; Aspergillus sclerotioniger CBS115572 v1.0; Aspergillus uvarum CBS 121591 v1.0; Aspergillus vadensis CBS 113365 v1.0; Aspergillus violaceofuscus CBS 115571 v1.0; Aspergillus welwitschiae CBS139.54b v1.0* [69]*; Aspergillus aculeatus ATCC16872 v1.1; Aspergillus brasiliensis v1.0; Aspergillus carbonarius ITEM 5010 v3; Aspergillus glaucus v1.0; Aspergillus luchuensis CBS 106.47 v1.0; Aspergillus sydowii CBS 593.65 v1.0; Aspergillus tubingensis v1.0; Aspergillus versicolor v1.0; Aspergillus wentii v1.0; Aspergillus zonatus v1.0; Penicillium chrysogenum v1.0* [70]*; Aspergillus bombycis NRRL 26010* [71]*; Aspergillus calidoustus* [72]*; Aspergillus campestris IBT 28561 v1.0; Aspergillus candidus CBS 102.13 v1.0; Aspergillus novofumigatus IBT 16806 v1.0; Aspergillus ochraceoroseus IBT 24754 v1.0; Aspergillus steynii IBT 23096 v1.0; Aspergillus taichungensis IBT 19404 v1.0; Aspergillus albertensis v1.0; Aspergillus alliaceus CBS 536.65 v1.0; Aspergillus arachidicola v1.0; Aspergillus avenaceus IBT 18842 v1.0; Aspergillus bertholletius IBT 29228 v1.0; Aspergillus caelatus CBS 763.97 v1.0; Aspergillus coremiiformis CBS 553.77 v1.0; Aspergillus leporis CBS 151.66 v1.0; Aspergillus minisclerotigenes CBS 117635 v1.0; Aspergillus nomius IBT 12657 v1.0; Aspergillus novoparasiticus CBS 126849 v1.0; Aspergillus parasiticus CBS 117618 v1.0; Aspergillus parvisclerotigenus CBS 121.62 v1.0; Aspergillus pseudocaelatus CBS 117616 v1.0; Aspergillus pseudonomius CBS 119388 v1.0; Aspergillus pseudotamarii CBS 117625 v1.0; Aspergillus sergii CBS 130017 v1.0; Aspergillus tamarii CBS 117626 v1.0; Aspergillus transmontanensis CBS 130015 v1.0* [73]*; Aspergillus clavatus NRRL 1 from AspGD; Aspergillus flavus NRRL3357; Aspergillus nidulans; Aspergillus oryzae RIB40; Aspergillus terreus NIH 2624; Neosartorya fischeri NRRL 181* [74]*; Aspergillus cristatus GZAAS20.1005* [75]*; Aspergillus fumigatus A1163* [76]*; Aspergillus fumigatus Af293 from AspGD* [77]*; Aspergillus kawachii IFO 4308* [78]*; Aspergillus niger ATCC 1015 v4.0* [79]*; Aspergillus niger CBS 513.88* [80]*; Aspergillus nomius NRRL 13137; Aspergillus ochraceoroseus SRRC1432; Aspergillus rambellii SRRC1468* [81]*; Aspergillus udagawae IFM 46973* [82]*; Aureobasidium pullulans var. melanogenum CBS 110374; Aureobasidium pullulans var. namibiae CBS 147.97; Aureobasidium pullulans var. pullulans EXF-150; Aureobasidium pullulans var. subglaciale EXF-2481* [83]*; Auricularia subglabra v2.0; Coniophora puteana v1.0; Dichomitus squalens LYAD-421 SS1 v1.0; Fomitiporia mediterranea v1.0; Fomitopsis pinicola FP-58527 SS1 v3.0; Gloeophyllum trabeum v1.0; Punctularia strigosozonata v1.0; Stereum hirsutum FP-91666 SS1 v1.0; Trametes versicolor v1.0; Wolfiporia cocos MD-104 SS10 v1.0; Dacryopinax primogenitus DJM 731 SSP1 v1.0; Tremella mesenterica Fries v1.0* [84]*; Auriculariopsis ampla NL-1724 v1.0* [85]*; Baudoinia compniacensis UAMH 10762 (4089826) v1.0; Cochliobolus heterostrophus C4 v1.0; Cochliobolus heterostrophus C5 v2.0; Cochliobolus lunatus m118 v2.0; Cochliobolus sativus ND90Pr v1.0; Hysterium pulicare; Rhytidhysteron rufulum; Septoria musiva SO2202 v1.0; Septoria populicola v1.0; Setosphaeria turcica Et28A v2.0* [86]*; Beauveria bassiana ARSEF 2860* [87]*; Bjerkandera adusta v1.0; Ganoderma sp. 10597 SS1 v1.0; Phlebia brevispora HHB-7030 SS6 v1.0; Neofusicoccum parvum UCRNP2; Eutypa lata UCREL1; Phaeoacremonium aleophilum UCRPA7* [88]*; Blastobotrys (Arxula) adeninivorans* [89]*; Blastomyces dermatitidis SLH14081* [90]*; Blumeria graminis f. sp. hordei DH14; Blumeria graminis f. sp. hordei Race1* [91]*; Blumeria graminis f. sp. tritici 96224* [92]*; Botryobasidium botryosum v1.0; Galerina marginata v1.0; Jaapia argillacea v1.0; Pleurotus ostreatus PC15 v2.0; Ascoidea rubescens NRRL Y17699 v1.0; Babjeviella inositovora NRRL Y-12698 v1.0; Candida arabinofermentans NRRL YB-2248 v1.0; Candida tanzawaensis NRRL Y-17324 v1.0; Cyberlindnera jadinii NRRL Y-1542 v1.0; Hanseniaspora valbyensis NRRL Y-1626 v1.1; Hyphopichia burtonii NRRL Y-1933 v1.0; Lipomyces starkeyi NRRL Y-11557 v1.0; Metschnikowia bicuspidata NRRL YB-4993 v1.0; Nadsonia fulvescens var. elongata DSM 6958 v1.0; Ogataea polymorpha NCYC 495 leu1.1 v2.0; Pachysolen tannophilus NRRL Y-2460 v1.2; Pichia membranifaciens v2.0; Tortispora caseinolytica Y-17796 v1.0; Wickerhamomyces anomalus NRRL Y-366-8 v1.0; Saitoella complicata NRRL Y-17804 v1.0* [93]*; Botryosphaeria dothidea* [94]*; Botrytis cinerea v1.0* [95]*; Byssochlamys spectabilis No. 5* [96]*; Candida albicans SC5314* [97]*; Candida tenuis NRRL Y-1498 v1.0; Spathaspora passalidarum NRRL Y-27907 v2.0* [98]*; Capronia coronata CBS 617.96; Capronia epimyces CBS 606.96; Capronia semiimmersa CBS27337; Cladophialophora bantiana CBS 173.52; Cladophialophora carrionii CBS 160.54; Cladophialophora immunda CBS83496; Cladophialophora psammophila CBS 110553; Cladophialophora yegresii CBS 114405; Cyphellophora europaea CBS 101466; Exophiala aquamarina CBS 119918; Exophiala mesophila CBS40295; Exophiala oligosperma CBS72588; Exophiala sideris CBS121828; Exophiala spinifera CBS89968; Exophiala xenobiotica CBS118157; Fonsecaea multimorphosa CBS 102226; Fonsecaea pedrosoi CBS 271.37; Coniosporium apollinis CBS 100218; Verruconis gallopava* [99]*; Cenococcum geophilum 1.58 v2.0; Glonium stellatum CBS 207.34 v1.0; Lepidopterella palustris v1.0* [100]*; Ceriporiopsis (Gelatoporia) subvermispora B* [101]*; Chaetomium globosum v1.0* [102]*; Chaetomium thermophilum var thermophilum DSM 1495* [103]*; Cladonia grayi Cgr/DA2myc/ss v2.0* [104]*; Cladosporium fulvum v1.0; Dothistroma septosporum NZE10 v1.0* [105]*; Cladosporium sphaerospermum UM 843* [106]*; Clavispora lusitaniae ATCC 42720; Lodderomyces elongisporus NRRL YB-4239; Meyerozyma guilliermondii ATCC 6260* [107]*; Coccidioides immitis RS ; Coccidioides posadasii C735 delta SOWgp; Histoplasma capsulatum NAm1; Uncinocarpus reesii UAMH 1704* [108]*; Cochliobolus carbonum 26-R-13 v1.0; Cochliobolus miyabeanus ATCC 44560 v1.0; Cochliobolus victoriae FI3 v1.0* [109]*; Colletotrichum chlorophyti NTL11* [110]*; Colletotrichum fioriniae PJ7* [111]*; Colletotrichum graminicola M1.001* [112]*; Colletotrichum higginsianum IMI 349063* [113]*; Colletotrichum incanum MAFF 238712* [114]*; Colletotrichum nymphaeae SA-01; Colletotrichum salicis CBS607.94; Colletotrichum simmondsii CBS122122; Trichoderma gamsii T6085* [115]*; Colletotrichum orbiculare 104-T* [116]*; Colletotrichum orchidophilum IMI 309357* [117]*; Colletotrichum tofieldiae 0861* [118]*; Conidiobolus coronatus NRRL28638 v1.0; Coemansia reversa NRRL 1564 v1.0; Gonapodya prolifera v1.0* [119]*; Coniochaeta ligniaria NRRL 30616 v1.0* [120]*; Coniochaeta sp. 2T2.1 v1.0* [121]*; Coniophora olivacea MUCL 20566 v1.0* [122]*; Coprinellus micaceus FP101781 v2.0; Coprinopsis marcescibilis CBS121175 v1.0; Crucibulum laeve CBS 166.37 v1.0; Dendrothele bispora CBS 962.96 v1.0; Heliocybe sulcata OMC1185 v1.0; Peniophora sp. CONTA v1.0; Pluteus cervinus NL-1719 v1.0; Polyporus arcularius v1.0; Pterula gracilis CBS309.79 v1.0* [123]*; Coprinopsis cinerea* [124]*; Coprinopsis cinerea AmutBmut pab1-1 v1.0* [125]*; Cordyceps militaris CM01* [126]*; Corynespora cassiicola CCP v1.0* [127]*; Cronartium quercuum f. sp. fusiforme G11 v1.0* [128]*; Cryphonectria parasitica EP155 v2.0* [129]*; Cryptococcus curvatus ATCC 20509 v1.0; Cryptococcus terricola JCM 24523 v1.0* [130]*; Cryptococcus neoformans var neoformans JEC21* [131]*; Cryptococcus neoformans var. grubii H99* [132]*; Cylindrobasidium torrendii FP15055 v1.0; Fistulina hepatica v1.0* [133]*; Cystobasidium minutum MCA 4210 v1.0* [134]*; Daedalea quercina v1.0; Exidia glandulosa v1.0; Fibulorhizoctonia sp*. *CBS 109695 v1.0; Laetiporus sulphureus var. sulphureus v1.0; Neolentinus lepideus v1.0; Peniophora sp. v1.0; Sistotremastrum niveocremeum HHB9708 ss-1 1.0; Sistotremastrum suecicum v1.0; Calocera cornea v1.0; Calocera viscosa v1.0* [135]*; Daldinia eschscholtzii EC12 v1.0; Hypoxylon sp. CI-4A v1.0; Hypoxylon sp. CO27-5 v1.0; Hypoxylon sp. EC38 v3.0* [136]*; Debaryomyces hansenii* [137]*; Dekkera bruxellensis CBS 2499 v2.0* [138]*; Dentipellis sp. KUC8613 v1.0* [139]*; Dichomitus squalens CBS463.89 v1.0; Dichomitus squalens CBS464.89 v1.0; Dichomitus squalens OM18370.1 v1.0* [140]*; Encephalitozoon cuniculi GB-M1* [141]*; Encephalitozoon hellem ATCC 50504; Encephalitozoon romaleae SJ-2008* [142]*; Encephalitozoon intestinalis ATCC 50506* [143]*; Endocarpon pusillum Z07020* [144]*; Endogone sp FLAS 59071; Jimgerdemannia flammicorona AD002; Jimgerdemannia flammicorona GMNB39; Jimgerdemannia lactiflua OSC166217* [145]*; Enterocytozoon bieneusi H348* [146]*; Eremothecium gossypii ATCC 10895* [147]*; Erysiphe necator c* [148]*; Eurotium rubrum v1.0* [149]*; Exophiala dermatitidis UT8656* [150]*; Fibroporia radiculosa TFFH 294* [151]*; Fonsecaea monophora CBS 269.37* [152]*; Fonsecaea nubica CBS 269.64* [153]*; Fusarium fujikuroi IMI 58289* [154]*; Fusarium graminearum v1.0* [155]*; Fusarium oxysporum f. sp. conglutinans race 2 54008 (PHW808); Fusarium oxysporum f. sp. cubense tropical race 4 54006 (II5); Fusarium oxysporum f. sp. lycopersici MN25 (FoMN25) NRRL 54003; Fusarium oxysporum f. sp. radicis-lycopersici 26381 (CL57); Fusarium oxysporum f. sp. raphani 54005; Fusarium oxysporum f. sp. vasinfectum 25433 (Cotton); Fusarium oxysporum Fo47; Fusarium oxysporum NRRL 32931* [156]*; Fusarium oxysporum f. sp. lycopersici 4287 v2; Fusarium verticillioides 7600 v2* [157]*; Fusarium oxysporum f. sp. melonis (FoMelon) NRRL 26406* [158]*; Fusarium oxysporum f. sp. pisi HDV247* [159]*; Fusarium pseudograminearum CS3096* [160]*; Gaeumannomyces graminis var. tritici R3-111a-1; Magnaporthiopsis poae ATCC 64411* [161]*; Glarea lozoyensis ATCC 20868* [162]*; Grosmannia clavigera kw1407* [163]*; Gymnopus androsaceus JB14 v1.0; Chalara longipes BDJ v1.0* [164]*; Heterobasidion annosum v2.0* [165]*; Homolaphlyctis polyrhiza JEL142 v1.0* [166]*; Hortaea werneckii EXF-2000 M0 v1.0* [167]*; Ilyonectria sp. v1.0* [168]*; Kazachstania africana CBS 2517; Torulaspora delbrueckii CBS 1146* [169]*; Kluyveromyces lactis; Yarrowia lipolytica (strain CLIB122)* [170]*; Kuraishia capsulata CBS 1993* [171]*; Laccaria bicolor v2.0* [172]*; Lentinula edodes B17 v1.1* [173]*; Lentinula edodes W1-26 v1.0* [174]*; Lentinus tigrinus ALCF2SS1-6 v1.0; Lentinus tigrinus ALCF2SS1-7 v1.0* [175]*; Leptosphaeria maculans* [176]*; Leucoagaricus gongylophorus Ac12* [177]*; Leucosporidiella creatinivora 62-1032 v1.0; Kockovaella imperatae NRRL Y-17943 v1.0; Naematella encephela UCDFST 68-887.2 v1.0; Clohesyomyces aquaticus v1.0; Pseudomassariella vexata CBS 129021 v1.0; Protomyces lactucaedebilis 12-1054 v1.0; Lobosporangium transversale NRRL 3116 v1.0; Absidia repens NRRL 1336 v1.0; Hesseltinella vesiculosa NRRL3301 v2.0; Rhizopus microsporus var. microsporus ATCC52813 v1.0; Syncephalastrum racemosum NRRL 2496 v1.0; Basidiobolus meristosporus CBS 931.73 v1.0; Linderina pennispora ATCC 12442 v1.0; Catenaria anguillulae PL171 v2.0; Rhizoclosmatium globosum JEL800 v1.0* [178]*; Lichtheimia corymbifera JMRC:FSU:9682* [179]*; Macrophomina phaseolina MS6* [180]*; Magnaporthe oryzae 70-15 v3.0* [181]*; Malassezia globosa* [182]*; Malassezia sympodialis ATCC 42132* [183]*; Marssonina brunnea f. sp. multigermtubi MB_m1* [184]*; Melampsora larici-populina v2.0; Puccinia graminis f. sp. tritici v2.0* [185]*; Melampsora lini CH5* [186]*; Metarhizium acridum CQMa 102;* [187]*; Metarhizium robertsii ARSEF 23;* [188]*; Metschnikowia bicuspidata single-cell v1.0; Dimargaris cristalligena RSA 468 single-cell v1.0; Piptocephalis cylindrospora RSA 2659 single-cell v3.0; Syncephalis pseudoplumigaleata Benny S71-1 single-cell v1.0; Thamnocephalis sphaerospora RSA 1356 single-cell v1.0; Blyttiomyces helicus single-cell v1.0; Caulochytrium protostelioides ATCC 52028 v1.0; Rozella allomycis CSF55 single-cell v1.0* [189]*; Metschnikowia fructicola 277* [190]*; Microbotryum lychnidis-dioicae p1A1 Lamole* [191]*; Microdochium bolleyi J235TASD1 v1.0* [192]*; Microsporum canis CBS 113480; Trichophyton rubrum CBS 118892* [193]*; Mitosporidium daphniae UGP3* [194]*; Mixia osmundae IAM 14324 v1.0; Tilletiaria anomala UBC 951 v1.0* [195]*; Moesziomyces aphidis DSM 70725* [196]*; Monacrosporium haptotylum CBS 200.50* [197]*; Moniliophthora perniciosa FA553* [198]*; Morchella importuna SCYDJ1-A1 v1.0* [199]*; Mortierella elongata AG-77 v2.0* [200]*; Mucor endophyticus; Mucor fuscus; Mucor lanceolatus; Mucor racemosus* [201]*; Mucor lusitanicus (circinelloides) MU402 v1.0* [202]*; Mucor lusitanicus CBS277.49 v2.0; Phycomyces blakesleeanus NRRL1555 v2.0* [203]*; Myceliophthora thermophila (Sporotrichum thermophile) v2.0; Thielavia terrestris v2.0* [204]*; Mycosphaerella graminicola v2.0* [205]*; Nakaseomyces bacillisporus CBS 7720; Nakaseomyces delphensis CBS 2170* [206]*; Nectria haematococca v2.0* [207]*; Nematocida parisii ERTm1* [208]*; Neolecta irregularis DAH-1 v1.0* [209]*; Neonectria ditissima R09/05* [210]*; Neurospora crassa FGSC 73 trp-3 v1.0* [211]*; Neurospora crassa OR74A v2.0* [212]*; Neurospora tetrasperma FGSC 2508 mat A v2.0; Neurospora tetrasperma FGSC 2509 mat a v1.0* [213]*; Nosema ceranae BRL01* [214]*; Obba rivulosa 3A-2 v1.0* [215]*; Omphalotus olearius* [216]*; Ophiostoma novo-ulmi subsp. novo-ulmi H327* [217]*; Ophiostoma piceae UAMH 11346* [218]*; Orpinomyces sp.* [219]*; Paecilomyces niveus CO7 v1.0* [220]*; Paecilomyces variotii CBS 101075 v1.0; Paecilomyces variotii CBS144490 HYG1 v1.0* [221]*; Paracoccidioides brasiliensis Pb03; Paracoccidioides brasiliensis Pb18* [222]*; Paraconiothyrium sporulosum AP3s5-JAC2a v1.0; Pyrenochaeta sp. DS3sAY3a v1.0; Stagonospora sp. SRC1lsM3a v1.0; Alternaria alternata SRC1lrK2f v1.0* [223]*; Penicillium antarcticum IBT 31811; Penicillium coprophilum IBT 31321; Penicillium decumbens IBT 11843; Penicillium flavigenum IBT 14082; Penicillium nalgiovense FM193; Penicillium polonicum IBT 4502; Penicillium solitum IBT 29525; Penicillium steckii IBT 24891; Penicillium vulpinum IBT 29486* [224]*; Penicillium chrysogenum Wisconsin 54-1255* [225]*; Penicillium digitatum Pd1; Penicillium digitatum PHI26* [226]*; Penicillium expansum d1; Penicillium italicum PHI-1* [227]*; Penicillium griseofulvum PG3* [228]*; Penicillium nordicum DAOMC 185683* [229]*; Penicillium oxalicum 114-2* [230]*; Penicillium subrubescens FBCC1632 / CBS132785* [231]*; Penicillium thymicola DAOMC 180753 v1.0* [232]*; Periconia macrospinosa DSE2036 v1.0; Cadophora sp. DSE1049 v1.0* [233]*; Phaeomoniella chlamydospora UCRPC4; Diplodia seriata DS831; Diaporthe ampelina UCDDA912* [234]*; Phanerochaete carnosa HHB-10118-Sp v1.0* [235]*; Phanerochaete chrysosporium RP-78 v2.2* [236]*; Phialocephala scopiformis 5WS22E1 v1.0* [237]*; Phialophora attae CBS 131958* [238]*; Phlebia centrifuga FBCC195* [239]*; Phlebia radiata Fr. (isolate 79, FBCC0043)* [240]*; Phlebiopsis gigantea v1.0* [241]*; Phyllosticta capitalensis CBS 128856 v1.0; Phyllosticta citriasiana CBS 120486 v1.0; Phyllosticta citribraziliensis CBS 100098 v1.0; Phyllosticta citricarpa CBS 127454 v1.0; Phyllosticta citrichinaensis CBS 130529 v1.0; Phyllosticta paracitricarpa CBS 141357 v1.0; Phyllosticta sp. CPC 27913 v1.0* [242]*; Pichia kudriavzevii CBS573* [243]*; Pichia pastoris* [244]*; Piriformospora indica DSM 11827 from MPI* [245]*; Pleurotus ostreatus PC9 v1.0* [246]*; Pneumocystis jirovecii;* [247]*; Pochonia chlamydosporia 170* [248]*; Podospora anserina S mat+* [249]*; Polyporus brumalis BRFM 1820 v1.0* [250]*; Postia placenta MAD 698-R v1.0* [251]*; Postia placenta MAD-698-R-SB12 v1.0* [252]*; Pseudocercospora (Mycosphaerella) fijiensis v2.0* [253]*; Pseudogymnoascus destructans 20631-21* [254]*; Pseudozyma antarctica T-34* [255]*; Pseudozyma hubeiensis SY62* [256]*; Psilocybe cubensis v1.0; Psilocybe serbica v1.0* [257]*; Puccinia coronata avenae 12NC29; Puccinia coronata avenae 12SD80* [258]*; Puccinia graminis f. sp. tritici 21-0 haplotype A; Puccinia graminis f. sp. tritici 21-0 haplotype B; Puccinia graminis f. sp. tritici Ug99 haplotype A; Puccinia graminis f. sp. tritici Ug99 haplotype C* [259]*; Puccinia striiformis f. sp. tritici 104 E137 A-* [260]*; Puccinia striiformis f. sp. tritici PST-130* [261]*; Puccinia striiformis f. sp. tritici PST-78 v1.0; Puccinia triticina 1-1 BBBD Race 1* [262]*; Pycnoporus cinnabarinus BRFM 137* [263]*; Pycnoporus coccineus BRFM 310 v1.0; Pycnoporus puniceus CIRM-BRFM 1868 v1.0; Pycnoporus sanguineus BRFM 1264 v1.0; Ramaria rubella (R. acris) UT-36052-T v1.0* [264]*; Pyrenophora teres f. teres* [265]*; Pyrenophora tritici-repentis* [266]*; Pyronema confluens CBS100304* [267]*; Rhizoctonia solani AG-1 IB* [268]*; Rhizophagus irregularis A1 v1.0; Rhizophagus irregularis A4 v1.0; Rhizophagus irregularis A5 v1.0; Rhizophagus irregularis B3 v1.0; Rhizophagus irregularis C2 v1.0; Rhizophagus irregularis DAOM 197198 v2.0* [269]*; Rhizophagus irregularis DAOM 181602 v1.0* [270]*; Rhizopogon vesiculosus Smith; Rhizopogon vinicolor AM-OR11-026 v1.0* [271]*; Rhizopus delemar 99-880 from Broad* [272]*; Rhizopus microsporus ATCC11559 v1.0; Rhizopus microsporus var. microsporus ATCC52814 v1.0* [273]*; Rhizopus microsporus var. chinensis CCTCC M201021* [274]*; Rhodosporidium toruloides IFO0559_1; Rhodosporidium toruloides IFO0880 v2.0; Rhodosporidium toruloides IFO1236_1* [275]*; Rhodosporidium toruloides IFO0880 v4.0* [276]*; Rhodosporidium toruloides NP11* [277]*; Rhodotorula graminis strain WP1 v1.1* [278]*; Rhodotorula sp. JG-1b* [279]*; Rickenella fibula HBK330-10 v1.0* [280]*; Rickenella mellea v1.0 (SZMC22713;* [281]*; Rigidoporus microporus ED310 v1.0* [282]*; Rozella allomycis CSF55* [283]*; Saccharomyces arboricola H-6* [284]*; Saccharomyces cerevisiae M3707 Dikaryon; Saccharomyces cerevisiae M3836 v1.0; Saccharomyces cerevisiae M3837 v1.0; Saccharomyces cerevisiae M3838 v1.0; Saccharomyces cerevisiae M3839 v1.0; Arthroderma benhamiae CBS 112371; Trichophyton verrucosum HKI 0517* [285]*; Saccharomyces cerevisiae S288C* [286]*; Saksenaea vasiformis B4078* [287]*; Scheffersomyces stipitis NRRL Y-11545 v2.0* [288]*; Schizophyllum commune H4-8 v3.0* [289]*; Schizopora paradoxa KUC8140 v1.0* [290]*; Schizosaccharomyces cryophilus OY26; Schizosaccharomyces japonicus yFS275; Schizosaccharomyces octosporus yFS286* [291]*; Schizosaccharomyces pombe* [292]*; Sclerotinia sclerotiorum v1.0* [293]*; Serpula himantioides (S.lacrymans var shastensis) MUCL38935 v1.0* [294]*; Serpula lacrymans S7.3 v2.0; Serpula lacrymans S7.9 v2.0* [295]*; Smittium culicis GSMNP; Smittium culicis ID-206-W2; Smittium mucronatum ALG-7-W6; Zancudomyces culisetae COL-18-3* [296]*; Sodiomyces alkalinus v1.0* [297]*; Sphaerosporella brunnea Sb_GMNB300 v2.0* [298]*; Spizellomyces punctatus DAOM BR117* [299]*; Sporisorium reilianum SRZ2* [300]*; Stagonospora nodorum SN15 v2.0* [301]*; Stemphylium lycopersici CIDEFI-216* [302]*; Suillus brevipes Sb2 v2.0* [303]*; Talaromyces borbonicus CBS 141340* [304]*; Talaromyces marneffei ATCC 18224* [305]*; Taphrina deformans* [306]*; Thermomyces lanuginosus SSBP* [307]*; Tolypocladium inflatum NRRL 8044* [308]*; Trametes pubescens FBCC735* [309]*; Trichoderma arundinaceum IBT 40837; Trichoderma brevicompactum IBT40841* [310] *; Trichoderma asperellum CBS 433.97 v1.0; Trichoderma citrinoviride TUCIM 6016 v4.0; Trichoderma guizhouense NJAU 4742; Trichoderma harzianum CBS 226.95 v1.0; Trichoderma longibrachiatum ATCC 18648 v3.0* [311]*; Trichoderma asperellum TR356 v1.0; Trichoderma harzianum TR274 v1.0* [312]*; Trichoderma atrobrunneum ITEM 908* [313]*; Trichoderma atroviride v2.0; Trichoderma virens Gv29-8 v2.0* [314]*; Trichoderma hamatum GD12* [315]*; Trichoderma parareesei CBS 125925* [316]*; Trichoderma pleuroti TPhu1* [317]*; Trichoderma reesei QM6a* [318]*; Trichoderma reesei RUT C-30 v1.0* [319]*; Trichoderma reesei v2.0* [320]*; Trichosporon asahii var. asahii CBS 2479* [321]*; Trichosporon asahii var. asahii CBS 8904* [322]*; Trichosporon oleaginosus IBC0246 v1.0* [323]*; Tuber melanosporum Mel28 v1.2* [324]*; Ustilaginoidea virens* [325]*; Ustilago hordei Uh4857_4* [326]*; Ustilago maydis 521 v2.0* [327]*; Venturia inaequalis; Venturia pirina* [328]*; Verticillium alfalfae VaMs.102; Verticillium dahliae VdLs.17* [329]*; Volvariella volvacea V23* [330]*; Wallemia ichthyophaga EXF-994* [331]*; Wallemia mellicola v1.0* [332]*; Xylona heveae TC161 v1.0* [333]*; Yarrowia lipolytica CLIB89(W29)* [334]*; Yarrowia lipolytica FKP355 v1.0* [335]*; Yarrowia lipolytica PO1f v1.0; Yarrowia lipolytica YlCW001 v1.0* [336]*; Yarrowia lipolytica YB392 v1.0; Yarrowia lipolytica YB419 v1.0; Yarrowia lipolytica YB420 v1.0; Yarrowia lipolytica YB566 v1.0; Yarrowia lipolytica YB567 v1.0* [337]*; Zygosaccharomyces rouxii CBS732* [338]*; Zymoseptoria ardabiliae STIR04_1.1.1; Zymoseptoria pseudotritici STIR04_2.2.1* [339]*; Zymoseptoria brevis Zb18110* [340]*; Melampsora allii-populina 12AY07 v1.0; Melampsora americana R15-033-03 v1.0* (unpublished).

### 2.3. Protein sequence analysis and search for potential subcellular targeting

The protein sequences were aligned using Clustal Omega [341] (https://www.ebi.ac.uk/Tools/msa/clustalo/) and manually checked. The predictions for subcellular localizations were done based on the assessment obtained from the TargetP [342] (http://www.cbs.dtu.dk/services/TargetP/), SignalP [343] (http://www.cbs.dtu.dk/services/SignalP/abstract.php), WolfPsort [344] (https://wolfpsort.hgc.jp/) and ESLpred [345] (http://crdd.osdd.net/raghava/eslpred/) software suites.

### 2.4. Phylogenomic analyses and search for horizontal gene transfers

For each type of Msr, we built a multiple sequence alignment with all the identified sequences using MAFFT version v7.429 [346]. Each alignment was trimmed to remove poorly aligned regions using trimAl 1.2 [347] and manually inspected for the conservation of the catalytic residues. A phylogenetic tree was constructed for each type of Msr using RAxML Master Pthread [348] version 8.2.12 (PROTGAMMAWAG model and 500 bootstraps). The phylogenetic tree was represented using iTOL (https://itol.embl.de/) [349]. To investigate potential horizontal gene transfers, the non-canonical Msr sequences along with the sequences whose phylogeny did not fit the species phylogeny were retrieved. Those sequences were used as BLAST queries against the non-redundant protein sequence (nr) database of NCBI (https://blast.ncbi.nlm.nih.gov/Blast.cgi?PROGRAM=blastp&PAGE_TYPE=BlastSearch&LINK_LOC=blasthome) and the best 100 hits were retrieved. The fungal sequences for which protein accessions outside the fungal kingdom were identified among the highest identity scores were in further used as BLAST queries against nr excluding the taxon Fungi (taxid: 4751) and the 25 hits with the highest identity scores were retrieved. In parallel, the same fungal genes were used as queries against the fungal nr database and the 50 hits with the highest identity scores were retrieved, excluding those with 100% identity with the query. The retrieved fungal hits were manually inspected to exclude incomplete sequences. Non-canonical sequences (lacking canonical active site or of abnormal length) were also removed to avoid potential inconsistencies and long branches in the phylogenetic trees. Finally, for each type of Msr, a phylogenetic tree was constructed that included the candidate fungal gene for horizontal gene transfer, plus 25 non-fungal and 25 fungal sequences with the highest identity scores. For each candidate to HGT, the numbers of exons, the GC content of the coding sequence and the GC content at the third positions of the codons were calculated using the gene structures and coding sequences downloaded from the MycoCosm database. The GC content on the third positions of the codons were calculated only for the coding sequences starting with ATG. All calculations were made using Microsoft® Excel® version 2101.

### 3. Results

### 3.1. MsrA, MsrB and fRMsr are largely conserved across the fungal kingdom

Currently, around 136,000 species of fungi are known and classified into 9 phyla (**Figure 1**) [37, 350]. Together, the Ascomycota and the Basidiomycota form the subkingdom Dikarya, which regroups more than 97 % of all described fungal species (∼84,000 species and ∼48,000 species, respectively) [350]. Each phylum contains three monophyletic subphyla (Pucciniomycotina, Ustilaginomycotina and Agaricomycotina for the Basidiomycota, and Taphrinomycotina, Saccharomycotina and Pezizomycotina for the Ascomycota). The other seven phyla are described as *‘early-diverging fungi’* [350] (**Fig. 1, Data S1**). Using the *S. cerevisiae* MsrA, MsrB and fRMsr protein sequences as queries, we used the BLASTP and TBLASTN software suites to search for *msr* genes in 683 available genomes in the MycoCosm database [50]. The selected genomes were from 595 species that spanned the kingdom Fungi (**Fig. 1, Data S1**), including 65% Ascomycota species, 25% Basidiomycota species and 10% early-diverging fungi. We found that the very great majority of these genomes contained one gene coding for a MsrA and one coding for a MsrB (**Fig. 1, Data S1**), indicating that most fungi have a simple Msr system dedicated to protein oxidation repair, similarly to most other known organisms [24, 26]. Most interestingly, we found that a *fRmsr* gene was present in almost all the analyzed genomes (**Fig. 1, Data S1**). The distribution of *fRmsr* across the fungal kingdom clearly showed that the presence of fRMsr is not limited to bacteria and unicellular eukaryotes as previously described [7–9]. Of note, the search for homologs of the bacterial molybdoenzymes able to reduce MetO gave no significant hits (**data not shown**).

**Figure 1.**
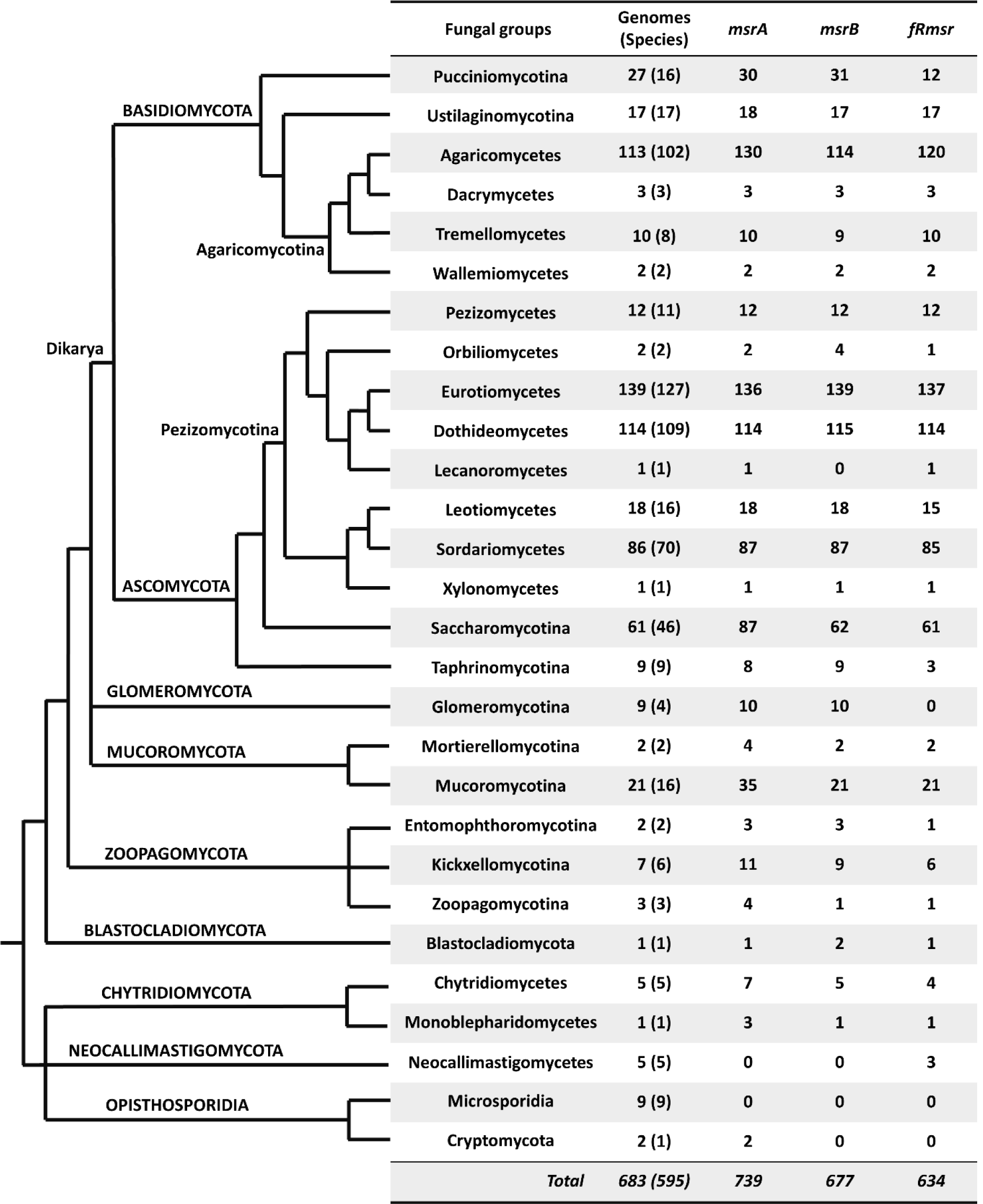
Numbers of *msrA*, *msrB* and *fRmsr* genes identified in fungal genomes. The simplified phylogenetic tree was built according to [37, 350]. The precise number of *msr* genes per genome is available in **Data S1**.

We observed some variations in the numbers of *msr* genes in several genomes. We found 72 genomes, corresponding to 57 species, that had more than one copy of at least one *msr* gene (**Table 1, Data S1-4**). We found 42 genomes having two copies of *msrA* and 17 having three or four copies (**Table 1, Data S1-2**). The highest number of *msrA* copies was 4 in the genome of the Agaricomycetes *Crucibulum laeve* CBS 166.37 and *Dendrothele bispora* CBS 962.96 (**Table 1**). In the case of *msrB*, only 14 genomes had two gene copies (**Data S1, S3**), and the Kickxellomycotina *Smittium culicis* GSMNP was the only genome with three copies (**Table 1**). For *fRmsr*, only six genomes had two copies (**Data S1, S4**), and the Agaricomycetes *Ramaria rubella* (*R. acris*) UT-36052-T and *Coprinellus micaceus* FP101781 had 4 and 3 gene copies, respectively (**Table 1**). Overall, for the three types of Msr, the fungal species having more than one copy were sporadically spread across the fungal kingdom (**Table 1, Data S1-4**), indicating that gene enrichment was not a characteristic of a specific fungal group.

**Table 1.** Fungi with five or six Msr genes.

| Genome | MsrA | MsrB | fRMsr | Lifestyle | Nb. Genes<br>(Genome size in<br>Mbp) | Ref. |
| --- | --- | --- | --- | --- | --- | --- |
| Agaricomycetes |  |  |  |  |  |  |
| *Crucibulum laeve* CBS 166.37 v1.0 | 4 | 1 | 1 | Aerobic (saprotroph) | 14,218 (44) | [351] |
| *Dendrothele bispora* CBS 962.96 v1.0 | 4 | 1 | 1 |  | 33,645 (131) | [351] |
| *Tulasnella calospora* AL13/4D v1.0 | 3 | 1 | 1 |  | 19,659 (62) | [352] |
| *Ramaria rubella* (*R. acris*) UT-36052-T v1.0 | 1 | 1 | 4 |  | 14,207 (62) | [353] |
| *Coprinellus micaceus* FP101781 v2.0 | 1 | 1 | 3 |  | 23,559 (77) | [351] |
| Dothideomycetes |  |  |  |  |  |  |
| *Hortaea werneckii* EXF-2000 M0 v1.0 | 2 | 2 | 2 | Aerobic | 15,748 (50) | [167] |
| Sordariomycetes |  |  |  |  |  |  |
| *Coniochaeta* sp. 2T2.1 v1.0 | 2 | 2 | 2 | Aerobic (phytopathogen) | 24,735 (74) | [121] |
| Saccharomycotina |  |  |  |  |  |  |
| *Pichia kudriavzevii* CBS573 | 3 | 1 | 1 | Aerobic | 5,140 (11) | [354] |
| *Yarrowia lipolytica* (strain CLIB122) | 3 | 1 | 1 |  | 6,447 (21) | [355] |
| *Yarrowia lipolytica* CLIB89(W29) | 3 | 1 | 1 |  | 7,919 (21) | [356] |
| *Yarrowia lipolytica* FKP355 v1.0 | 3 | 1 | 1 |  | 6,858 (20) | [357] |
| *Yarrowia lipolytica* PO1f v1.0 | 3 | 1 | 1 |  | 6,798 (20) | [358] |
| *Yarrowia lipolytica* YB392 v1.0 | 3 | 1 | 1 |  | 6,750 (20) | [359] |
| *Yarrowia lipolytica* YB419 v1.0 | 3 | 1 | 1 |  | 6,751 (20) | [359] |
| *Yarrowia lipolytica* YB420 v1.0 | 3 | 1 | 1 |  | 6,772 (20) | [359] |
| *Yarrowia lipolytica* YB566 v1.0 | 3 | 1 | 1 |  | 6,764 (20) | [359] |
| *Yarrowia lipolytica* YB567 v1.0 | 3 | 1 | 1 |  | 6,776 (20) | [359] |
| *Yarrowia lipolytica* YICW001 v1.0 | 3 | 1 | 1 |  | 6,800 (20) | [358] |
| Mucoromycotina |  |  |  |  |  |  |
| *Absidia repens* NRRL 1336 v1.0 | 3 | 1 | 1 | Aerobic (saprotroph) | 14 919 (47) | [360] |
| *Rhizopus microsporus* ATCC11559 v1.0 | 2 | 1 | 2 | Aerobic (phytopathogen) | 11,135 (24) | [273] |
| *Rhizopus microsporus* var. *chinensis* CCTCC M201021 | 2 | 1 | 2 |  | 17,676 (46) | [274] |
| Kickxellomycotina |  |  |  |  |  |  |
| *Smittium mucronatum* ALG-7-W6 | 3 | 1 | 1 | Anaerobic (insect gut) | 8 247 (102) | [361] |
| *Smittium culicis* GSMNP | 2 | 3 | 1 |  | 12 166 (77) | [361] |
| Monoblepharidomycetes |  |  |  |  |  |  |
| *Gonapodya prolifera* v1.0 | 3 | 1 | 1 | Aerobic (aquatic) | 13 902 (49) | [362] |

Also, we found only 74 genomes (58 species), in which one or more Msr types were absent (**Table 2**). Among them, the nine Microsporidia species, two Neocallimastigomycetes species (out of five species analyzed) and one Taphrinomycotina species, *Pneumocystis jirovecii* (out of eight species analyzed) lacked all three Msr types. These species are either obligate intracellular parasites (Microsporidia and *Pneumocystis jirovecii*) or live in the anaerobic conditions of the animal gut (*Piromyces finnis* and *Piromyces* sp. E2) [37, 363]. Interestingly, the three other Neocallimastigomycetes species (*Anaeromyces robustus, Neocallimastix californiae* G1 and *Orpinomyces* sp.), also living in anaerobic conditions [350], do not possess MsrA nor MsrB but have a gene coding for a fRMsr (**Table 2**). Our results show that, additionally to *Encephalitozoon cuniculi,* previously identified as the unique eukaryote lacking both MsrA and MsrB [24], 14 other fungal species are devoid of any protein-bound MetO reduction enzyme (**Table 2**). Furthermore, we found six species lacking *msrA* but having a *msrB* and a *fRmsr* gene (**Table 2**). These are the agaricomycete *Moniliophthora perniciosa*, the eurotiomycetes *Cladophialophora immunda*, *Penicillium coprophilum* and *Penicillium flavigenum*, the sordiariomycete *Magnaporthiopsis poae* and the saccharomycotina *Saturnispora dispora*. To our knowledge, they constitute the first species, over all kingdoms, described to have only a MsrB to reduce and repair oxidized proteins, as none was found so far in genome surveys [24, 26]. Of note, we did not find fungal species having only a MsrB, as these six species also possessed a fRMsr (**Table 2**). We found that the five species *Trichosporon asahii* (Tremellomycetes), *Clohesyomyces aquaticus* (Dothideomycetes), *Cladonia grayi* (Lecaronomycetes) and *Piptocephalis cylindrospora* (Zoopagomycotina) possessed a MsrA and a fRMsr but lacked a MsrB and that the Zoopagomycotina *Syncephalis pseudoplumigaleata* had a MsrA only (**Table 2**). Finally, we found only 27 species having both a MsrA and a MsrB but lacking a fRMsr (**Table 2**). These species were sporadically dispersed among the fungal kingdom, but three groups stood out as remarkable: the Glomeromycotina, for which none of the four species analyzed had a fRMsr, the Taphrinomycotina for which six species out of eight analyzed (including the four *Schizosaccharomyces* species analyzed) were devoid of fRMsr, and the Pucciniomycotina for which all the *Melampsora* and *Puccinia* species analyzed here lacked a fRMsr (**Table 2**).

**Table 2.**
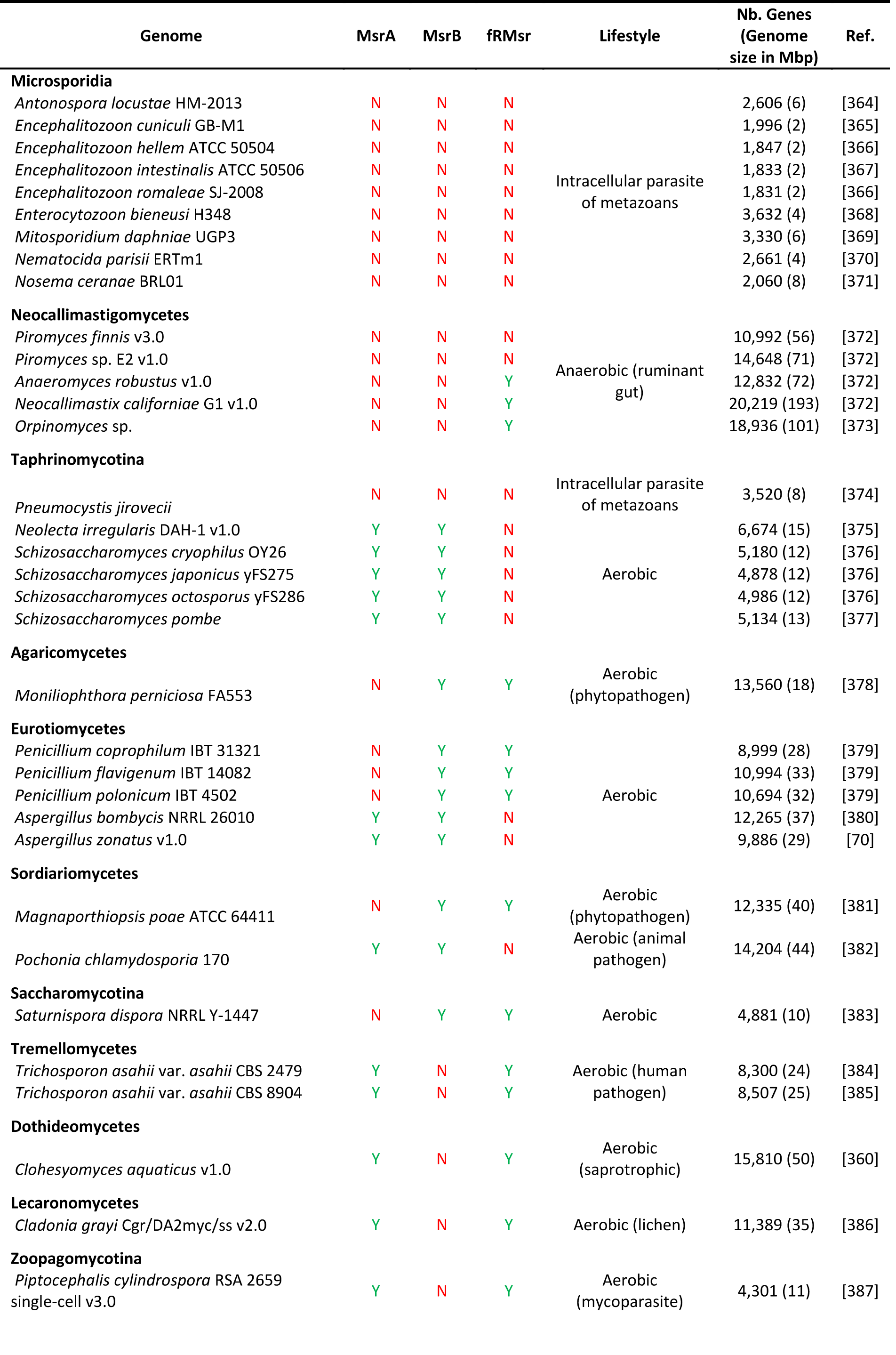
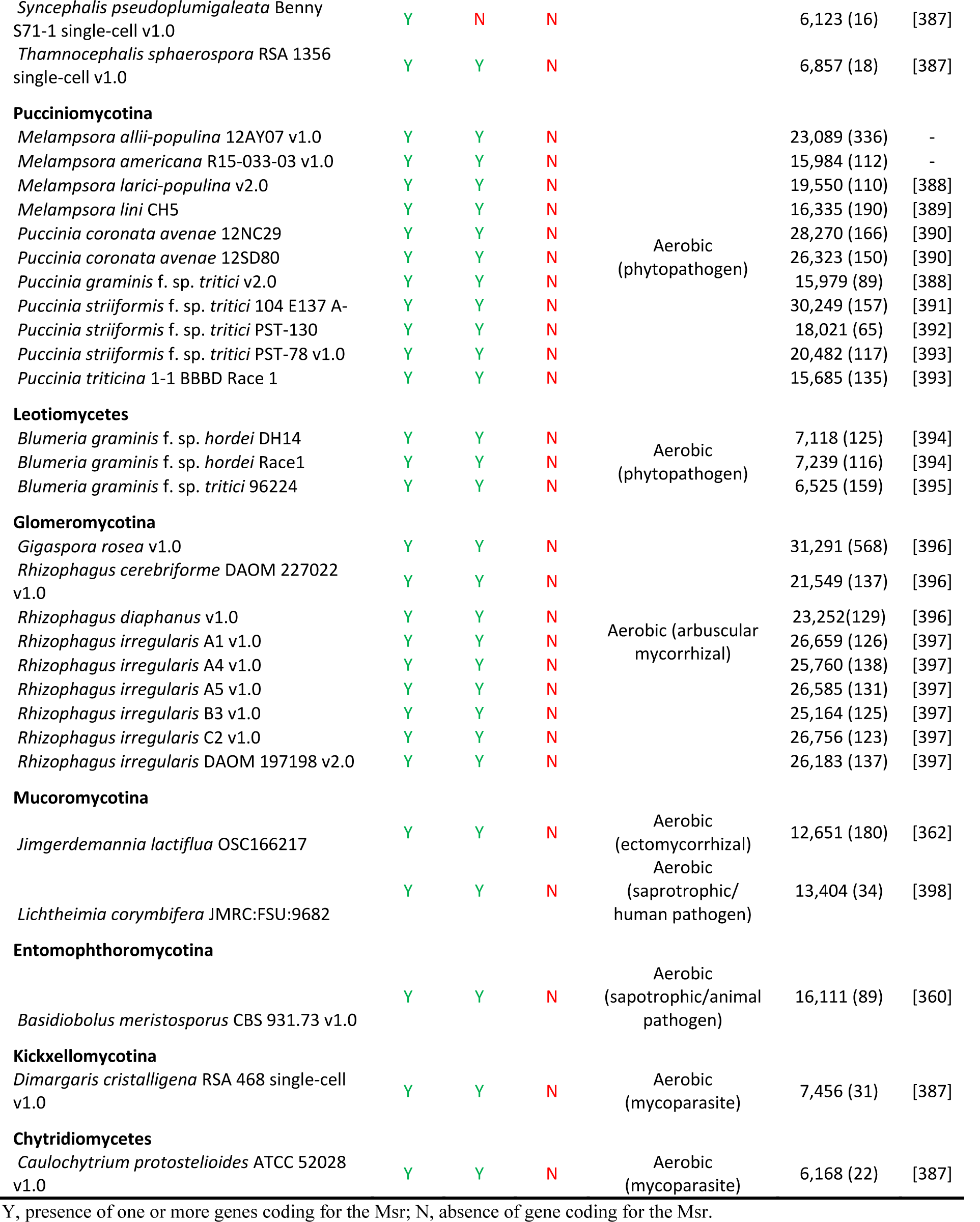
Fungal genomes lacking one or more Msr types.

Altogether, these results showed that the great majority of fungi possess one gene coding for each protein-repairing Msr type (i.e. MsrA and MsrB), as well as one gene coding for the free MetO reductase fRMsr.

### 3.2. Fungal Msrs globally retained canonical sequence features for activity

Sequence analysis and enzymatic characterizations of MsrAs from several organisms showed that the catalytic residue is located within a G[C/U]FW motif, located in the N-terminal part of the protein [6,16,17,399,400]. The great majority of known MsrAs possess a catalytic Cys, whereas a few Sec-containing proteins are found in some insects, marine organisms and unicellular algae [25, 400]. In canonical MsrAs, the resolving Cys, involved in the regeneration of the catalytic Cys, is located in the C-terminal part of the protein, althought the number and positions of resolving Cys vary [6,17,22,23]. To determine whether the fungal MsrAs share these properties, we analyzed the 709 full length MsrAs identified. These MsrAs ranged from 142 to 322 amino acids, and all of them were made of a single MsrA domain (**Data S2A**). The length differences were mostly due to the presence of N-terminal extensions of variable sizes, indicating the possible presence of signal peptides for protein distribution in subcellular compartments. We used several targeting prediction programs (see section 2.3) to evaluate the potential subcellular localization of the fungal MsrAs (**Data S2A**). Most MsrAs were predicted to be localized into the cytoplasm (45 %) or in the mitochondria (44 %) (**Fig. 2A, Data S2A**). The other sequences were predicted to be secreted (6%), to be localized in other compartments, or had no clearly assigned localization (6%) (**Fig. 2A, Data S2A**). The alignment of the primary sequences revealed that the great majority (∼ 90 %) shared common features with the previously characterized *S. cerevisiae* MsrA [6, 401] (**Fig. 3A, Data S2B**). The catalytic Cys (position 25 in the *S. cerevisiae* MsrA) was located in the conserved motif ^24^GCFW^27^. The Tyr^64^, Glu^76^, Asp^111^ and Tyr^116^ residues involved in substrate stabilization and catalysis [399], and the residues Gly^47^, His^100^, Gln^108^, Gly^113^, His^163^ and Tyr^166^ were also conserved. Finally, the Cys^176^, previously identified as resolving Cys [6, 401], was included in the ^173^GYXC^176^ motif (**Fig. 3A, Data S2B**). Because of their predominance in all the fungal kingdom, we defined the fungal MsrAs having these properties as *‘canonical’* sequences. The fungal MsrAs that did not match these sequence features were defined as *‘non-canonical’* MsrAs (**Data S2A, C, D**). Particularly, we observed that in ∼ 5 % of the identified sequences, the Phe^26^ residue in the ^24^GCFW^27^ motif containing the catalytic Cys, was substituted by a Tyr. We also observed the replacement of Asp^111^ by an Asn residue in few sequences (∼ 2 %) (**Data S2C**). Moreover, some variations were also observed for the resolving Cys (**Data S2C**). Fourteen sequences (∼ 2 %) lacked the conserved ^173^GYXC^176^ motif but possessed two to four Cys in a Q[C/S/K]X2KX[C/N][C/X]XI[R/L]CYG motif, similar to poplar MsrAs [23]. Some other sequences possessed a Cys residue in the C-terminal region, but not in a GYXC motif, and others lacked any potential resolving Cys (**Data S2C**). Finally, a special case could be made for MsrAs from the early-diverging fungus *Gonapodya prolifera* (Monoblepharidomycetes). This fungus has three non-canonical MsrAs, two of which had the catalytic Cys replaced by a Sec [402]. Each had another Cys outside the conserved position of the resolving Cys in canonical fungal MsrAs. These two MsrA sequences had high similarity with the Sec-MsrAs from the bacterium *Alkaliphilus oremlandii* and the single-cell green alga *Chlamydomonas reinhardtii*, previously shown to use the Sec residue for the regeneration of their activity [400, 403] (**Data S2D**).

**Figure 2.**
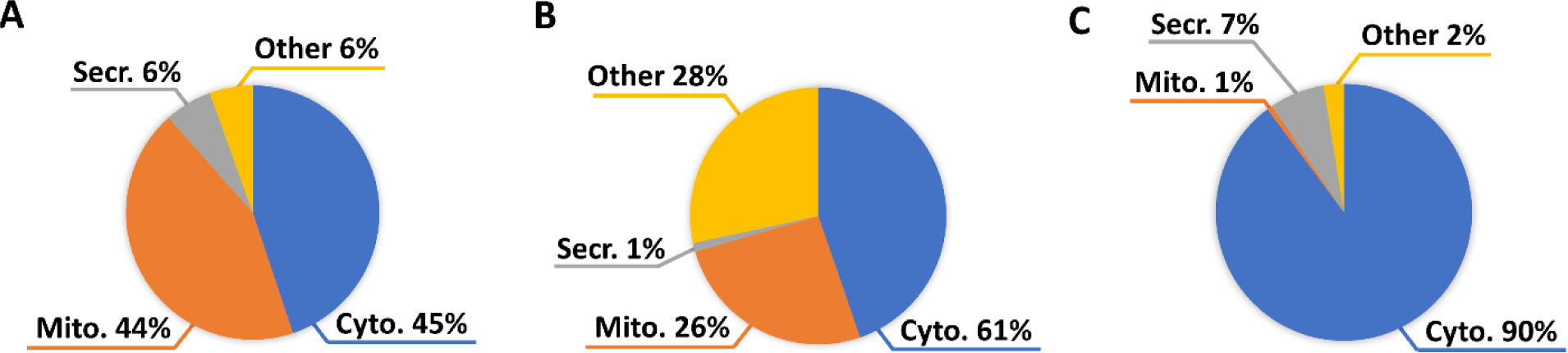
Potential subcellular targeting of fungal Msrs. The circle charts show the subcellular predictions for fungal MsrA (**A**), MsrB (**B**) and fRMsr (**C**), the proportion of proteins predicted to be localized in the cytosol (‘*Cyto.’*), to be addressed to the mitochondria (‘*Mito.’*) or secreted (‘*Secr.’*). The label *‘Other’* indicates the proportion of proteins predicted to be addressed to other compartments or for which no consensus prediction was obtained.

**Figure 3.**
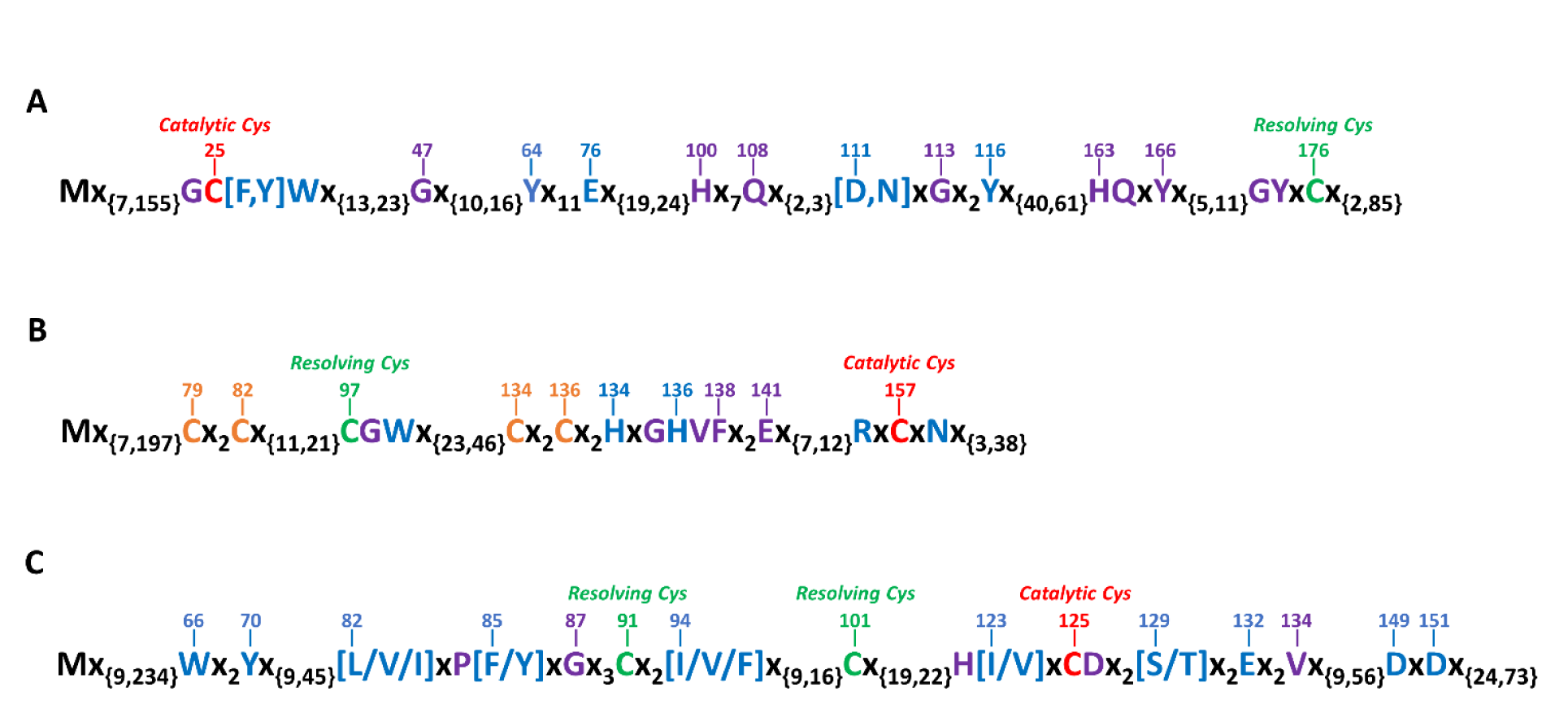
Protein sequence characteristics of canonical fungal Msrs. In this representation of canonical MsrAs (**A**), MsrBs, (**B**) and fRMsrs (**C**), the catalytic Cys (in *red*), the resolving Cys (in *green*) and the residues previously shown to be involved in catalysis and/or substrate binding (in *blue*) are shown. The residues in *purple* are conserved in all canonical fungal Msrs. In **B**, the Cys residues labeled in *orange* correspond to Zn binding residues. The numberings are based on *S. cerevisiae* MsrA (**A**), MsrB (**B**) and fRMsr (**C**).

In the case of MsrBs, previous biochemical characterizations demonstrated that the catalytic Cys is located in a RXCXN motif in the C-terminal part of the protein [16,21,404–407]. Mammals express a Sec-containing form in which the Asp is replaced by a Phe [404]. The resolving Cys is generally located in a CGWP motif present in the N-terminal part of the protein [21, 406]. In the case of mammalian MsrBs, one or two resolving Cys, located in the N-terminal extremity of the protein, can be involved in the regeneration process [404, 408]. Of note, most MsrBs possess two CX2C clusters coordinating a structural Zn atom [409]. Here, we analyzed the 651 complete fungal MsrB protein sequences. The fungal MsrBs consisted of a single domain and ranged in length from 95 to 289 amino acids (**Data S3A**). Most of the variations in size were due to the presence of an N-terminal extension potentially involved in subcellular targeting. The majority of fungal MsrBs were predicted to be addressed to the cytosol (61 %) (**Fig. 2B, Data S3A**). The other proteins were either predicted to be localized in the mitochondria (26 %), secreted (1 %), targeted to another compartment, or were not clearly predicted to be addressed to a subcellular compartment (28 %) (**Fig. 2B, Data S3A**). Almost all fungal MsrBs (> 99 %) possessed the features of the *S. cerevisiae* enzyme (**Fig. 3B, Data S3B**): i) the two CX2C motifs involved in the coordination of a Zn atom, ii) the resolving Cys^97^ (according to *S. cerevisiae* MsrB residue numbering) included in a ^97^CGW^99^ motif, iii) the conserved His^134^ and His^136^ implicated in substrate binding, together with Arg^155^ and Asn^159^ [16, 405], and iv) the catalytic Cys^157^ located in the ^155^RXCXN^159^ motif. The Gly^98^, Gly^135^, Val^137^, Phe^138^ and Glu^141^ residues were also conserved in all these canonical fungal MsrBs (**Fig. 3B, Data S3B**). Only four sequences (< 1 %), from the orbiliomycetes *Arthrobotrys oligospora* and *Monacrosporium haptotylum*, the taphrinomycotina *Protomyces lactucaedebilis* and the chytridiomycete *Blyttiomyces helices*, presented remarkable differences in primary sequence features (**Data S3C**). These MsrBs lacked the resolving Cys at position 97, which was substituted by a Ser or a Thr, like plant and human orthologs that use an unusual regeneration process [20,21,408]. Finally, another unusual feature was found in the MsrB from the taphrinomycotina *Protomyces lactucaedebilis*, with the location of the catalytic Cys in an HYCIN motif, instead of the typical RXCXN motif (**Data S3C**). Interestingly, searching in the NCBI nr database, we found 50 sequences, mainly from poorly characterized bacteria and archaea, that possessed this unusual motif (**Data S3D**). Considering His and Arg have similar physicochemical properties, we anticipate such non-canonical MsrBs might have conserved a catalytic activity.

Very few fRMsrs have been characterized so far. However, sequence comparison studies and biochemical characterizations indicated that the catalytic Cys is located in the HIAC motif situated in the middle of the protein sequence and that the two resolving Cys are located ∼30 and ∼40 amino acids upstream in the N-terminal direction [7–9]. The 589 full length fRMsr sequences analyzed here had a single fRMsr domain. Their length varied from 77 to 394 amino acids, with variations in the size of the N-terminal extension (**Data S4A**). Most of the proteins (90 %) were predicted to be localized in the cytoplasm (**Fig. 2C, Data S4A**). A few proteins were predicted to be secreted (7 %), targeted to the mitochondria (1 %), to other compartments or had no reliable prediction for subcellular targeting (2 %) (**Fig. 2C, Data S4A**). Similar to MsrAs and MsrBs, fRMsrs showed a strong conservation of the sequence features. Almost all (> 99 %) sequences possessed the catalytic Cys^125^ included in a ^122^H[I/V]XCD^126^ motif and the resolving Cys in positions 91 and 101 (according to the *S. cerevisiae* fRMsr residue numbering) (**Fig. 3C, Data S4B**). We also observed the strict conservation of Trp^66^, Tyr^70^, Glu^132^, Asp^149^ and Asp^151^, previously shown to be involved in substrate binding and catalysis. Other important residues involved in substrate binding and catalysis [7] were also conserved or substituted by residues with similar properties in positions 82, 85, 94, 123, 129 and 132. The Pro^84^, Gly^87^, His^122^ and Val^134^ were also strictly conserved in these canonical fungal fRMsrs (**Fig. 3C, Data S4B**). Only three fRMsr sequences from the Agaricomycetes *Scleroderma citrinum*, *Dendrothele bispora* and *Pisolithus tinctorius* presented non-canonical characteristics. The first two lacked the potential resolving Cys^101^ and may be still able to reduce the free MetO, but in the latest, the catalytic Cys^125^ was substituted by an Arg, likely precluding catalytic activity (**Data S4C**).

Altogether, these results uncover few proteins with non-canonical sequence features, but principally showed that for each Msr type, the residues involved in catalysis are globally conserved throughout the fungal kingdom.

### 3.3. The phylogenetic analysis of fungal Msrs revealed horizontal gene transfers from bacteria

The phylogenetic relationship of fungal MsrAs globally matched the expected clustering for early-diverging fungi, Ascomycota and Basidiomycota sequences (**Fig. S1-3**). However, a few Ascomycota MsrA sequences clustered with Basidiomycota sequences (indicated by an asterisk on **Fig. S1**). We also noticed the clustering of all the MsrB sequences from Pucciniomycotina (Basidiomycetes) species with Ascomycota sequences (**Fig. S2**). Strikingly, the MsrB sequences from early-diverging fungi did not group in a single cluster but were interspersed in clusters containing Basidiomycota or Ascomycota sequences (**Fig. S2**). For fRMsrs, we observed three distinct clusters containing the protein sequences from Basidiomycota, Ascomycota and early-diverging fungi, respectively (**Fig. S3**). However, two sequences from early-diverging fungi were included in the cluster containing the Ascomycota sequences (**Fig. S3**). Altogether, these results showed that the phylogeny of Msrs was globally congruent with the phylogeny of the species, except for a few protein sequences.

Several discrepancies suggested that some Msrs could have arisen from horizontal gene transfer. These discrepancies were: i) the positioning in phylogenetic clusters not reflecting the phylogeny of the species from which they were isolated (**Fig. S1-3**), ii) the presence of non-canonical sequence features (**Data S2-4**), and iii) the presence of *fRmsr* genes in the genome of three strictly anaerobic neocallimastigomycetes, whereas other organisms from the same phylum had no *msr* genes (**Table 2**).

To evaluate the possibility of horizontal gene transfers, we selected all Msr sequences (67 MsrAs, 48 MsrBs and 8 fRMsrs) with one or more of these discrepancies and searched for their closest putative homologs by BLASTP search in all organisms recorded in the NCBI nr database. We discarded all the sequences for which the closest homologs were found in other fungal taxa. Indeed, because of the strong conservation of the protein sequences, it would be difficult to ascertain the occurrence of fungus-to-fungus horizontal gene transfers. We retrieved 17 MsrA, three MsrB and three fRMsr protein sequences for further analysis (**Fig. 4-6**). For each type of Msr, we used the selected protein sequences, together with the respective 25 sequences with the highest identity score from fungi on the one hand and from non-fungal organisms on the other hand for phylogenetic analyses (**Fig. 4-6**). The phylogenetic trees highlighted two MsrA, three MsrB and three fRMsr sequences, among the selected candidates, clustering with bacterial and amoeba homologs (**Fig. 4-6, Table 3**). Moreover, we identified two additional fungal fRMsrs, not included in our primary data set, which clustered with bacterial proteins (**Fig. 6, Table 3**). Except for two MsrBs from Orbiliomycetes (Ascomycota), all these Msr sequences were from early-diverging fungi (**Table 3**). Interestingly, in the cases of MsrA and MsrB, they were all from organisms having another gene coding for a canonical enzyme in their genomes (**Data S1-3**).

**Figure 4.**
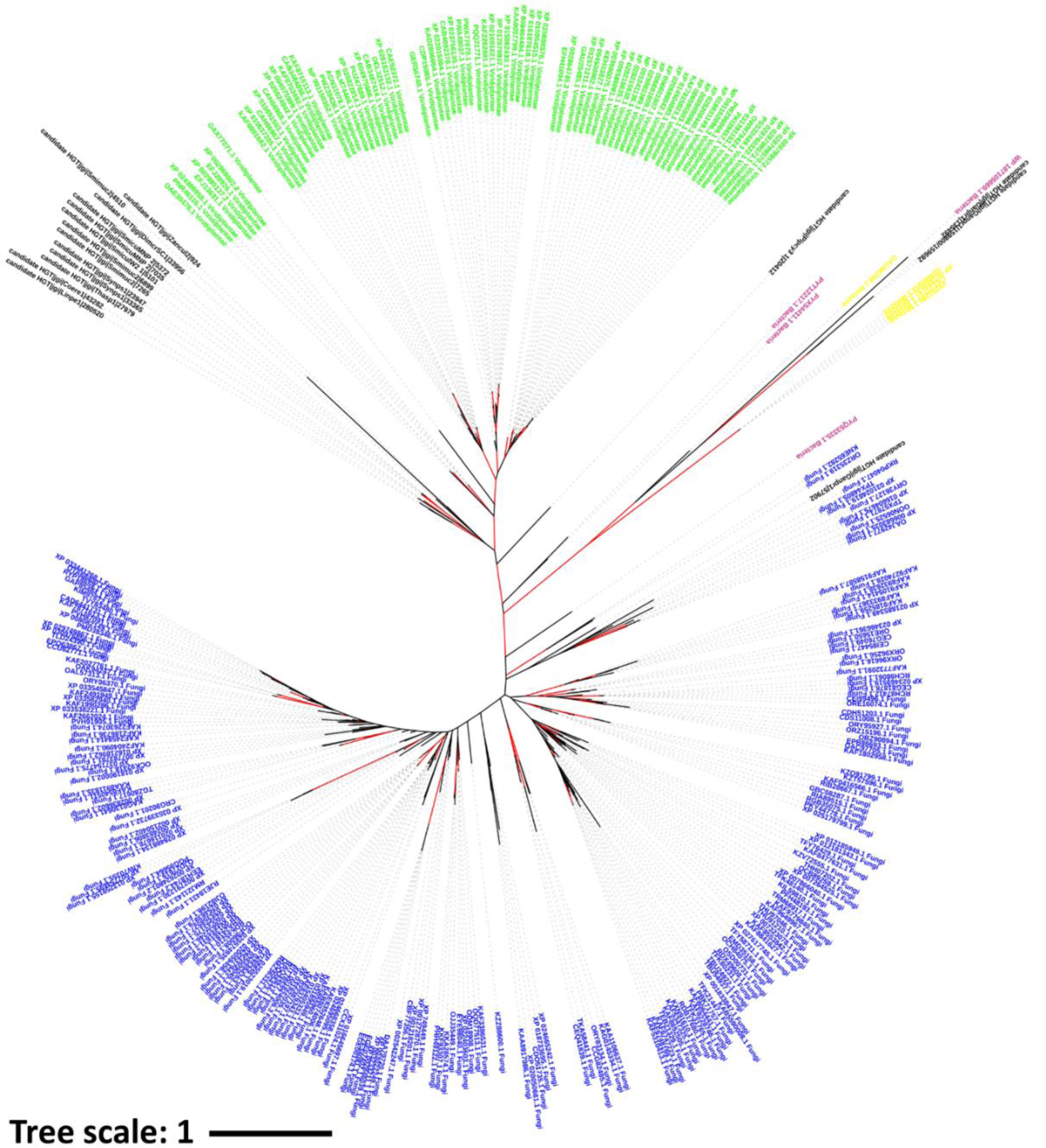
Phylogenetic analysis of MsrA candidates to horizontal gene transfer. The fungal MsrAs tested for horizontal gene transfer (in *black*) are from the following fungal genomes (protein accessions in brackets): *Piptocephalis cylindrospora* RSA 2659 single-cell v3.0 (20412); *Syncephalis pseudoplumigaleata* Benny S71-1 single-cell v1.0 (33365; 23947); *Thamnocephalis sphaerospora* RSA 1356 single-cell v1.0 (27979); *Coemansia reversa* NRRL 1564 v1.0 (43282); *Dimargaris cristalligena* RSA 468 single-cell v1.0 (33956); *Linderina pennispora* ATCC 12442 v1.0 (280520); *Smittium culicis* GSMNP (5372; 7035); *Smittium culicis* ID-206-W2 (5101; 8129); *Smittium mucronatum* ALG-7-W6 (4510; 6899; 7265); *Zancudomyces culisetae* COL-18-3 (924) and *Gonapodya prolifera* v1.0 (135492; 159800/159692; 57902). The MsrA sequences from fungi, plants, amoeba and bacteria are in *blue*, *green*, *yellow* and *purple*, respectively. The phylogenetic tree was built with RAxML v. 8.2 [348] and represented using iTOL (https://itol.embl.de/) [412]. The branches with bootstrap values over 70 are in *red*.

**Figure 5.**
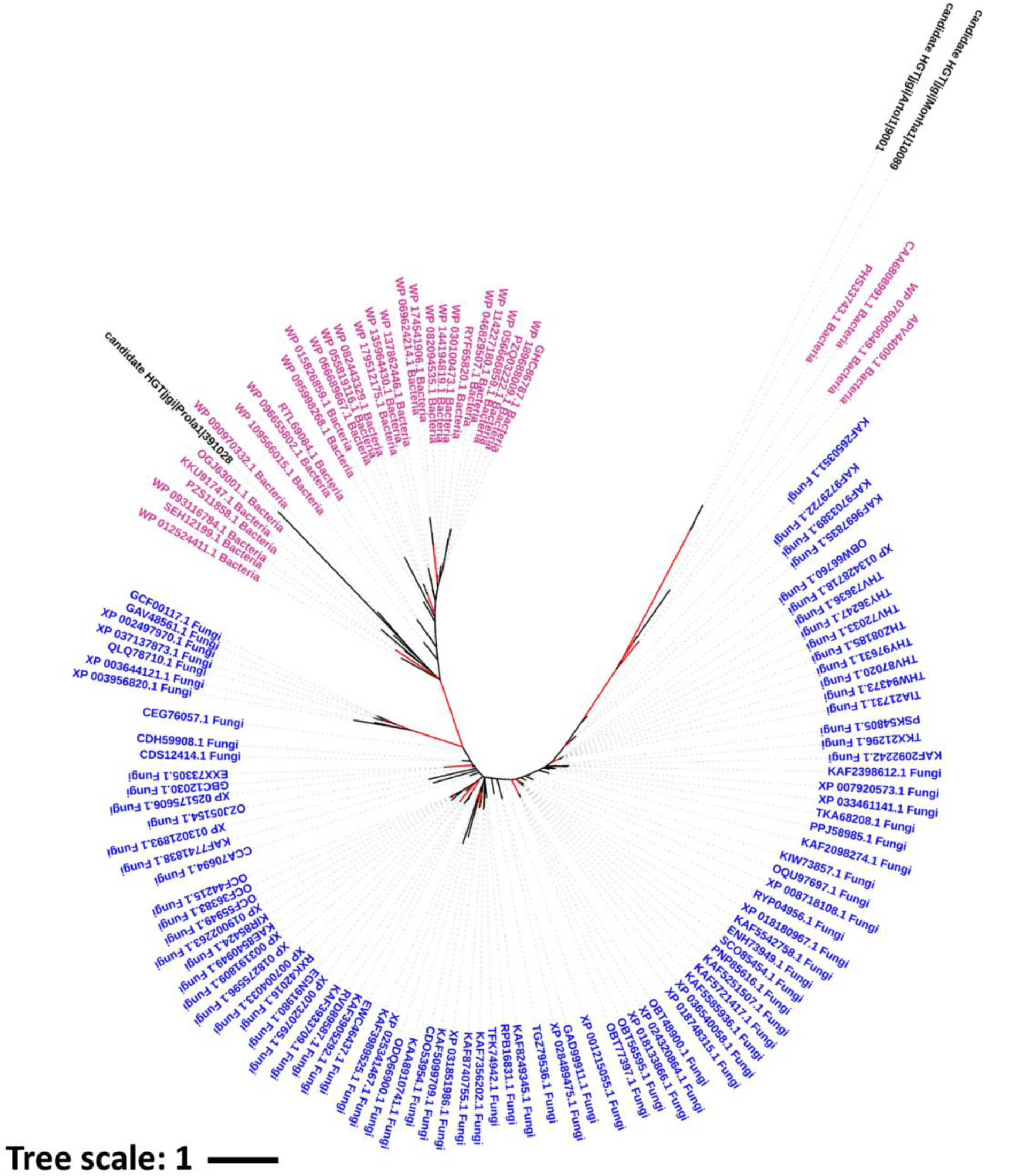
Phylogenetic analysis of MsrB candidates to horizontal gene transfer. The fungal MsrBs tested for horizontal gene transfer (in *black*) are from the following genomes (protein accessions in brackets): *Arthrobotrys oligospora* ATCC 24927 (9001); *Monacrosporium haptotylum* CBS 200.50 (10089) and *Protomyces lactucaedebilis* 12-1054 v1.0 (391028). The MsrBs sequences from fungi and bacteria are in *blue* and *purple*, respectively. The phylogenetic tree was built with RAxML v. 8.2 [348] and represented using iTOL (https://itol.embl.de/) [412]. The branches with bootstrap values over 70 are in *red*.

**Figure 6.**
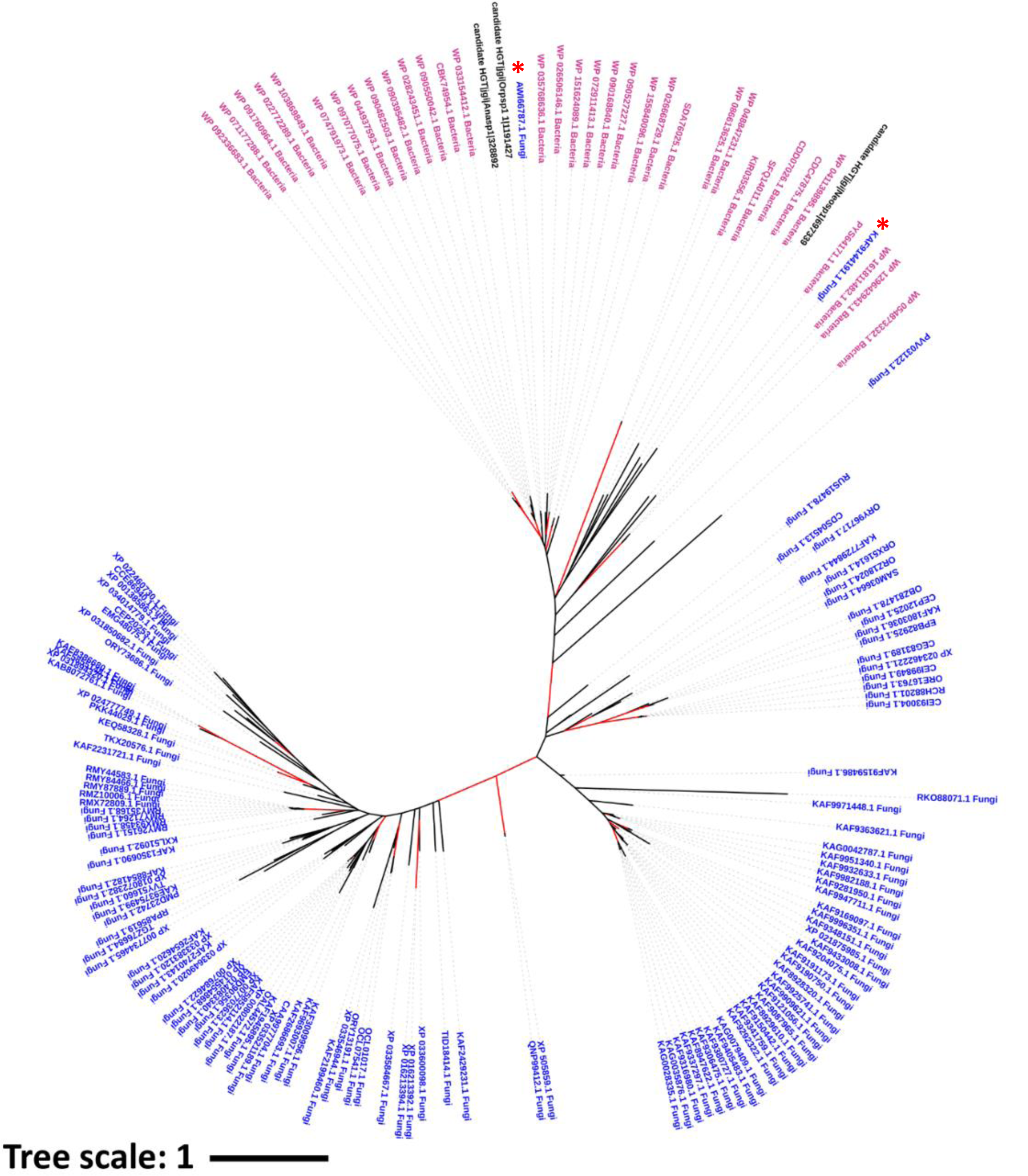
Phylogenetic analysis of fRMsr candidates to horizontal gene transfer. The fungal fRMsrs tested for horizontal gene transfer (in *black*) are from the following genomes (protein accessions are in brackets): *Anaeromyces robustus* v1.0 (328892); *Neocallimastix californiae* G1 v1.0 (697339) and *Orpinomyces sp* (1191427). The fRMsrs sequences from fungi and bacteria are in *blue* and *purple*, respectively. *Red* stars indicate *fRmsr* genes possibly acquired by horizontal gene transfer, which were not identified in our genomics search but found as homologs of the selected fungal fRMsr candidates. The phylogenetic tree was built with RAxML v 8.2 [348] and represented using iTOL (https://itol.embl.de/) [412]. The branches with bootstrap values over 70 are in *red*.

**Table 3.**
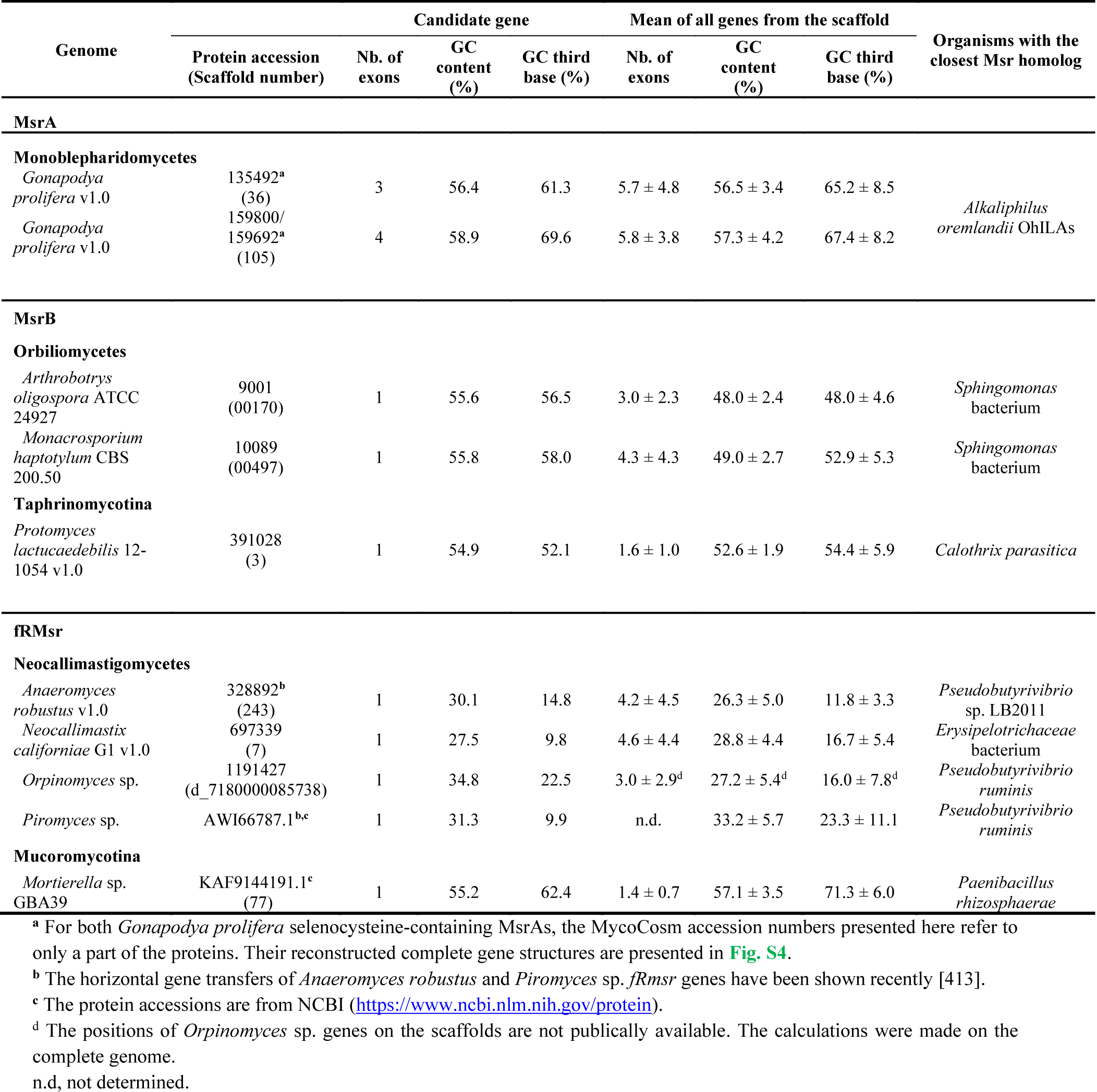
Potential horizontal *msr* gene transfers from bacteria to fungi

For MsrAs, a potential horizontal gene transfer was observed for the two selenocysteine-containing enzymes from *Gonapodya prolifera* (**Table 3**). Very interestingly, their closest homolog was also a selenocysteine-containing MsrA, from the bacteria *Alkaliphilus oremlandii*. Bacteria from the genus *Alkaliphilus* are found in sediments and ponds [410], and *Gonapodya prolifera* occurs on fruits submerged into ponds [411]. The presence of both organisms in a same ecological niche could have favored the horizontal gene transfer. The similarity between the two *Gonapodya prolifera* Sec-MsrA suggested they arose from one horizontal gene transfer event followed by a gene duplication (**Fig. 4, Fig. S4**).

In the case of MsrBs, the three selected sequences clustered with bacterial MsrBs (Fig. 5). The two MsrBs from the Orbiliomycetes species *Arthrobotrys oligospora* and *Monacrosporium haptotylum* had MsrBs from *Sphingomonas* bacteria as closest homologs. Both fungi trap nematodes in soil [197], and sphingomonads have been isolated from many different land and water habitats [414], indicating that the co-occurrence in the same habitat of a sphingomonad donor and a common ancestor of *A. oligospora* and *M. haptotylum* is plausible, which could have allowed horizontal gene transfers. Another potential horizontal gene transfer for MsrB was identified, from a donor cyanobacterium related to the marine *Calothrix parasitica* to the plant pathogen fission yeast *Protomyces lactucaedebilis* (Taphrinomycotina) (**Table 3**). In this case, the ecology of the extant candidate donor and acceptor do not support the co-occurrence of both organisms in a same ecological niche.

The three fRMsrs we selected as candidates to horizontal gene transfer were from the neocallimastigomycetes species *Anaeromyces robustus, Neocallimastix californiae* and *Orpinomyces sp.* (**Table 3**). In our phylogenetic analysis, they clustered with sequences of bacteria from the phylum Firmicutes which, similarly to neocallimastigomycetes, live in the gut of ruminants (**Fig. 6**). Interestingly, two other fungal fRMsr sequences, one from *Piromyces* sp. and one from *Mortierella* sp. GBA39, were also present in the same cluster. These two sequences were omitted from our genomic search because of the absence of the corresponding genomes in the MycoCosm database. Of note, the horizontal transfer of *fRmsr* genes from Firmicutes to neocallimastigomycetes has been shown recently for *Anaeromyces robustus* and *Piromyces* sp. [413]. Altogether, these results strongly argue for the fact that the presence of *fRmsr* in the genomes of these anaerobic fungi arose from horizontal gene transfers from Firmicutes (**Table 3**).

In addition to the phylogenetic method, we performed a parametric analysis of the numbers of exons, the overall GC contents of their coding sequence (CDS), and the GC contents of the third position of each codon to identify potential bias as supporting arguments of horizontal gene transfer (**Table 3**). The comparison of these parameters with those of the other genes included in the same genomic scaffold were shown as potential indicators of horizontal gene transfers [415]. Excepted for the two *Gonapodya prolifera msrA* genes, which had no distinguishable values from the other genes included in their scaffolds, all other genes coding MsrBs or fRMsrs, had extreme values for at least two of the three parameters considered (**Table 3**). First of all, all these eight genes were made of a single exon, like bacterial genes. The global GC content of *Arthrobotrys oligospora* and *Monacrosporium haptotylum msrB* genes coding sequences were the highest of all those included in their scaffold. Similarly, the global GC content of *Protomyces lactucaedebilis msrB*, *Anaeromyces robustus fRmsr* and *Orpinomyces* sp. *fRmsr* were among the highest (**Table 3**). In the cases of *fRmsrs* from *Neocallimastix californiae, Piromyces* sp. and *Mortierella* sp., the GC contents of the third base of the codons were among the lowest by comparison with the other genes present in the scaffold. Noticeably, we also observed a lower GC content at the third position of the codons in the *fRmsr* gene from *Piromyces* sp., for which the acquisition via HGT was previously shown [412].

Altogether, these analyses showed that the phylogeny of fungal Msrs globally matched the phylogeny of the fungi, in accordance with vertical inheritance of the genes from bacterial ancestors [8,9,24,26]. However, a few horizontal gene transfer events occurred in each Msr family, that contributed to the distribution of Msr genes in extant species.

## 4. Discussion

This global genomic search for *msr* genes in nearly 700 fungal genomes covering the fungal kingdom showed that most fungi contain one gene coding for each thiol-oxidoreductase type, i.e., MsrA, MsrB and fRMsr. The phylogenetic analyses and inspection of protein sequence features revealed that Msrs from each type are globally strongly conserved across the fungal kingdom. This is consistent with the prokaryotic origin of these genes. Yet, the identification of *fRmsr* genes in almost all genomes across the fungal kingdom was surprising. Indeed, no *fRmsr* genes were identified so far from multicellular eukaryotes [8]. Very likely, the most obvious reason was the smaller number of eukaryotic genomes considered (i.e. 160), and the smaller number of fungal genomes available at the time of the previous study [8, 50]. Because the number of sequenced genomes has increased dramatically in the last years, we took the opportunity of this study to reevaluate the presence of *fRmsr* genes in other multicellular eukaryotes. We searched for *fRmsr* genes in the plant and animal genomes available in the nr database of NCBI by BLAST search and found only a handful of multicellular eukaryotes apparently possessing a gene coding for a fRMsr (**Table S1**). These organisms, a plant and a few insects, are not phylogenetically related, indicating that the gene is not conserved in their lineage. This highlights the prevalence of *fRmsr* genes in the fungal kingdom, as opposite to their extremely low occurrence in other multicellular eukaryotes. What would be the advantage for the fungi to produce fRMsr whereas other multicellular organisms do not? An obvious possibility would be to protect the intracellular free Met during exposure to oxidative constraints as it was shown for *S. cerevisiae* [8]. Another possibility would be to allow the reduction of the free MetO coming from the external environment or after the degradation of proteins assimilated by the fungi and its use in protein synthesis and in sulfur metabolism. Combined with MsrA, the presence of fRMsr could virtually allow to reduce the complete pool of MetO coming from the external environment. This hypothesis arose from the observation of the presence of *fRmsr* genes in four Neocallimastigomycetes species that lacked MsrA and MsrB. Living in the anaerobic environment of ruminant gut, the fungi very likely do not suffer oxidative stress and the fRMsr could reduce the MetO coming from the animal’s food. Interestingly, the *fRmsr* genes in these Neocallimastigomycetes have been probably acquired through horizontal gene transfer, reinforcing the idea that a fRMsr would provide a selective advantage in an environment unlikely to generate oxidative constraints.

We found few variations in the numbers of gene copies of each type of Msr, with only 74 genomes lacking one gene, and 72 genomes with an extra copy of at least one *msr* gene. In both cases, it roughly corresponded to 10 percent of the analyzed genomes. Our study revealed the absence of both MsrA and MsrB in 15 species of fungi, twelve of these being also devoid of fRMsr. The lack of all Msr appears to be related to the fungal lifestyles, since it concerned the nine Microsporidia, and the *Pneumocystis jirovecii* species, which live as intracellular parasites of metazoans. These 10 species have small genomes with reduced numbers of genes, from 1,831 for *Encephalitozoon romaleae* SJ-2008 to 3,632 for *Enterocytozoon bieneusi* H348, as compared to the average ∼12,200 genes for the fungal genomes considered in this study. It has been proposed that the intracellular lifestyle allowed for genome compaction and gene loss, making the fungi highly dependent on the infected cell for numerous biochemical pathways [27]. Our results suggest the fungi could also rely on the host detoxication system for the protection against oxidative constraints and limitation of protein oxidation. The absence of Msr was also observed in Neocallimastigomycetes, which live in the anaerobic environment of ruminant gut, where the protection of proteins from oxidative damage is likely not crucial. This hypothesis is reinforced by the fact that the numbers of genes in Neocallimastigomycetes genomes is above the average of the analyzed fungi (∼15,500) and thus, the loss of *msr* genes was not due to a global genome compaction, but potentially due to the lack of selection pressure. Besides these species, for which the lack of Msr is consistent with the lifestyle, we also identified 12 other species for which *msrA* or *msrB* genes were absent from the genomes. As they are very likely living in aerobic conditions and are probably exposed to protein oxidation, the lack of MsrA or MsrB is surprising and we cannot exclude that the missing genes are due to incomplete genome sequencing. On the other hand, most of the 37 analyzed genomes from Glomeromycotina, Pucciniomycotina (*Melampsora* and *Puccinia*) and Taphrinomycotina (*Schizosaccharomyces*) species lacked the fRMsr. These fungi may have lost the capacity to reduce the free Met-*R*-O, as it has been shown for mammals [416], or the MsrB might have significant activity on the free Met-*R*-O, as suggested for plants [417].

Regarding the presence of supernumerary copies of *msr* genes in few genomes, it seems not to be related to the phylogeny, nor to the numbers of genes per genome, as the genomes of fungi having two copies or more of one *msr* gene contain an average number of ∼12,700 genes, similar to the average number of genes in the considered genomes (∼12,200). In most cases, the presence of several copies was due to gene duplications, as indicated by the close phylogenetic relationships of paralogous proteins and the conservation of canonical features. However, in few fungi, the presence of extra copies was due to horizontal gene transfers, like for *msrsA* genes in *Gonapodya prolifera*, and *msrB* genes in *Arthrobotrys oligospora*, *Monacrosporium haptotylum* and *Protomyces lactucaedebilis*. Together with the horizontal transfers of *fRmsr* genes in *Neocallimastigomycetes*, most of these events were strongly supported by the phylogeny and parametric values as well as by shared ecological niches for donor bacteria and recipient fungi. The case of *Gonapodya prolifera msrA* was remarkably interesting as it illustrated a prokaryote-to-eukaryote transfer of a selenoprotein gene. If such a transfer was observed from bacteria to archaea [418], to our knowledge, the bacteria-to-eukaryote transfer of a selenoprotein gene has never been described. As this fungus possesses an eukaryotic selenocysteine insertion machinery [402], we hypothesize that the transferred prokaryotic *msrA* gene was compatible with the eukaryotic machinery, or that it was modified after the transfer to allow proper Sec insertion. In all cases, the presence of several Msr gene copies could allow a beneficial increase in gene dosage or, alternatively, different spatial and temporal expression profiles during the life cycle of the fungi through distinct transcriptional activities. It could also allow different subcellular targeting as shown for plant Msrs [419]. Indeed, in 23 fungal genomes, out of the 59 having more than one *msrA* gene, the MsrAs were predicted to be addressed to different subcellular compartments. Similarly, MsrBs from 7 genomes out of the 15 with more than one gene were predicted to be addressed to different cellular compartments. Conversely, all the multicopy fRMsrs were predicted to be cytoplasmic. Our results indicate that, as experimentally demonstrated for yeast [39], most fungi might not have the ability to reduce the two diastereomers of protein-bound MetO in each subcellular compartment, suggesting variations in protein oxidation in the different cellular compartments. Another interesting aspect is the presence of potentially secreted Msrs in few fungal species, mostly from the Ascomycota genera *Aspergillus* and *Penicillium*. Although it remains to be demonstrated experimentally, the presence of Msrs in the extracellular environment of the fungal cells could help to protect from oxidation the numerous secreted carbohydrate-active enzymes (‘*CAZymes*’) and other enzymes used by saprotrophic and pathogen fungi to degrade plant cell walls or insect chitins [231].

Finally, we observed that most MsrAs, MsrBs and fRMsrs had conserved canonical features, but we also highlighted potentially interesting discrepancies. For the three types of Msrs, a few sequences lacked the resolving Cys at conserved positions and might use alternative Cys residues for the regeneration of their activity, whereas others possessed only the catalytic Cys and might be regenerated by the direct reduction of the sulfenic acid formed after MetO reduction, as shown for Msrs from other organisms [20, 408]. The presence of Sec in two *Gonapodya prolifera* MsrAs might confer them a catalytic advantage, similarly to other Sec-MsrAs [400]. We also observed that ∼5% of fungal MsrAs harbor a GCYW motif containing the catalytic Cys, where the Tyr residue replaces the Phe residue of the canonical GCFW motif. To our knowledge, no MsrA with such motif has been described to date. Because of the similarity between Phe and Tyr physicochemical properties, this substitution should not prevent the catalysis, but it could induce a change in substrate specificity, as demonstrated for the *E. coli* MsrA. Indeed, direct evolution assays showed that the substitution of the Phe by a Leu conferred to the enzyme the capacity to efficiently reduce alkyl-aryl sulfoxides [420]. Similarly, the atypical HYCIN motif of the *Protomyces lactucaedebilis* MsrB could modify the enzyme substrate specificity. The substitution of residues around the catalytic Cys of Msrs could affect the specificity for oxidized protein substrates or free MetO, but also potentially confer to the enzymes the ability to reduce other molecules such as the oxidized thioether-containing metabolites involved in sulfur metabolism (e.g S-adenosyl homocysteine, methylthio-ribose…) [421]. It would be interesting to evaluate the catalytic properties of these atypical Msrs on various sulfoxide-containing substrates.

## Acknowledgements

The authors thank the U.S. Department of Energy Joint Genome Institute (JGI), Dr. Sebastien Duplessis (Université de Lorraine, INRAE, UMR1136, Interactions Arbres/Microorganismes, France) of and Dr. Christine Smart (Cornell University, NY, USA) for sharing *Melampsora allii-populina* 12AY07 v1.0 and *Melampsora americana* R15-033-03 v1.0 unpublished genome data, respectively. Financial support to Hayat Hage from Région SUD Provence-Alpes-Côte d’Azur and The French National Research Institute for Agriculture, Food and Environment (INRAE) is acknowledged.

## Declaration of competing interest

None.

## Notes

### Competing Interest Statement

The authors have declared no competing interest.

## References

1. H. Sies, D.P. Jones, Reactive oxygen species (ROS) as pleiotropic physiological signalling agents, Nat. Rev. Mol. Cell Biol. 21 (2020) 363–383. https://doi.org/10.1038/s41580-020-0230-3.

2. V.S. Sharov, D.A. Ferrington, T.C. Squier, C. Schöneich, Diastereoselective reduction of protein-bound methionine sulfoxide by methionine sulfoxide reductase, FEBS Lett. 455 (1999) 247–250.

3. N. Brot, L. Weissbach, J. Werth, H. Weissbach, Enzymatic reduction of protein-bound methionine sulfoxide, Proc. Natl. Acad. Sci. U. S. A. 78 (1981) 2155–2158.

4. R. Grimaud, B. Ezraty, J.K. Mitchell, D. Lafitte, C. Briand, P.J. Derrick, F. Barras, Repair of oxidized proteins. Identification of a new methionine sulfoxide reductase, J. Biol. Chem. 276 (2001) 48915–48920. https://doi.org/10.1074/jbc.M105509200.

5. V.S. Sharov, C. Schöneich, Diastereoselective protein methionine oxidation by reactive oxygen species and diastereoselective repair by methionine sulfoxide reductase, Free Radic. Biol. Med. 29 (2000) 986–994.

6. L. Tarrago, A. Kaya, E. Weerapana, S.M. Marino, V.N. Gladyshev, Methionine sulfoxide reductases preferentially reduce unfolded oxidized proteins and protect cells from oxidative protein unfolding, J. Biol. Chem. 287 (2012) 24448–24459. https://doi.org/10.1074/jbc.M112.374520.

7. A. Gruez, M. Libiad, S. Boschi-Muller, G. Branlant, Structural and biochemical characterization of free methionine-R-sulfoxide reductase from Neisseria meningitidis, J. Biol. Chem. 285 (2010) 25033–25043. https://doi.org/10.1074/jbc.M110.134528.

8. D.T. Le, B.C. Lee, S.M. Marino, Y. Zhang, D.E. Fomenko, A. Kaya, E. Hacioglu, G.-H. Kwak, A. Koc, H.-Y. Kim, V.N. Gladyshev, Functional analysis of free methionine-R-sulfoxide reductase from Saccharomyces cerevisiae, J. Biol. Chem. 284 (2009) 4354–4364. https://doi.org/10.1074/jbc.M805891200.

9. Z. Lin, L.C. Johnson, H. Weissbach, N. Brot, M.O. Lively, W.T. Lowther, Free methionine-(R)-sulfoxide reductase from Escherichia coli reveals a new GAF domain function, Proc. Natl. Acad. Sci. U. S. A. 104 (2007) 9597–9602. https://doi.org/10.1073/pnas.0703774104.

10. R. Dhouib, D.S.M.P. Othman, V. Lin, X.J. Lai, H.G.S. Wijesinghe, A.-T. Essilfie, A. Davis, M. Nasreen, P.V. Bernhardt, P.M. Hansbro, A.G. McEwan, U. Kappler, A Novel, Molybdenum-Containing Methionine Sulfoxide Reductase Supports Survival of Haemophilus influenzae in an In vivo Model of Infection, Front. Microbiol. 7 (2016) 1743. https://doi.org/10.3389/fmicb.2016.01743.

11. B. Ezraty, J. Bos, F. Barras, L. Aussel, Methionine sulfoxide reduction and assimilation in Escherichia coli: new role for the biotin sulfoxide reductase BisC, J. Bacteriol. 187 (2005) 231– 237. https://doi.org/10.1128/JB.187.1.231-237.2005.

12. C. Andrieu, A. Vergnes, L. Loiseau, L. Aussel, B. Ezraty, Characterisation of the periplasmic methionine sulfoxide reductase (MsrP) from Salmonella Typhimurium, Free Radic. Biol. Med. 160 (2020) 506–512. https://doi.org/10.1016/j.freeradbiomed.2020.06.031.

13. A. Gennaris, B. Ezraty, C. Henry, R. Agrebi, A. Vergnes, E. Oheix, J. Bos, P. Leverrier, L. Espinosa, J. Szewczyk, D. Vertommen, O. Iranzo, J.-F. Collet, F. Barras, Repairing oxidized proteins in the bacterial envelope using respiratory chain electrons, Nature. 528 (2015) 409–412. https://doi.org/10.1038/nature15764.

14. L. Tarrago, S. Grosse, D. Lemaire, L. Faure, M. Tribout, M.I. Siponen, M. Kojadinovic-Sirinelli, D. Pignol, P. Arnoux, M. Sabaty, Reduction of Protein Bound Methionine Sulfoxide by a Periplasmic Dimethyl Sulfoxide Reductase, Antioxid. Basel Switz. 9 (2020). https://doi.org/10.3390/antiox9070616.

15. L. Tarrago, S. Grosse, M.I. Siponen, D. Lemaire, B. Alonso, G. Miotello, J. Armengaud, P. Arnoux, D. Pignol, M. Sabaty, Rhodobactersphaeroides methionine sulfoxide reductase P reduces R- and S-diastereomers of methionine sulfoxide from a broad-spectrum of protein substrates, Biochem. J. 475 (2018) 3779–3795. https://doi.org/10.1042/BCJ20180706.

16. W.T. Lowther, H. Weissbach, F. Etienne, N. Brot, B.W. Matthews, The mirrored methionine sulfoxide reductases of Neisseria gonorrhoeae pilB, Nat. Struct. Biol. 9 (2002) 348–352. https://doi.org/10.1038/nsb783.

17. S. Boschi-Muller, S. Azza, S. Sanglier-Cianferani, F. Talfournier, A. Van Dorsselear, G. Branlant, A sulfenic acid enzyme intermediate is involved in the catalytic mechanism of peptide methionine sulfoxide reductase from Escherichia coli, J. Biol. Chem. 275 (2000) 35908–35913. https://doi.org/10.1074/jbc.M006137200.

18. L. Tarrago, V.N. Gladyshev, Recharging oxidative protein repair: catalysis by methionine sulfoxide reductases towards their amino acid, protein, and model substrates, Biochem. Biokhimiia. 77 (2012) 1097–1107. https://doi.org/10.1134/S0006297912100021.

19. H.-Y. Kim, Glutaredoxin serves as a reductant for methionine sulfoxide reductases with or without resolving cysteine, Acta Biochim. Biophys. Sin. 44 (2012) 623–627. https://doi.org/10.1093/abbs/gms038.

20. L. Tarrago, E. Laugier, M. Zaffagnini, C.H. Marchand, P. Le Maréchal, S.D. Lemaire, P. Rey, Plant thioredoxin CDSP32 regenerates 1-cys methionine sulfoxide reductase B activity through the direct reduction of sulfenic acid, J. Biol. Chem. 285 (2010) 14964–14972. https://doi.org/10.1074/jbc.M110.108373.

21. L. Tarrago, E. Laugier, M. Zaffagnini, C. Marchand, P. Le Maréchal, N. Rouhier, S.D. Lemaire, P. Rey, Regeneration mechanisms of Arabidopsis thaliana methionine sulfoxide reductases B by glutaredoxins and thioredoxins, J. Biol. Chem. 284 (2009) 18963–18971. https://doi.org/10.1074/jbc.M109.015487.

22. W.T. Lowther, N. Brot, H. Weissbach, J.F. Honek, B.W. Matthews, Thiol-disulfide exchange is involved in the catalytic mechanism of peptide methionine sulfoxide reductase, Proc. Natl. Acad. Sci. U. S. A. 97 (2000) 6463–6468.

23. N. Rouhier, B. Kauffmann, F. Tete-Favier, P. Palladino, P. Gans, G. Branlant, J.-P. Jacquot, S. Boschi-Muller, Functional and structural aspects of poplar cytosolic and plastidial type a methionine sulfoxide reductases, J. Biol. Chem. 282 (2007) 3367–3378. https://doi.org/10.1074/jbc.M605007200.

24. L. Delaye, A. Becerra, L. Orgel, A. Lazcano, Molecular evolution of peptide methionine sulfoxide reductases (MsrA and MsrB): on the early development of a mechanism that protects against oxidative damage, J. Mol. Evol. 64 (2007) 15–32. https://doi.org/10.1007/s00239-005-0281-2.

25. L. Tarrago, E. Laugier, P. Rey, Protein-repairing methionine sulfoxide reductases in photosynthetic organisms: gene organization, reduction mechanisms, and physiological roles, Mol. Plant. 2 (2009) 202–217. https://doi.org/10.1093/mp/ssn067.

26. X.-H. Zhang, H. Weissbach, Origin and evolution of the protein-repairing enzymes methionine sulphoxide reductases, Biol. Rev. Camb. Philos. Soc. 83 (2008) 249–257. https://doi.org/10.1111/j.1469-185X.2008.00042.x.

27. M.D. Katinka, S. Duprat, E. Cornillot, G. Méténier, F. Thomarat, G. Prensier, V. Barbe, E. Peyretaillade, P. Brottier, P. Wincker, F. Delbac, H. El Alaoui, P. Peyret, W. Saurin, M. Gouy, J. Weissenbach, C.P. Vivarès, Genome sequence and gene compaction of the eukaryote parasite Encephalitozoon cuniculi, Nature. 414 (2001) 450–453. https://doi.org/10.1038/35106579.

28. A. Drazic, H. Miura, J. Peschek, Y. Le, N.C. Bach, T. Kriehuber, J. Winter, Methionine oxidation activates a transcription factor in response to oxidative stress, Proc. Natl. Acad. Sci. U. S. A. 110 (2013) 9493–9498. https://doi.org/10.1073/pnas.1300578110.

29. J.R. Erickson, M.A. Joiner, X. Guan, W. Kutschke, J. Yang, C.V. Oddis, R.K. Bartlett, J.S. Lowe, S.E. O’Donnell, N. Aykin-Burns, M.C. Zimmerman, K. Zimmerman, A.-J.L. Ham, R.M. Weiss, D.R. Spitz, M.A. Shea, R.J. Colbran, P.J. Mohler, M.E. Anderson, A dynamic pathway for calcium-independent activation of CaMKII by methionine oxidation, Cell. 133 (2008) 462–474. https://doi.org/10.1016/j.cell.2008.02.048.

30. B.C. Lee, Z. Péterfi, F.W. Hoffmann, R.E. Moore, A. Kaya, A. Avanesov, L. Tarrago, Y. Zhou, E. Weerapana, D.E. Fomenko, P.R. Hoffmann, V.N. Gladyshev, MsrB1 and MICALs regulate actin assembly and macrophage function via reversible stereoselective methionine oxidation, Mol. Cell. 51 (2013) 397–404. https://doi.org/10.1016/j.molcel.2013.06.019.

31. B.C. Lee, H.M. Lee, S. Kim, A.S. Avanesov, A. Lee, B.-H. Chun, G. Vorbruggen, V.N. Gladyshev, Expression of the methionine sulfoxide reductase lost during evolution extends Drosophila lifespan in a methionine-dependent manner, Sci. Rep. 8 (2018) 1–11. https://doi.org/10.1038/s41598-017-15090-5.

32. B. Jiang, J. Moskovitz, The Functions of the Mammalian Methionine Sulfoxide Reductase System and Related Diseases, Antioxid. Basel Switz. 7 (2018). https://doi.org/10.3390/antiox7090122.

33. J.M. Lim, G. Kim, R.L. Levine, Methionine in Proteins: It’s Not Just for Protein Initiation Anymore, Neurochem. Res. (2018). https://doi.org/10.1007/s11064-017-2460-0.

34. S. Lourenço dos Santos, I. Petropoulos, B. Friguet, The Oxidized Protein Repair Enzymes Methionine Sulfoxide Reductases and Their Roles in Protecting against Oxidative Stress, in Ageing and in Regulating Protein Function, Antioxidants. 7 (2018) 191. https://doi.org/10.3390/antiox7120191.

35. P. Rey, L. Tarrago, Physiological Roles of Plant Methionine Sulfoxide Reductases in Redox Homeostasis and Signaling, Antioxidants. 7 (2018) 114. https://doi.org/10.3390/antiox7090114.

36. V.K. Singh, K. Singh, K. Baum, The Role of Methionine Sulfoxide Reductases in Oxidative Stress Tolerance and Virulence of Staphylococcus aureus and Other Bacteria, Antioxidants. 7 (2018). https://doi.org/10.3390/antiox7100128.

37. M.A. Naranjo-Ortiz, T. Gabaldón, Fungal evolution: diversity, taxonomy and phylogeny of the Fungi, Biol. Rev. Camb. Philos. Soc. 94 (2019) 2101–2137. https://doi.org/10.1111/brv.12550.

38. M. Breitenbach, M. Weber, M. Rinnerthaler, T. Karl, L. Breitenbach-Koller, Oxidative Stress in Fungi: Its Function in Signal Transduction, Interaction with Plant Hosts, and Lignocellulose Degradation, Biomolecules. 5 (2015) 318–342. https://doi.org/10.3390/biom5020318.

39. A. Kaya, A. Koc, B.C. Lee, D.E. Fomenko, M. Rederstorff, A. Krol, A. Lescure, V.N. Gladyshev, Compartmentalization and regulation of mitochondrial function by methionine sulfoxide reductases in yeast, Biochemistry. 49 (2010) 8618–8625. https://doi.org/10.1021/bi100908v.

40. E.E. Nicklow, C.S. Sevier, Activity of the yeast cytoplasmic Hsp70 nucleotide-exchange factor Fes1 is regulated by reversible methionine oxidation, J. Biol. Chem. 295 (2020) 552–569. https://doi.org/10.1074/jbc.RA119.010125.

41. A. Koc, A.P. Gasch, J.C. Rutherford, H.-Y. Kim, V.N. Gladyshev, Methionine sulfoxide reductase regulation of yeast lifespan reveals reactive oxygen species-dependent and -independent components of aging, Proc. Natl. Acad. Sci. U. S. A. 101 (2004) 7999–8004. https://doi.org/10.1073/pnas.0307929101.

42. J. Moskovitz, E. Flescher, B.S. Berlett, J. Azare, J.M. Poston, E.R. Stadtman, Overexpression of peptide-methionine sulfoxide reductase in Saccharomyces cerevisiae and human T cells provides them with high resistance to oxidative stress, Proc. Natl. Acad. Sci. U. S. A. 95 (1998) 14071–14075.

43. H. Jo, Y.-W. Cho, S.-Y. Ji, G.-Y. Kang, C.-J. Lim, Protective roles of methionine-R-sulfoxide reductase against stresses in Schizosaccharomyces pombe, J. Basic Microbiol. 54 (2014) 72–80. https://doi.org/10.1002/jobm.201200397.

44. C. Yin, L. Zheng, J. Zhu, L. Chen, A. Ma, Enhancing stress tolerance by overexpression of a methionine sulfoxide reductase A (MsrA) gene in Pleurotus ostreatus, Appl. Microbiol. Biotechnol. 99 (2015) 3115–3126. https://doi.org/10.1007/s00253-014-6365-4.

45. F.M. Soriani, M.R. Kress, P. Fagundes de Gouvêa, I. Malavazi, M. Savoldi, A. Gallmetzer, J. Strauss, M.H.S. Goldman, G.H. Goldman, Functional characterization of the Aspergillus nidulans methionine sulfoxide reductases (msrA and msrB), Fungal Genet. Biol. FG B. 46 (2009) 410–417.

46. M. Kato, Y.-S. Yang, B.M. Sutter, Y. Wang, S.L. McKnight, B.P. Tu, Redox State Controls Phase Separation of the Yeast Ataxin-2 Protein via Reversible Oxidation of Its Methionine-Rich Low-Complexity Domain, Cell. 177 (2019) 711–721.e8. https://doi.org/10.1016/j.cell.2019.02.044.

47. A.M. Kachurin, A.M. Golubev, M.M. Geisow, O.S. Veselkina, L.S. Isaeva-Ivanova, K.N. Neustroev, Role of methionine in the active site of alpha-galactosidase from Trichoderma reesei., Biochem. J. 308 (1995) 955–964.

48. A. Gallmetzer, L. Silvestrini, T. Schinko, B. Gesslbauer, P. Hortschansky, C. Dattenböck, M.I. Muro-Pastor, A. Kungl, A.A. Brakhage, C. Scazzocchio, J. Strauss, Reversible Oxidation of a Conserved Methionine in the Nuclear Export Sequence Determines Subcellular Distribution and Activity of the Fungal Nitrate Regulator NirA, PLoS Genet. 11 (2015) e1005297. https://doi.org/10.1371/journal.pgen.1005297.

49. S.F. Altschul, W. Gish, W. Miller, E.W. Myers, D.J. Lipman, Basic local alignment search tool, J. Mol. Biol. 215 (1990) 403–410. https://doi.org/10.1016/S0022-2836(05)80360-2.

50. I.V. Grigoriev, R. Nikitin, S. Haridas, A. Kuo, R. Ohm, R. Otillar, R. Riley, A. Salamov, X. Zhao, F. Korzeniewski, T. Smirnova, H. Nordberg, I. Dubchak, I. Shabalov, MycoCosm portal: gearing up for 1000 fungal genomes, Nucleic Acids Res. 42 (2014) D699–D704. https://doi.org/10.1093/nar/gkt1183.

51. S. Haridas, R. Albert, M. Binder, J. Bloem, K. LaButti, A. Salamov, B. Andreopoulos, S.E. Baker, K. Barry, G. Bills, B.H. Bluhm, C. Cannon, R. Castanera, D.E. Culley, C. Daum, D. Ezra, J.B. González, B. Henrissat, A. Kuo, C. Liang, A. Lipzen, F. Lutzoni, J. Magnuson, S.J. Mondo, M. Nolan, R.A. Ohm, J. Pangilinan, H.-J. Park, L. Ramírez, M. Alfaro, H. Sun, A. Tritt, Y. Yoshinaga, L.-H. Zwiers, B.G. Turgeon, S.B. Goodwin, J.W. Spatafora, P.W. Crous, I.V. Grigoriev, 101 Dothideomycetes genomes: A test case for predicting lifestyles and emergence of pathogens, Stud. Mycol. 96 (2020) 141–153. https://doi.org/10.1016/j.simyco.2020.01.003.

52. T. Kijpornyongpan, S.J. Mondo, K. Barry, L. Sandor, J. Lee, A. Lipzen, J. Pangilinan, K. LaButti, M. Hainaut, B. Henrissat, I.V. Grigoriev, J.W. Spatafora, M.C. Aime, Broad Genomic Sampling Reveals a Smut Pathogenic Ancestry of the Fungal Clade Ustilaginomycotina, Mol. Biol. Evol. 35 (2018) 1840–1854. https://doi.org/10.1093/molbev/msy072.

53. A.C. Mosier, C.S. Miller, K.R. Frischkorn, R.A. Ohm, Z. Li, K. LaButti, A. Lapidus, A. Lipzen, C. Chen, J. Johnson, E.A. Lindquist, C. Pan, R.L. Hettich, I.V. Grigoriev, S.W. Singer, J.F. Banfield, Fungi Contribute Critical but Spatially Varying Roles in Nitrogen and Carbon Cycling in Acid Mine Drainage, Front. Microbiol. 7 (2016) 238. https://doi.org/10.3389/fmicb.2016.00238.

54. D. Terfehr, T.A. Dahlmann, T. Specht, I. Zadra, H. Kürnsteiner, U. Kück, Genome Sequence and Annotation of Acremonium chrysogenum, Producer of the β-Lactam Antibiotic Cephalosporin C, Genome Announc. 2 (2014). https://doi.org/10.1128/genomeA.00948-14.

55. E. Morin, S. Miyauchi, H. San Clemente, E.C.H. Chen, A. Pelin, I. de la Providencia, S. Ndikumana, D. Beaudet, M. Hainaut, E. Drula, A. Kuo, N. Tang, S. Roy, J. Viala, B. Henrissat, I.V. Grigoriev, N. Corradi, C. Roux, F.M. Martin, Comparative genomics of Rhizophagus irregularis, R. cerebriforme, R. diaphanus and Gigaspora rosea highlights specific genetic features in Glomeromycotina, New Phytol. 222 (2019) 1584–1598. https://doi.org/10.1111/nph.15687.

56. A. Kohler, A. Kuo, L.G. Nagy, E. Morin, K.W. Barry, F. Buscot, B. Canbäck, C. Choi, N. Cichocki, A. Clum, J. Colpaert, A. Copeland, M.D. Costa, J. Doré, D. Floudas, G. Gay, M. Girlanda, B. Henrissat, S. Herrmann, J. Hess, N. Högberg, T. Johansson, H.-R. Khouja, K. LaButti, U. Lahrmann, A. Levasseur, E.A. Lindquist, A. Lipzen, R. Marmeisse, E. Martino, C. Murat, C.Y. Ngan, U. Nehls, J.M. Plett, A. Pringle, R.A. Ohm, S. Perotto, M. Peter, R. Riley, F. Rineau, J. Ruytinx, A. Salamov, F. Shah, H. Sun, M. Tarkka, A. Tritt, C. Veneault-Fourrey, A. Zuccaro, Mycorrhizal Genomics Initiative Consortium, A. Tunlid, I.V. Grigoriev, D.S. Hibbett, F. Martin, Convergent losses of decay mechanisms and rapid turnover of symbiosis genes in mycorrhizal mutualists, Nat. Genet. 47 (2015) 410–415. https://doi.org/10.1038/ng.3223.

57. J. Hess, I. Skrede, B.E. Wolfe, K. LaButti, R.A. Ohm, I.V. Grigoriev, A. Pringle, Transposable element dynamics among asymbiotic and ectomycorrhizal Amanita fungi, Genome Biol. Evol. 6 (2014) 1564–1578. https://doi.org/10.1093/gbe/evu121.

58. T. Krassowski, A.Y. Coughlan, X.-X. Shen, X. Zhou, J. Kominek, D.A. Opulente, R. Riley, I.V. Grigoriev, N. Maheshwari, D.C. Shields, C.P. Kurtzman, C.T. Hittinger, A. Rokas, K.H. Wolfe, Evolutionary instability of CUG-Leu in the genetic code of budding yeasts, Nat. Commun. 9 (2018) 1887. https://doi.org/10.1038/s41467-018-04374-7.

59. E. Martino, E. Morin, G.-A. Grelet, A. Kuo, A. Kohler, S. Daghino, K.W. Barry, N. Cichocki, A. Clum, R.B. Dockter, M. Hainaut, R.C. Kuo, K. LaButti, B.D. Lindahl, E.A. Lindquist, A. Lipzen, H.-R. Khouja, J. Magnuson, C. Murat, R.A. Ohm, S.W. Singer, J.W. Spatafora, M. Wang, C. Veneault-Fourrey, B. Henrissat, I.V. Grigoriev, F.M. Martin, S. Perotto, Comparative genomics and transcriptomics depict ericoid mycorrhizal fungi as versatile saprotrophs and plant mutualists, New Phytol. 217 (2018) 1213–1229. https://doi.org/10.1111/nph.14974.

60. C.H. Haitjema, S.P. Gilmore, J.K. Henske, K.V. Solomon, R. de Groot, A. Kuo, S.J. Mondo, A.A. Salamov, K. LaButti, Z. Zhao, J. Chiniquy, K. Barry, H.M. Brewer, S.O. Purvine, A.T. Wright, M. Hainaut, B. Boxma, T. van Alen, J.H.P. Hackstein, B. Henrissat, S.E. Baker, I.V. Grigoriev, M.A. O’Malley, A parts list for fungal cellulosomes revealed by comparative genomics, Nat. Microbiol. 2 (2017) 17087. https://doi.org/10.1038/nmicrobiol.2017.87.

61. C.H. Slamovits, N.M. Fast, J.S. Law, P.J. Keeling, Genome compaction and stability in microsporidian intracellular parasites, Curr. Biol. CB. 14 (2004) 891–896. https://doi.org/10.1016/j.cub.2004.04.041.

62. G. Sipos, A.N. Prasanna, M.C. Walter, E. O’Connor, B. Bálint, K. Krizsán, B. Kiss, J. Hess, T. Varga, J. Slot, R. Riley, B. Bóka, D. Rigling, K. Barry, J. Lee, S. Mihaltcheva, K. LaButti, A. Lipzen, R. Waldron, N.M. Moloney, C. Sperisen, L. Kredics, C. Vágvölgyi, A. Patrignani, D. Fitzpatrick, I. Nagy, S. Doyle, J.B. Anderson, I.V. Grigoriev, U. Güldener, M. Münsterkötter, L.G. Nagy, Genome expansion and lineage-specific genetic innovations in the forest pathogenic fungi Armillaria, Nat. Ecol. Evol. 1 (2017) 1931–1941. https://doi.org/10.1038/s41559-017-0347-8.

63. C. Collins, T.M. Keane, D.J. Turner, G. O’Keeffe, D.A. Fitzpatrick, S. Doyle, Genomic and proteomic dissection of the ubiquitous plant pathogen, Armillaria mellea: toward a new infection model system, J. Proteome Res. 12 (2013) 2552–2570. https://doi.org/10.1021/pr301131t.

64. J. Yang, L. Wang, X. Ji, Y. Feng, X. Li, C. Zou, J. Xu, Y. Ren, Q. Mi, J. Wu, S. Liu, Y. Liu, X. Huang, H. Wang, X. Niu, J. Li, L. Liang, Y. Luo, K. Ji, W. Zhou, Z. Yu, G. Li, Y. Liu, L. Li, M. Qiao, L. Feng, K.-Q. Zhang, Genomic and proteomic analyses of the fungus Arthrobotrys oligospora provide insights into nematode-trap formation, PLoS Pathog. 7 (2011) e1002179. https://doi.org/10.1371/journal.ppat.1002179.

65. C. Murat, T. Payen, B. Noel, A. Kuo, E. Morin, J. Chen, A. Kohler, K. Krizsán, R. Balestrini, C. Da Silva, B. Montanini, M. Hainaut, E. Levati, K.W. Barry, B. Belfiori, N. Cichocki, A. Clum, R.B. Dockter, L. Fauchery, J. Guy, M. Iotti, F. Le Tacon, E.A. Lindquist, A. Lipzen, F. Malagnac, A. Mello, V. Molinier, S. Miyauchi, J. Poulain, C. Riccioni, A. Rubini, Y. Sitrit, R. Splivallo, S. Traeger, M. Wang, L. Žifčáková, D. Wipf, A. Zambonelli, F. Paolocci, M. Nowrousian, S. Ottonello, P. Baldrian, J.W. Spatafora, B. Henrissat, L.G. Nagy, J.-M. Aury, P. Wincker, I.V. Grigoriev, P. Bonfante, F.M. Martin, Pezizomycetes genomes reveal the molecular basis of ectomycorrhizal truffle lifestyle, Nat. Ecol. Evol. 2 (2018) 1956–1965. https://doi.org/10.1038/s41559-018-0710-4.

66. S. Verma, R.K. Gazara, S. Nizam, S. Parween, D. Chattopadhyay, P.K. Verma, Draft genome sequencing and secretome analysis of fungal phytopathogen Ascochyta rabiei provides insight into the necrotrophic effector repertoire, Sci. Rep. 6 (2016) 24638. https://doi.org/10.1038/srep24638.

67. T.A. Gianoulis, M.A. Griffin, D.J. Spakowicz, B.F. Dunican, C.J. Alpha, A. Sboner, A.M. Sismour, C. Kodira, M. Egholm, G.M. Church, M.B. Gerstein, S.A. Strobel, Genomic analysis of the hydrocarbon-producing, cellulolytic, endophytic fungus Ascocoryne sarcoides, PLoS Genet. 8 (2012) e1002558. https://doi.org/10.1371/journal.pgen.1002558.

68. R. Lütkenhaus, S. Traeger, J. Breuer, L. Carreté, A. Kuo, A. Lipzen, J. Pangilinan, D. Dilworth, L. Sandor, S. Pöggeler, T. Gabaldón, K. Barry, I.V. Grigoriev, M. Nowrousian, Comparative Genomics and Transcriptomics To Analyze Fruiting Body Development in Filamentous Ascomycetes, Genetics. 213 (2019) 1545–1563. https://doi.org/10.1534/genetics.119.302749.

69. T.C. Vesth, J.L. Nybo, S. Theobald, J.C. Frisvad, T.O. Larsen, K.F. Nielsen, J.B. Hoof, J. Brandl, A. Salamov, R. Riley, J.M. Gladden, P. Phatale, M.T. Nielsen, E.K. Lyhne, M.E. Kogle, K. Strasser, E. McDonnell, K. Barry, A. Clum, C. Chen, K. LaButti, S. Haridas, M. Nolan, L. Sandor, A. Kuo, A. Lipzen, M. Hainaut, E. Drula, A. Tsang, J.K. Magnuson, B. Henrissat, A. Wiebenga, B.A. Simmons, M.R. Mäkelä, R.P. de Vries, I.V. Grigoriev, U.H. Mortensen, S.E. Baker, M.R. Andersen, Investigation of inter- and intraspecies variation through genome sequencing of Aspergillus section Nigri, Nat. Genet. 50 (2018) 1688–1695. https://doi.org/10.1038/s41588-018-0246-1.

70. R.P. de Vries, R. Riley, A. Wiebenga, G. Aguilar-Osorio, S. Amillis, C.A. Uchima, G. Anderluh, M. Asadollahi, M. Askin, K. Barry, E. Battaglia, Ö. Bayram, T. Benocci, S.A. Braus-Stromeyer, C. Caldana, D. Cánovas, G.C. Cerqueira, F. Chen, W. Chen, C. Choi, A. Clum, R.A.C. Dos Santos, A.R. de L. Damásio, G. Diallinas, T. Emri, E. Fekete, M. Flipphi, S. Freyberg, A. Gallo, C. Gournas, R. Habgood, M. Hainaut, M.L. Harispe, B. Henrissat, K.S. Hildén, R. Hope, A. Hossain, E. Karabika, L. Karaffa, Z. Karányi, N. Kraševec, A. Kuo, H. Kusch, K. LaButti, E.L. Lagendijk, A. Lapidus, A. Levasseur, E. Lindquist, A. Lipzen, A.F. Logrieco, A. MacCabe, M.R. Mäkelä, I. Malavazi, P. Melin, V. Meyer, N. Mielnichuk, M. Miskei, Á.P. Molnár, G. Mulé, C.Y. Ngan, M. Orejas, E. Orosz, J.P. Ouedraogo, K.M. Overkamp, H.-S. Park, G. Perrone, F. Piumi, P.J. Punt, A.F.J. Ram, A. Ramón, S. Rauscher, E. Record, D.M. Riaño-Pachón, V. Robert, J. Röhrig, R. Ruller, A. Salamov, N.S. Salih, R.A. Samson, E. Sándor, M. Sanguinetti, T. Schütze, K. Sepčić, E. Shelest, G. Sherlock, V. Sophianopoulou, F.M. Squina, H. Sun, A. Susca, R.B. Todd, A. Tsang, S.E. Unkles, N. van de Wiele, D. van Rossen-Uffink, J.V. de C. Oliveira, T.C. Vesth, J. Visser, J.-H. Yu, M. Zhou, M.R. Andersen, D.B. Archer, S.E. Baker, I. Benoit, A.A. Brakhage, G.H. Braus, R. Fischer, J.C. Frisvad, G.H. Goldman, J. Houbraken, B. Oakley, I. Pócsi, C. Scazzocchio, B. Seiboth, P.A. vanKuyk, J. Wortman, P.S. Dyer, I.V. Grigoriev, Comparative genomics reveals high biological diversity and specific adaptations in the industrially and medically important fungal genus Aspergillus, Genome Biol. 18 (2017) 28. https://doi.org/10.1186/s13059-017-1151-0.

71. G.G. Moore, B.M. Mack, S.B. Beltz, M.K. Gilbert, Draft Genome Sequence of an Aflatoxigenic Aspergillus Species, A. bombycis, Genome Biol. Evol. 8 (2016) 3297–3300. https://doi.org/10.1093/gbe/evw238.

72. F. Horn, J. Linde, D.J. Mattern, G. Walther, R. Guthke, K. Scherlach, K. Martin, A.A. Brakhage, L. Petzke, V. Valiante, Draft Genome Sequences of Fungus Aspergillus calidoustus, Genome Announc. 4 (2016). https://doi.org/10.1128/genomeA.00102-16.

73. I. Kjærbølling, T. Vesth, J.C. Frisvad, J.L. Nybo, S. Theobald, S. Kildgaard, T.I. Petersen, A. Kuo, A. Sato, E.K. Lyhne, M.E. Kogle, A. Wiebenga, R.S. Kun, R.J.M. Lubbers, M.R. Mäkelä, K. Barry, M. Chovatia, A. Clum, C. Daum, S. Haridas, G. He, K. LaButti, A. Lipzen, S. Mondo, J. Pangilinan, R. Riley, A. Salamov, B.A. Simmons, J.K. Magnuson, B. Henrissat, U.H. Mortensen, T.O. Larsen, R.P. de Vries, I.V. Grigoriev, M. Machida, S.E. Baker, M.R. Andersen, A comparative genomics study of 23 Aspergillus species from section Flavi, Nat. Commun. 11 (2020) 1106. https://doi.org/10.1038/s41467-019-14051-y.

74. M.B. Arnaud, G.C. Cerqueira, D.O. Inglis, M.S. Skrzypek, J. Binkley, M.C. Chibucos, J. Crabtree, C. Howarth, J. Orvis, P. Shah, F. Wymore, G. Binkley, S.R. Miyasato, M. Simison, G. Sherlock, J.R. Wortman, The Aspergillus Genome Database (AspGD): recent developments in comprehensive multispecies curation, comparative genomics and community resources, Nucleic Acids Res. 40 (2012) D653–659. https://doi.org/10.1093/nar/gkr875.

75. Y. Ge, Y. Wang, Y. Liu, Y. Tan, X. Ren, X. Zhang, K.D. Hyde, Y. Liu, Z. Liu, Comparative genomic and transcriptomic analyses of the Fuzhuan brick tea-fermentation fungus Aspergillus cristatus, BMC Genomics. 17 (2016) 428. https://doi.org/10.1186/s12864-016-2637-y.

76. N.D. Fedorova, N. Khaldi, V.S. Joardar, R. Maiti, P. Amedeo, M.J. Anderson, J. Crabtree, J.C. Silva, J.H. Badger, A. Albarraq, S. Angiuoli, H. Bussey, P. Bowyer, P.J. Cotty, P.S. Dyer, A. Egan, K. Galens, C.M. Fraser-Liggett, B.J. Haas, J.M. Inman, R. Kent, S. Lemieux, I. Malavazi, J. Orvis, T. Roemer, C.M. Ronning, J.P. Sundaram, G. Sutton, G. Turner, J.C. Venter, O.R. White, B.R. Whitty, P. Youngman, K.H. Wolfe, G.H. Goldman, J.R. Wortman, B. Jiang, D.W. Denning, W.C. Nierman, Genomic islands in the pathogenic filamentous fungus Aspergillus fumigatus, PLoS Genet. 4 (2008) e1000046. https://doi.org/10.1371/journal.pgen.1000046.

77. W.C. Nierman, A. Pain, M.J. Anderson, J.R. Wortman, H.S. Kim, J. Arroyo, M. Berriman, K. Abe, D.B. Archer, C. Bermejo, J. Bennett, P. Bowyer, D. Chen, M. Collins, R. Coulsen, R. Davies, P.S. Dyer, M. Farman, N. Fedorova, N. Fedorova, T.V. Feldblyum, R. Fischer, N. Fosker, A. Fraser, J.L. García, M.J. García, A. Goble, G.H. Goldman, K. Gomi, S. Griffith-Jones, R. Gwilliam, B. Haas, H. Haas, D. Harris, H. Horiuchi, J. Huang, S. Humphray, J. Jiménez, N. Keller, H. Khouri, K. Kitamoto, T. Kobayashi, S. Konzack, R. Kulkarni, T. Kumagai, A. Lafon, A. Lafton, J.-P. Latgé, W. Li, A. Lord, C. Lu, W.H. Majoros, G.S. May, B.L. Miller, Y. Mohamoud, M. Molina, M. Monod, I. Mouyna, S. Mulligan, L. Murphy, S. O’Neil, I. Paulsen, M.A. Peñalva, M. Pertea, C. Price, B.L. Pritchard, M.A. Quail, E. Rabbinowitsch, N. Rawlins, M.-A. Rajandream, U. Reichard, H. Renauld, G.D. Robson, S. Rodriguez de Córdoba, J.M. Rodríguez-Peña, C.M. Ronning, S. Rutter, S.L. Salzberg, M. Sanchez, J.C. Sánchez-Ferrero, D. Saunders, K. Seeger, R. Squares, S. Squares, M. Takeuchi, F. Tekaia, G. Turner, C.R. Vazquez de Aldana, J. Weidman, O. White, J. Woodward, J.-H. Yu, C. Fraser, J.E. Galagan, K. Asai, M. Machida, N. Hall, B. Barrell, D.W. Denning, Genomic sequence of the pathogenic and allergenic filamentous fungus Aspergillus fumigatus, Nature. 438 (2005) 1151– 1156. https://doi.org/10.1038/nature04332.

78. T. Futagami, K. Mori, A. Yamashita, S. Wada, Y. Kajiwara, H. Takashita, T. Omori, K. Takegawa, K. Tashiro, S. Kuhara, M. Goto, Genome sequence of the white koji mold Aspergillus kawachii IFO 4308, used for brewing the Japanese distilled spirit shochu, Eukaryot. Cell. 10 (2011) 1586– 1587. https://doi.org/10.1128/EC.05224-11.

79. M.R. Andersen, M.P. Salazar, P.J. Schaap, P.J.I. van de Vondervoort, D. Culley, J. Thykaer, J.C. Frisvad, K.F. Nielsen, R. Albang, K. Albermann, R.M. Berka, G.H. Braus, S.A. Braus-Stromeyer, L.M. Corrochano, Z. Dai, P.W.M. van Dijck, G. Hofmann, L.L. Lasure, J.K. Magnuson, H. Menke, M. Meijer, S.L. Meijer, J.B. Nielsen, M.L. Nielsen, A.J.J. van Ooyen, H.J. Pel, L. Poulsen, R.A. Samson, H. Stam, A. Tsang, J.M. van den Brink, A. Atkins, A. Aerts, H. Shapiro, J. Pangilinan, A. Salamov, Y. Lou, E. Lindquist, S. Lucas, J. Grimwood, I.V. Grigoriev, C.P. Kubicek, D. Martinez, N.N.M.E. van Peij, J.A. Roubos, J. Nielsen, S.E. Baker, Comparative genomics of citric-acid-producing Aspergillus niger ATCC 1015 versus enzyme-producing CBS 513.88, Genome Res. 21 (2011) 885–897. https://doi.org/10.1101/gr.112169.110.

80. H.J. Pel, J.H. de Winde, D.B. Archer, P.S. Dyer, G. Hofmann, P.J. Schaap, G. Turner, R.P. de Vries, R. Albang, K. Albermann, M.R. Andersen, J.D. Bendtsen, J.A.E. Benen, M. van den Berg, S. Breestraat, M.X. Caddick, R. Contreras, M. Cornell, P.M. Coutinho, E.G.J. Danchin, A.J.M. Debets, P. Dekker, P.W.M. van Dijck, A. van Dijk, L. Dijkhuizen, A.J.M. Driessen, C. d’Enfert, S. Geysens, C. Goosen, G.S.P. Groot, P.W.J. de Groot, T. Guillemette, B. Henrissat, M. Herweijer, J.P.T.W. van den Hombergh, C.A.M.J.J. van den Hondel, R.T.J.M. van der Heijden, R.M. van der Kaaij, F.M. Klis, H.J. Kools, C.P. Kubicek, P.A. van Kuyk, J. Lauber, X. Lu, M.J.E.C. van der Maarel, R. Meulenberg, H. Menke, M.A. Mortimer, J. Nielsen, S.G. Oliver, M. Olsthoorn, K. Pal, N.N.M.E. van Peij, A.F.J. Ram, U. Rinas, J.A. Roubos, C.M.J. Sagt, M. Schmoll, J. Sun, D. Ussery, J. Varga, W. Vervecken, P.J.J. van de Vondervoort, H. Wedler, H.A.B. Wösten, A.-P. Zeng, A.J.J. van Ooyen, J. Visser, H. Stam, Genome sequencing and analysis of the versatile cell factory Aspergillus niger CBS 513.88, Nat. Biotechnol. 25 (2007) 221–231. https://doi.org/10.1038/nbt1282.

81. G.G. Moore, B.M. Mack, S.B. Beltz, Draft Genome Sequences of Two Closely Related Aflatoxigenic Aspergillus Species Obtained from the Ivory Coast, Genome Biol. Evol. 8 (2015) 729–732. https://doi.org/10.1093/gbe/evv246.

82. Y. Kusuya, A. Takahashi-Nakaguchi, H. Takahashi, T. Yaguchi, Draft Genome Sequence of the Pathogenic Filamentous Fungus Aspergillus udagawae Strain IFM 46973T, Genome Announc. 3 (2015). https://doi.org/10.1128/genomeA.00834-15.

83. C. Gostinčar, R.A. Ohm, T. Kogej, S. Sonjak, M. Turk, J. Zajc, P. Zalar, M. Grube, H. Sun, J. Han, A. Sharma, J. Chiniquy, C.Y. Ngan, A. Lipzen, K. Barry, I.V. Grigoriev, N. Gunde-Cimerman, Genome sequencing of four Aureobasidium pullulans varieties: biotechnological potential, stress tolerance, and description of new species, BMC Genomics. 15 (2014) 549. https://doi.org/10.1186/1471-2164-15-549.

84. D. Floudas, M. Binder, R. Riley, K. Barry, R.A. Blanchette, B. Henrissat, A.T. Martínez, R. Otillar, J.W. Spatafora, J.S. Yadav, A. Aerts, I. Benoit, A. Boyd, A. Carlson, A. Copeland, P.M. Coutinho, R.P. de Vries, P. Ferreira, K. Findley, B. Foster, J. Gaskell, D. Glotzer, P. Górecki, J. Heitman, C. Hesse, C. Hori, K. Igarashi, J.A. Jurgens, N. Kallen, P. Kersten, A. Kohler, U. Kües, T.K.A. Kumar, A. Kuo, K. LaButti, L.F. Larrondo, E. Lindquist, A. Ling, V. Lombard, S. Lucas, T. Lundell, R. Martin, D.J. McLaughlin, I. Morgenstern, E. Morin, C. Murat, L.G. Nagy, M. Nolan, R.A. Ohm, A. Patyshakuliyeva, A. Rokas, F.J. Ruiz-Dueñas, G. Sabat, A. Salamov, M. Samejima, J. Schmutz, J.C. Slot, F. St John, J. Stenlid, H. Sun, S. Sun, K. Syed, A. Tsang, A. Wiebenga, D. Young, A. Pisabarro, D.C. Eastwood, F. Martin, D. Cullen, I.V. Grigoriev, D.S. Hibbett, The Paleozoic origin of enzymatic lignin decomposition reconstructed from 31 fungal genomes, Science. 336 (2012) 1715–1719. https://doi.org/10.1126/science.1221748.

85. É. Almási, N. Sahu, K. Krizsán, B. Bálint, G.M. Kovács, B. Kiss, J. Cseklye, E. Drula, B. Henrissat, I. Nagy, M. Chovatia, C. Adam, K. LaButti, A. Lipzen, R. Riley, I.V. Grigoriev, L.G. Nagy, Comparative genomics reveals unique wood-decay strategies and fruiting body development in the Schizophyllaceae, New Phytol. 224 (2019) 902–915. https://doi.org/10.1111/nph.16032.

86. R.A. Ohm, N. Feau, B. Henrissat, C.L. Schoch, B.A. Horwitz, K.W. Barry, B.J. Condon, A.C. Copeland, B. Dhillon, F. Glaser, C.N. Hesse, I. Kosti, K. LaButti, E.A. Lindquist, S. Lucas, A.A. Salamov, R.E. Bradshaw, L. Ciuffetti, R.C. Hamelin, G.H.J. Kema, C. Lawrence, J.A. Scott, J.W. Spatafora, B.G. Turgeon, P.J.G.M. de Wit, S. Zhong, S.B. Goodwin, I.V. Grigoriev, Diverse lifestyles and strategies of plant pathogenesis encoded in the genomes of eighteen Dothideomycetes fungi, PLoS Pathog. 8 (2012) e1003037. https://doi.org/10.1371/journal.ppat.1003037.

87. G. Xiao, S.-H. Ying, P. Zheng, Z.-L. Wang, S. Zhang, X.-Q. Xie, Y. Shang, R.J. St Leger, G.-P. Zhao, C. Wang, M.-G. Feng, Genomic perspectives on the evolution of fungal entomopathogenicity in Beauveria bassiana, Sci. Rep. 2 (2012) 483. https://doi.org/10.1038/srep00483.

88. B. Blanco-Ulate, P. Rolshausen, D. Cantu, Draft Genome Sequence of Neofusicoccum parvum Isolate UCR-NP2, a Fungal Vascular Pathogen Associated with Grapevine Cankers, Genome Announc. 1 (2013). https://doi.org/10.1128/genomeA.00339-13.

89. G. Kunze, C. Gaillardin, M. Czernicka, P. Durrens, T. Martin, E. Böer, T. Gabaldón, J.A. Cruz, E. Talla, C. Marck, A. Goffeau, V. Barbe, P. Baret, K. Baronian, S. Beier, C. Bleykasten, R. Bode, S. Casaregola, L. Despons, C. Fairhead, M. Giersberg, P.P. Gierski, U. Hähnel, A. Hartmann, D. Jankowska, C. Jubin, P. Jung, I. Lafontaine, V. Leh-Louis, M. Lemaire, M. Marcet-Houben, M. Mascher, G. Morel, G.-F. Richard, J. Riechen, C. Sacerdot, A. Sarkar, G. Savel, J. Schacherer, D.J. Sherman, N. Stein, M.-L. Straub, A. Thierry, A. Trautwein-Schult, B. Vacherie, E. Westhof, S. Worch, B. Dujon, J.-L. Souciet, P. Wincker, U. Scholz, C. Neuvéglise, The complete genome of Blastobotrys (Arxula) adeninivorans LS3 - a yeast of biotechnological interest, Biotechnol. Biofuels. 7 (2014) 66. https://doi.org/10.1186/1754-6834-7-66.

90. J.F. Muñoz, G.M. Gauthier, C.A. Desjardins, J.E. Gallo, J. Holder, T.D. Sullivan, A.J. Marty, J.C. Carmen, Z. Chen, L. Ding, S. Gujja, V. Magrini, E. Misas, M. Mitreva, M. Priest, S. Saif, E.A. Whiston, S. Young, Q. Zeng, W.E. Goldman, E.R. Mardis, J.W. Taylor, J.G. McEwen, O.K. Clay, B.S. Klein, C.A. Cuomo, The Dynamic Genome and Transcriptome of the Human Fungal Pathogen Blastomyces and Close Relative Emmonsia, PLoS Genet. 11 (2015) e1005493. https://doi.org/10.1371/journal.pgen.1005493.

91. L. Frantzeskakis, B. Kracher, S. Kusch, M. Yoshikawa-Maekawa, S. Bauer, C. Pedersen, P.D. Spanu, T. Maekawa, P. Schulze-Lefert, R. Panstruga, Signatures of host specialization and a recent transposable element burst in the dynamic one-speed genome of the fungal barley powdery mildew pathogen, BMC Genomics. 19 (2018) 381. https://doi.org/10.1186/s12864-018-4750-6.

92. T. Wicker, S. Oberhaensli, F. Parlange, J.P. Buchmann, M. Shatalina, S. Roffler, R. Ben-David, J. Doležel, H. Šimková, P. Schulze-Lefert, P.D. Spanu, R. Bruggmann, J. Amselem, H. Quesneville, E. Ver Loren van Themaat, T. Paape, K.K. Shimizu, B. Keller, The wheat powdery mildew genome shows the unique evolution of an obligate biotroph, Nat. Genet. 45 (2013) 1092–1096. https://doi.org/10.1038/ng.2704.

93. R. Riley, S. Haridas, K.H. Wolfe, M.R. Lopes, C.T. Hittinger, M. Göker, A.A. Salamov, J.H. Wisecaver, T.M. Long, C.H. Calvey, A.L. Aerts, K.W. Barry, C. Choi, A. Clum, A.Y. Coughlan, S. Deshpande, A.P. Douglass, S.J. Hanson, H.-P. Klenk, K.M. LaButti, A. Lapidus, E.A. Lindquist, A.M. Lipzen, J.P. Meier-Kolthoff, R.A. Ohm, R.P. Otillar, J.L. Pangilinan, Y. Peng, A. Rokas, C.A. Rosa, C. Scheuner, A.A. Sibirny, J.C. Slot, J.B. Stielow, H. Sun, C.P. Kurtzman, M. Blackwell, I.V. Grigoriev, T.W. Jeffries, Comparative genomics of biotechnologically important yeasts, Proc. Natl. Acad. Sci. U. S. A. 113 (2016) 9882–9887. https://doi.org/10.1073/pnas.1603941113.

94. A. Marsberg, M. Kemler, F. Jami, J.H. Nagel, A. Postma-Smidt, S. Naidoo, M.J. Wingfield, P.W. Crous, J.W. Spatafora, C.N. Hesse, B. Robbertse, B. Slippers, Botryosphaeria dothidea: a latent pathogen of global importance to woody plant health, Mol. Plant Pathol. 18 (2017) 477–488. https://doi.org/10.1111/mpp.12495.

95. M. Staats, J.A.L. van Kan, Genome update of Botrytis cinerea strains B05.10 and T4, Eukaryot. Cell. 11 (2012) 1413–1414. https://doi.org/10.1128/EC.00164-12.

96. T. Oka, K. Ekino, K. Fukuda, Y. Nomura, Draft Genome Sequence of the Formaldehyde-Resistant Fungus Byssochlamys spectabilis No. 5 (Anamorph Paecilomyces variotii No. 5) (NBRC109023), Genome Announc. 2 (2014). https://doi.org/10.1128/genomeA.01162-13.

97. T. Jones, N.A. Federspiel, H. Chibana, J. Dungan, S. Kalman, B.B. Magee, G. Newport, Y.R. Thorstenson, N. Agabian, P.T. Magee, R.W. Davis, S. Scherer, The diploid genome sequence of Candida albicans, Proc. Natl. Acad. Sci. U. S. A. 101 (2004) 7329–7334. https://doi.org/10.1073/pnas.0401648101.

98. D.J. Wohlbach, A. Kuo, T.K. Sato, K.M. Potts, A.A. Salamov, K.M. Labutti, H. Sun, A. Clum, J.L. Pangilinan, E.A. Lindquist, S. Lucas, A. Lapidus, M. Jin, C. Gunawan, V. Balan, B.E. Dale, T.W. Jeffries, R. Zinkel, K.W. Barry, I.V. Grigoriev, A.P. Gasch, Comparative genomics of xylose-fermenting fungi for enhanced biofuel production, Proc. Natl. Acad. Sci. U. S. A. 108 (2011) 13212–13217. https://doi.org/10.1073/pnas.1103039108.

99. M.M. Teixeira, L.F. Moreno, B.J. Stielow, A. Muszewska, M. Hainaut, L. Gonzaga, A. Abouelleil, J.S.L. Patané, M. Priest, R. Souza, S. Young, K.S. Ferreira, Q. Zeng, M.M.L. da Cunha, A. Gladki, B. Barker, V.A. Vicente, E.M. de Souza, S. Almeida, B. Henrissat, A.T.R. Vasconcelos, S. Deng, H. Voglmayr, T. a. A. Moussa, A. Gorbushina, M.S.S. Felipe, C.A. Cuomo, G.S. de Hoog, Exploring the genomic diversity of black yeasts and relatives (Chaetothyriales, Ascomycota), Stud. Mycol. 86 (2017) 1–28. https://doi.org/10.1016/j.simyco.2017.01.001.

100. M. Peter, A. Kohler, R.A. Ohm, A. Kuo, J. Krützmann, E. Morin, M. Arend, K.W. Barry, M. Binder, C. Choi, A. Clum, A. Copeland, N. Grisel, S. Haridas, T. Kipfer, K. LaButti, E. Lindquist, A. Lipzen, R. Maire, B. Meier, S. Mihaltcheva, V. Molinier, C. Murat, S. Pöggeler, C.A. Quandt, C. Sperisen, A. Tritt, E. Tisserant, P.W. Crous, B. Henrissat, U. Nehls, S. Egli, J.W. Spatafora, I.V. Grigoriev, F.M. Martin, Ectomycorrhizal ecology is imprinted in the genome of the dominant symbiotic fungus Cenococcum geophilum, Nat. Commun. 7 (2016) 12662. https://doi.org/10.1038/ncomms12662.

101. E. Fernandez-Fueyo, F.J. Ruiz-Dueñas, P. Ferreira, D. Floudas, D.S. Hibbett, P. Canessa, L.F. Larrondo, T.Y. James, D. Seelenfreund, S. Lobos, R. Polanco, M. Tello, Y. Honda, T. Watanabe, T. Watanabe, J.S. Ryu, R.J. San, C.P. Kubicek, M. Schmoll, J. Gaskell, K.E. Hammel, F.J. St John, A. Vanden Wymelenberg, G. Sabat, S. Splinter BonDurant, K. Syed, J.S. Yadav, H. Doddapaneni, V. Subramanian, J.L. Lavín, J.A. Oguiza, G. Perez, A.G. Pisabarro, L. Ramirez, F. Santoyo, E. Master, P.M. Coutinho, B. Henrissat, V. Lombard, J.K. Magnuson, U. Kües, C. Hori, K. Igarashi, M. Samejima, B.W. Held, K.W. Barry, K.M. LaButti, A. Lapidus, E.A. Lindquist, S.M. Lucas, R. Riley, A.A. Salamov, D. Hoffmeister, D. Schwenk, Y. Hadar, O. Yarden, R.P. de Vries, A. Wiebenga, J. Stenlid, D. Eastwood, I.V. Grigoriev, R.M. Berka, R.A. Blanchette, P. Kersten, A.T. Martinez, R. Vicuna, D. Cullen, Comparative genomics of Ceriporiopsis subvermispora and Phanerochaete chrysosporium provide insight into selective ligninolysis, Proc. Natl. Acad. Sci. U. S. A. 109 (2012) 5458–5463. https://doi.org/10.1073/pnas.1119912109.

102. C.A. Cuomo, W.A. Untereiner, L.-J. Ma, M. Grabherr, B.W. Birren, Draft Genome Sequence of the Cellulolytic Fungus Chaetomium globosum, Genome Announc. 3 (2015). https://doi.org/10.1128/genomeA.00021-15.

103. S. Amlacher, P. Sarges, D. Flemming, V. van Noort, R. Kunze, D.P. Devos, M. Arumugam, P. Bork, E. Hurt, Insight into structure and assembly of the nuclear pore complex by utilizing the genome of a eukaryotic thermophile, Cell. 146 (2011) 277–289. https://doi.org/10.1016/j.cell.2011.06.039.

104. D. Armaleo, O. Müller, F. Lutzoni, Ó.S. Andrésson, G. Blanc, H.B. Bode, F.R. Collart, F. Dal Grande, F. Dietrich, I.V. Grigoriev, S. Joneson, A. Kuo, P.E. Larsen, J.M. Logsdon, D. Lopez, F. Martin, S.P. May, T.R. McDonald, S.S. Merchant, V. Miao, E. Morin, R. Oono, M. Pellegrini, N. Rubinstein, M.V. Sanchez-Puerta, E. Savelkoul, I. Schmitt, J.C. Slot, D. Soanes, P. Szövényi, N.J. Talbot, C. Veneault-Fourrey, B.B. Xavier, The lichen symbiosis re-viewed through the genomes of Cladonia grayi and its algal partner Asterochloris glomerata, BMC Genomics. 20 (2019) 605. https://doi.org/10.1186/s12864-019-5629-x.

105. P.J.G.M. de Wit, A. van der Burgt, B. Ökmen, I. Stergiopoulos, K.A. Abd-Elsalam, A.L. Aerts, A.H. Bahkali, H.G. Beenen, P. Chettri, M.P. Cox, E. Datema, R.P. de Vries, B. Dhillon, A.R. Ganley, S.A. Griffiths, Y. Guo, R.C. Hamelin, B. Henrissat, M.S. Kabir, M.K. Jashni, G. Kema, S. Klaubauf, A. Lapidus, A. Levasseur, E. Lindquist, R. Mehrabi, R.A. Ohm, T.J. Owen, A. Salamov, A. Schwelm, E. Schijlen, H. Sun, H.A. van den Burg, R.C.H.J. van Ham, S. Zhang, S.B. Goodwin, I.V. Grigoriev, J. Collemare, R.E. Bradshaw, The genomes of the fungal plant pathogens Cladosporium fulvum and Dothistroma septosporum reveal adaptation to different hosts and lifestyles but also signatures of common ancestry, PLoS Genet. 8 (2012) e1003088. https://doi.org/10.1371/journal.pgen.1003088.

106. K.P. Ng, S.M. Yew, C.L. Chan, T.S. Soo-Hoo, S.L. Na, H. Hassan, Y.F. Ngeow, C.-C. Hoh, K.-W. Lee, W.-Y. Yee, Sequencing of Cladosporium sphaerospermum, a Dematiaceous fungus isolated from blood culture, Eukaryot. Cell. 11 (2012) 705–706. https://doi.org/10.1128/EC.00081-12.

107. G. Butler, M.D. Rasmussen, M.F. Lin, M.A.S. Santos, S. Sakthikumar, C.A. Munro, E. Rheinbay, M. Grabherr, A. Forche, J.L. Reedy, I. Agrafioti, M.B. Arnaud, S. Bates, A.J.P. Brown, S. Brunke, M.C. Costanzo, D.A. Fitzpatrick, P.W.J. de Groot, D. Harris, L.L. Hoyer, B. Hube, F.M. Klis, C. Kodira, N. Lennard, M.E. Logue, R. Martin, A.M. Neiman, E. Nikolaou, M.A. Quail, J. Quinn, M.C. Santos, F.F. Schmitzberger, G. Sherlock, P. Shah, K.A.T. Silverstein, M.S. Skrzypek, D. Soll, R. Staggs, I. Stansfield, M.P.H. Stumpf, P.E. Sudbery, T. Srikantha, Q. Zeng, J. Berman, M. Berriman, J. Heitman, N.A.R. Gow, M.C. Lorenz, B.W. Birren, M. Kellis, C.A. Cuomo, Evolution of pathogenicity and sexual reproduction in eight Candida genomes, Nature. 459 (2009) 657–662. https://doi.org/10.1038/nature08064.

108. T.J. Sharpton, J.E. Stajich, S.D. Rounsley, M.J. Gardner, J.R. Wortman, V.S. Jordar, R. Maiti, C.D. Kodira, D.E. Neafsey, Q. Zeng, C.-Y. Hung, C. McMahan, A. Muszewska, M. Grynberg, M.A. Mandel, E.M. Kellner, B.M. Barker, J.N. Galgiani, M.J. Orbach, T.N. Kirkland, G.T. Cole, M.R. Henn, B.W. Birren, J.W. Taylor, Comparative genomic analyses of the human fungal pathogens Coccidioides and their relatives, Genome Res. 19 (2009) 1722–1731. https://doi.org/10.1101/gr.087551.108.

109. A. B.J. Condon, Y. Leng, D. Wu, K.E. Bushley, R.A. Ohm, R. Otillar, J. Martin, W. Schackwitz, J. Grimwood, N. MohdZainudin, C. Xue, R. Wang, V.A. Manning, B. Dhillon, Z.J. Tu, B.J. Steffenson, Salamov, H. Sun, S. Lowry, K. LaButti, J. Han, A. Copeland, E. Lindquist, K. Barry, J. Schmutz, S.E. Baker, L.M. Ciuffetti, I.V. Grigoriev, S. Zhong, B.G. Turgeon, Comparative genome structure, secondary metabolite, and effector coding capacity across Cochliobolus pathogens, PLoS Genet. 9 (2013) e1003233. https://doi.org/10.1371/journal.pgen.1003233.

110. P. Gan, M. Narusaka, A. Tsushima, Y. Narusaka, Y. Takano, K. Shirasu, Draft Genome Assembly of Colletotrichum chlorophyti, a Pathogen of Herbaceous Plants, Genome Announc. 5 (2017). https://doi.org/10.1128/genomeA.01733-16.

111. R. Baroncelli, S. Sreenivasaprasad, S.A. Sukno, M.R. Thon, E. Holub, Draft Genome Sequence of Colletotrichum acutatum Sensu Lato (Colletotrichum fioriniae), Genome Announc. 2 (2014). https://doi.org/10.1128/genomeA.00112-14.

112. R.J. O’Connell, M.R. Thon, S. Hacquard, S.G. Amyotte, J. Kleemann, M.F. Torres, U. Damm, E.A. Buiate, L. Epstein, N. Alkan, J. Altmüller, L. Alvarado-Balderrama, C.A. Bauser, C. Becker, B.W. Birren, Z. Chen, J. Choi, J.A. Crouch, J.P. Duvick, M.A. Farman, P. Gan, D. Heiman, B. Henrissat, R.J. Howard, M. Kabbage, C. Koch, B. Kracher, Y. Kubo, A.D. Law, M.-H. Lebrun, Y.-H. Lee, I. Miyara, N. Moore, U. Neumann, K. Nordström, D.G. Panaccione, R. Panstruga, M. Place, R.H. Proctor, D. Prusky, G. Rech, R. Reinhardt, J.A. Rollins, S. Rounsley, C.L. Schardl, D.C. Schwartz, N. Shenoy, K. Shirasu, U.R. Sikhakolli, K. Stüber, S.A. Sukno, J.A. Sweigard, Y. Takano, H. Takahara, F. Trail, H.C. van der Does, L.M. Voll, I. Will, S. Young, Q. Zeng, J. Zhang, S. Zhou, M.B. Dickman, P. Schulze-Lefert, E. Ver Loren van Themaat, L.-J. Ma, L.J. Vaillancourt, Lifestyle transitions in plant pathogenic Colletotrichum fungi deciphered by genome and transcriptome analyses, Nat. Genet. 44 (2012) 1060–1065. https://doi.org/10.1038/ng.2372.

113. A. Zampounis, S. Pigné, J.-F. Dallery, A.H.J. Wittenberg, S. Zhou, D.C. Schwartz, M.R. Thon, R.J. O’Connell, Genome Sequence and Annotation of Colletotrichum higginsianum, a Causal Agent of Crucifer Anthracnose Disease, Genome Announc. 4 (2016). https://doi.org/10.1128/genomeA.00821-16.

114. P. Gan, M. Narusaka, N. Kumakura, A. Tsushima, Y. Takano, Y. Narusaka, K. Shirasu, Genus-Wide Comparative Genome Analyses of Colletotrichum Species Reveal Specific Gene Family Losses and Gains during Adaptation to Specific Infection Lifestyles, Genome Biol. Evol. 8 (2016) 1467– 1481. https://doi.org/10.1093/gbe/evw089.

115. R. Baroncelli, D.B. Amby, A. Zapparata, S. Sarrocco, G. Vannacci, G. Le Floch, R.J. Harrison, E. Holub, S.A. Sukno, S. Sreenivasaprasad, M.R. Thon, Gene family expansions and contractions are associated with host range in plant pathogens of the genus Colletotrichum, BMC Genomics. 17 (2016) 555. https://doi.org/10.1186/s12864-016-2917-6.

116. P. Gan, K. Ikeda, H. Irieda, M. Narusaka, R.J. O’Connell, Y. Narusaka, Y. Takano, Y. Kubo, K. Shirasu, Comparative genomic and transcriptomic analyses reveal the hemibiotrophic stage shift of Colletotrichum fungi, New Phytol. 197 (2013) 1236–1249. https://doi.org/10.1111/nph.12085.

117. R. Baroncelli, S.A. Sukno, S. Sarrocco, G. Cafà, G. Le Floch, M.R. Thon, Whole-Genome Sequence of the Orchid Anthracnose Pathogen Colletotrichum orchidophilum, Mol. Plant-Microbe Interact. MPMI. 31 (2018) 979–981. https://doi.org/10.1094/MPMI-03-18-0055-A.

118. S. Hacquard, B. Kracher, K. Hiruma, P.C. Münch, R. Garrido-Oter, M.R. Thon, A. Weimann, U. Damm, J.-F. Dallery, M. Hainaut, B. Henrissat, O. Lespinet, S. Sacristán, E. Ver Loren van Themaat, E. Kemen, A.C. McHardy, P. Schulze-Lefert, R.J. O’Connell, Survival trade-offs in plant roots during colonization by closely related beneficial and pathogenic fungi, Nat. Commun. 7 (2016) 11362. https://doi.org/10.1038/ncomms11362.

119. Y. Chang, S. Wang, S. Sekimoto, A.L. Aerts, C. Choi, A. Clum, K.M. LaButti, E.A. Lindquist, C. Yee Ngan, R.A. Ohm, A.A. Salamov, I.V. Grigoriev, J.W. Spatafora, M.L. Berbee, Phylogenomic Analyses Indicate that Early Fungi Evolved Digesting Cell Walls of Algal Ancestors of Land Plants, Genome Biol. Evol. 7 (2015) 1590–1601. https://doi.org/10.1093/gbe/evv090.

120. D.J. Jiménez, R.E. Hector, R. Riley, A. Lipzen, R.C. Kuo, M. Amirebrahimi, K.W. Barry, I.V. Grigoriev, J.D. van Elsas, N.N. Nichols, Draft Genome Sequence of Coniochaeta ligniaria NRRL 30616, a Lignocellulolytic Fungus for Bioabatement of Inhibitors in Plant Biomass Hydrolysates, Genome Announc. 5 (2017). https://doi.org/10.1128/genomeA.01476-16.

121. S.J. Mondo, D.J. Jiménez, R.E. Hector, A. Lipzen, M. Yan, K. LaButti, K. Barry, J.D. van Elsas, I.V. Grigoriev, N.N. Nichols, Genome expansion by allopolyploidization in the fungal strain Coniochaeta 2T2.1 and its exceptional lignocellulolytic machinery, Biotechnol. Biofuels. 12 (2019) 229. https://doi.org/10.1186/s13068-019-1569-6.

122. R. Castanera, G. Pérez, L. López-Varas, J. Amselem, K. LaButti, V. Singan, A. Lipzen, S. Haridas, K. Barry, I.V. Grigoriev, A.G. Pisabarro, L. Ramírez, Comparative genomics of Coniophora olivacea reveals different patterns of genome expansion in Boletales, BMC Genomics. 18 (2017) 883. https://doi.org/10.1186/s12864-017-4243-z.

123. T. Varga, K. Krizsán, C. Földi, B. Dima, M. Sánchez-García, S. Sánchez-Ramírez, G.J. Szöllősi, J.G. Szarkándi, V. Papp, L. Albert, W. Andreopoulos, C. Angelini, V. Antonín, K.W. Barry, N.L. Bougher, P. Buchanan, B. Buyck, V. Bense, P. Catcheside, M. Chovatia, J. Cooper, W. Dämon, D. Desjardin, P. Finy, J. Geml, S. Haridas, K. Hughes, A. Justo, D. Karasiński, I. Kautmanova, B. Kiss, S. Kocsubé, H. Kotiranta, K.M. LaButti, B.E. Lechner, K. Liimatainen, A. Lipzen, Z. Lukács, S. Mihaltcheva, L.N. Morgado, T. Niskanen, M.E. Noordeloos, R.A. Ohm, B. Ortiz-Santana, C. Ovrebo, N. Rácz, R. Riley, A. Savchenko, A. Shiryaev, K. Soop, V. Spirin, C. Szebenyi, M. Tomšovský, R.E. Tulloss, J. Uehling, I.V. Grigoriev, C. Vágvölgyi, T. Papp, F.M. Martin, O. Miettinen, D.S. Hibbett, L.G. Nagy, Megaphylogeny resolves global patterns of mushroom evolution, Nat. Ecol. Evol. 3 (2019) 668–678. https://doi.org/10.1038/s41559-019-0834-1.

124. J.E. Stajich, S.K. Wilke, D. Ahrén, C.H. Au, B.W. Birren, M. Borodovsky, C. Burns, B. Canbäck, L.A. Casselton, C.K. Cheng, J. Deng, F.S. Dietrich, D.C. Fargo, M.L. Farman, A.C. Gathman, J. Goldberg, R. Guigó, P.J. Hoegger, J.B. Hooker, A. Huggins, T.Y. James, T. Kamada, S. Kilaru, C. Kodira, U. Kües, D. Kupfer, H.S. Kwan, A. Lomsadze, W. Li, W.W. Lilly, L.-J. Ma, A.J. Mackey, G. Manning, F. Martin, H. Muraguchi, D.O. Natvig, H. Palmerini, M.A. Ramesh, C.J. Rehmeyer, B.A. Roe, N. Shenoy, M. Stanke, V. Ter-Hovhannisyan, A. Tunlid, R. Velagapudi, T.J. Vision, Q. Zeng, M.E. Zolan, P.J. Pukkila, Insights into evolution of multicellular fungi from the assembled chromosomes of the mushroom Coprinopsis cinerea (Coprinus cinereus), Proc. Natl. Acad. Sci. U. S. A. 107 (2010) 11889–11894. https://doi.org/10.1073/pnas.1003391107.

125. H. Muraguchi, K. Umezawa, M. Niikura, M. Yoshida, T. Kozaki, K. Ishii, K. Sakai, M. Shimizu, K. Nakahori, Y. Sakamoto, C. Choi, C.Y. Ngan, E. Lindquist, A. Lipzen, A. Tritt, S. Haridas, K. Barry, I.V. Grigoriev, P.J. Pukkila, Strand-Specific RNA-Seq Analyses of Fruiting Body Development in Coprinopsis cinerea, PloS One. 10 (2015) e0141586. https://doi.org/10.1371/journal.pone.0141586.

126. P. Zheng, Y. Xia, G. Xiao, C. Xiong, X. Hu, S. Zhang, H. Zheng, Y. Huang, Y. Zhou, S. Wang, G.-P. Zhao, X. Liu, R.J. St Leger, C. Wang, Genome sequence of the insect pathogenic fungus Cordyceps militaris, a valued traditional Chinese medicine, Genome Biol. 12 (2011) R116. https://doi.org/10.1186/gb-2011-12-11-r116.

127. D. Lopez, S. Ribeiro, P. Label, B. Fumanal, J.-S. Venisse, A. Kohler, R.R. de Oliveira, K. Labutti, A. Lipzen, K. Lail, D. Bauer, R.A. Ohm, K.W. Barry, J. Spatafora, I.V. Grigoriev, F.M. Martin, V. Pujade-Renaud, Genome-Wide Analysis of Corynespora cassiicola Leaf Fall Disease Putative Effectors, Front. Microbiol. 9 (2018) 276. https://doi.org/10.3389/fmicb.2018.00276.

128. A.L. Pendleton, K.E. Smith, N. Feau, F.M. Martin, I.V. Grigoriev, R. Hamelin, C.D. Nelson, J.G. Burleigh, J.M. Davis, Duplications and losses in gene families of rust pathogens highlight putative effectors, Front. Plant Sci. 5 (2014) 299. https://doi.org/10.3389/fpls.2014.00299.

129. J.A. Crouch, A. Dawe, A. Aerts, K. Barry, A.C.L. Churchill, J. Grimwood, B.I. Hillman, M.G. Milgroom, J. Pangilinan, M. Smith, A. Salamov, J. Schmutz, J.S. Yadav, I.V. Grigoriev, D.L. Nuss, Genome Sequence of the Chestnut Blight Fungus Cryphonectria parasitica EP155: A Fundamental Resource for an Archetypical Invasive Plant Pathogen, Phytopathology. 110 (2020) 1180–1188. https://doi.org/10.1094/PHYTO-12-19-0478-A.

130. D. Close, J. Ojumu, Draft Genome Sequence of the Oleaginous Yeast Cryptococcus curvatus ATCC 20509, Genome Announc. 4 (2016). https://doi.org/10.1128/genomeA.01235-16.

131. B.J. Loftus, E. Fung, P. Roncaglia, D. Rowley, P. Amedeo, D. Bruno, J. Vamathevan, M. Miranda, I.J. Anderson, J.A. Fraser, J.E. Allen, I.E. Bosdet, M.R. Brent, R. Chiu, T.L. Doering, M.J. Donlin, C.A. D’Souza, D.S. Fox, V. Grinberg, J. Fu, M. Fukushima, B.J. Haas, J.C. Huang, G. Janbon, S.J.M. Jones, H.L. Koo, M.I. Krzywinski, J.K. Kwon-Chung, K.B. Lengeler, R. Maiti, M.A. Marra, R.E. Marra, C.A. Mathewson, T.G. Mitchell, M. Pertea, F.R. Riggs, S.L. Salzberg, J.E. Schein, A. Shvartsbeyn, H. Shin, M. Shumway, C.A. Specht, B.B. Suh, A. Tenney, T.R. Utterback, B.L. Wickes, J.R. Wortman, N.H. Wye, J.W. Kronstad, J.K. Lodge, J. Heitman, R.W. Davis, C.M. Fraser, R.W. Hyman, The genome of the basidiomycetous yeast and human pathogen Cryptococcus neoformans, Science. 307 (2005) 1321–1324. https://doi.org/10.1126/science.1103773.

132. G. Janbon, K.L. Ormerod, D. Paulet, E.J. Byrnes, V. Yadav, G. Chatterjee, N. Mullapudi, C.-C. Hon, R.B. Billmyre, F. Brunel, Y.-S. Bahn, W. Chen, Y. Chen, E.W.L. Chow, J.-Y. Coppée, A. Floyd-Averette, C. Gaillardin, K.J. Gerik, J. Goldberg, S. Gonzalez-Hilarion, S. Gujja, J.L. Hamlin, Y.-P. Hsueh, G. Ianiri, S. Jones, C.D. Kodira, L. Kozubowski, W. Lam, M. Marra, L.D. Mesner, P.A. Mieczkowski, F. Moyrand, K. Nielsen, C. Proux, T. Rossignol, J.E. Schein, S. Sun, C. Wollschlaeger, I.A. Wood, Q. Zeng, C. Neuvéglise, C.S. Newlon, J.R. Perfect, J.K. Lodge, A. Idnurm, J.E. Stajich, J.W. Kronstad, K. Sanyal, J. Heitman, J.A. Fraser, C.A. Cuomo, F.S. Dietrich, Analysis of the genome and transcriptome of Cryptococcus neoformans var. grubii reveals complex RNA expression and microevolution leading to virulence attenuation, PLoS Genet. 10 (2014) e1004261. https://doi.org/10.1371/journal.pgen.1004261.

133. D. Floudas, B.W. Held, R. Riley, L.G. Nagy, G. Koehler, A.S. Ransdell, H. Younus, J. Chow, J. Chiniquy, A. Lipzen, A. Tritt, H. Sun, S. Haridas, K. LaButti, R.A. Ohm, U. Kües, R.A. Blanchette, I.V. Grigoriev, R.E. Minto, D.S. Hibbett, Evolution of novel wood decay mechanisms in Agaricales revealed by the genome sequences of Fistulina hepatica and Cylindrobasidium torrendii, Fungal Genet. Biol. FG B. 76 (2015) 78–92. https://doi.org/10.1016/j.fgb.2015.02.002.

134. S. Camiolo, M. Toome-Heller, M.C. Aime, S. Haridas, I.V. Grigoriev, A. Porceddu, I. Mannazzu, An analysis of codon bias in six red yeast species, Yeast Chichester Engl. 36 (2019) 53–64. https://doi.org/10.1002/yea.3359.

135. L.G. Nagy, R. Riley, A. Tritt, C. Adam, C. Daum, D. Floudas, H. Sun, J.S. Yadav, J. Pangilinan, K.-H. Larsson, K. Matsuura, K. Barry, K. Labutti, R. Kuo, R.A. Ohm, S.S. Bhattacharya, T. Shirouzu, Y. Yoshinaga, F.M. Martin, I.V. Grigoriev, D.S. Hibbett, Comparative Genomics of Early-Diverging Mushroom-Forming Fungi Provides Insights into the Origins of Lignocellulose Decay Capabilities, Mol. Biol. Evol. 33 (2016) 959–970. https://doi.org/10.1093/molbev/msv337.

136. W. Wu, R.W. Davis, M.B. Tran-Gyamfi, A. Kuo, K. LaButti, S. Mihaltcheva, H. Hundley, M. Chovatia, E. Lindquist, K. Barry, I.V. Grigoriev, B. Henrissat, J.M. Gladden, Characterization of four endophytic fungi as potential consolidated bioprocessing hosts for conversion of lignocellulose into advanced biofuels, Appl. Microbiol. Biotechnol. 101 (2017) 2603–2618. https://doi.org/10.1007/s00253-017-8091-1.

137. C. Sacerdot, S. Casaregola, I. Lafontaine, F. Tekaia, B. Dujon, O. Ozier-Kalogeropoulos, Promiscuous DNA in the nuclear genomes of hemiascomycetous yeasts, FEMS Yeast Res. 8 (2008) 846–857. https://doi.org/10.1111/j.1567-1364.2008.00409.x.

138. J. Piškur, Z. Ling, M. Marcet-Houben, O.P. Ishchuk, A. Aerts, K. LaButti, A. Copeland, E. Lindquist, K. Barry, C. Compagno, L. Bisson, I.V. Grigoriev, T. Gabaldón, T. Phister, The genome of wine yeast Dekkera bruxellensis provides a tool to explore its food-related properties, Int. J. Food Microbiol. 157 (2012) 202–209. https://doi.org/10.1016/j.ijfoodmicro.2012.05.008.

139. H. Park, B. Min, Y. Jang, J. Kim, A. Lipzen, A. Sharma, B. Andreopoulos, J. Johnson, R. Riley, J.W. Spatafora, B. Henrissat, K.H. Kim, I.V. Grigoriev, J.-J. Kim, I.-G. Choi, Comprehensive genomic and transcriptomic analysis of polycyclic aromatic hydrocarbon degradation by a mycoremediation fungus, Dentipellis sp. KUC8613, Appl. Microbiol. Biotechnol. 103 (2019) >8145–8155. https://doi.org/10.1007/s00253-019-10089-6.

140. S. Casado López, M. Peng, P. Daly, B. Andreopoulos, J. Pangilinan, A. Lipzen, R. Riley, S. Ahrendt, V. Ng, K. Barry, C. Daum, I.V. Grigoriev, K.S. Hildén, M.R. Mäkelä, R.P. de Vries, Draft Genome Sequences of Three Monokaryotic Isolates of the White-Rot Basidiomycete Fungus Dichomitus squalens, Microbiol. Resour. Announc. 8 (2019). https://doi.org/10.1128/MRA.00264-19.

141. E. Peyretaillade, O. Gonçalves, S. Terrat, E. Dugat-Bony, P. Wincker, R.S. Cornman, J.D. Evans, F. Delbac, P. Peyret, Identification of transcriptional signals in Encephalitozoon cuniculi widespread among Microsporidia phylum: support for accurate structural genome annotation, BMC Genomics. 10 (2009) 607. https://doi.org/10.1186/1471-2164-10-607.

142. J.-F. Pombert, M. Selman, F. Burki, F.T. Bardell, L. Farinelli, L.F. Solter, D.W. Whitman, L.M. Weiss, N. Corradi, P.J. Keeling, Gain and loss of multiple functionally related, horizontally transferred genes in the reduced genomes of two microsporidian parasites, Proc. Natl. Acad. Sci. U. S. A. 109 (2012) 12638–12643. https://doi.org/10.1073/pnas.1205020109.

143. N. Corradi, J.-F. Pombert, L. Farinelli, E.S. Didier, P.J. Keeling, The complete sequence of the smallest known nuclear genome from the microsporidian Encephalitozoon intestinalis, Nat. Commun. 1 (2010) 77. https://doi.org/10.1038/ncomms1082.

144. Y.-Y. Wang, B. Liu, X.-Y. Zhang, Q.-M. Zhou, T. Zhang, H. Li, Y.-F. Yu, X.-L. Zhang, X.-Y. Hao, M. Wang, L. Wang, J.-C. Wei, Genome characteristics reveal the impact of lichenization on lichen-forming fungus Endocarpon pusillum Hedwig (Verrucariales, Ascomycota), BMC Genomics. 15 (2014) 34. https://doi.org/10.1186/1471-2164-15-34.

145. Y. Chang, A. Desirò, H. Na, L. Sandor, A. Lipzen, A. Clum, K. Barry, I.V. Grigoriev, F.M. Martin, J.E. Stajich, M.E. Smith, G. Bonito, J.W. Spatafora, Phylogenomics of Endogonaceae and evolution of mycorrhizas within Mucoromycota, New Phytol. 222 (2019) 511–525. https://doi.org/10.1111/nph.15613.

146. D.E. Akiyoshi, H.G. Morrison, S. Lei, X. Feng, Q. Zhang, N. Corradi, H. Mayanja, J.K. Tumwine, P.J. Keeling, L.M. Weiss, S. Tzipori, Genomic survey of the non-cultivatable opportunistic human pathogen, Enterocytozoon bieneusi, PLoS Pathog. 5 (2009) e1000261. https://doi.org/10.1371/journal.ppat.1000261.

147. F.S. Dietrich, S. Voegeli, S. Brachat, A. Lerch, K. Gates, S. Steiner, C. Mohr, R. Pöhlmann, P. Luedi, S. Choi, R.A. Wing, A. Flavier, T.D. Gaffney, P. Philippsen, The Ashbya gossypii genome as a tool for mapping the ancient Saccharomyces cerevisiae genome, Science. 304 (2004) 304– 307. https://doi.org/10.1126/science.1095781.

148. L. Jones, S. Riaz, A. Morales-Cruz, K.C.H. Amrine, B. McGuire, W.D. Gubler, M.A. Walker, D. Cantu, Adaptive genomic structural variation in the grape powdery mildew pathogen, Erysiphe necator, BMC Genomics. 15 (2014) 1081. https://doi.org/10.1186/1471-2164-15-1081.

149. T. Kis-Papo, A.R. Weig, R. Riley, D. Peršoh, A. Salamov, H. Sun, A. Lipzen, S.P. Wasser, G. Rambold, I.V. Grigoriev, E. Nevo, Genomic adaptations of the halophilic Dead Sea filamentous fungus Eurotium rubrum, Nat. Commun. 5 (2014) 3745. https://doi.org/10.1038/ncomms4745.

150. Z. Chen, D.A. Martinez, S. Gujja, S.M. Sykes, Q. Zeng, P.J. Szaniszlo, Z. Wang, C.A. Cuomo, Comparative genomic and transcriptomic analysis of wangiella dermatitidis, a major cause of phaeohyphomycosis and a model black yeast human pathogen, G3 Bethesda Md. 4 (2014) 561– 578. https://doi.org/10.1534/g3.113.009241.

151. J.D. Tang, A.D. Perkins, T.S. Sonstegard, S.G. Schroeder, S.C. Burgess, S.V. Diehl, Short-read sequencing for genomic analysis of the brown rot fungus Fibroporia radiculosa, Appl. Environ. Microbiol. 78 (2012) 2272–2281. https://doi.org/10.1128/AEM.06745-11.

152. A. Bombassaro, S. de Hoog, V.A. Weiss, E.M. Souza, A.C.R. Leão, F.F. Costa, V. Baura, M.Z. Tadra-Sfeir, E. Balsanelli, L.F. Moreno, R.T. Raittz, M.B.R. Steffens, F.O. Pedrosa, J. Sun, L. Xi, A.L. Bocca, M.S. Felipe, M. Teixeira, G.D. Santos, F.Q. Telles Filho, C.M.P.S. Azevedo, R.R. Gomes, V.A. Vicente, Draft Genome Sequence of Fonsecaea monophora Strain CBS 269.37, an Agent of Human Chromoblastomycosis, Genome Announc. 4 (2016). https://doi.org/10.1128/genomeA.00731-16.

153. F.F. Costa, S. de Hoog, R.T. Raittz, V.A. Weiss, A.C.R. Leão, A. Bombassaro, J. Sun, L.F. Moreno, E.M. Souza, F.O. Pedrosa, M.B.R. Steffens, V. Baura, M.Z. Tadra-Sfeir, E. Balsanelli, M.J. Najafzadeh, R.R. Gomes, M.S. Felipe, M. Teixeira, G.D. Santos, L. Xi, M.A. Alves de Castro, V.A. Vicente, Draft Genome Sequence of Fonsecaea nubica Strain CBS 269.64, Causative Agent of Human Chromoblastomycosis, Genome Announc. 4 (2016). https://doi.org/10.1128/genomeA.00735-16.

154. P. Wiemann, C.M.K. Sieber, K.W. von Bargen, L. Studt, E.-M. Niehaus, J.J. Espino, K. Huß, C.B. Michielse, S. Albermann, D. Wagner, S.V. Bergner, L.R. Connolly, A. Fischer, G. Reuter, K. Kleigrewe, T. Bald, B.D. Wingfield, R. Ophir, S. Freeman, M. Hippler, K.M. Smith, D.W. Brown, R.H. Proctor, M. Münsterkötter, M. Freitag, H.-U. Humpf, U. Güldener, B. Tudzynski, Deciphering the cryptic genome: genome-wide analyses of the rice pathogen Fusarium fujikuroi reveal complex regulation of secondary metabolism and novel metabolites, PLoS Pathog. 9 (2013) e1003475. https://doi.org/10.1371/journal.ppat.1003475.

155. C.A. Cuomo, U. Güldener, J.-R. Xu, F. Trail, B.G. Turgeon, A. Di Pietro, J.D. Walton, L.-J. Ma, S.E. Baker, M. Rep, G. Adam, J. Antoniw, T. Baldwin, S. Calvo, Y.-L. Chang, D. Decaprio, L.R. Gale, S. Gnerre, R.S. Goswami, K. Hammond-Kosack, L.J. Harris, K. Hilburn, J.C. Kennell, S. Kroken, J.K. Magnuson, G. Mannhaupt, E. Mauceli, H.-W. Mewes, R. Mitterbauer, G. Muehlbauer, M. Münsterkötter, D. Nelson, K. O’donnell, T. Ouellet, W. Qi, H. Quesneville, M.I.G. Roncero, K.-Y. Seong, I.V. Tetko, M. Urban, C. Waalwijk, T.J. Ward, J. Yao, B.W. Birren, H.C. Kistler, The Fusarium graminearum genome reveals a link between localized polymorphism and pathogen specialization, Science. 317 (2007) 1400–1402. https://doi.org/10.1126/science.1143708.

156. G.A. DeIulio, L. Guo, Y. Zhang, J.M. Goldberg, H.C. Kistler, L.-J. Ma, Kinome Expansion in the Fusarium oxysporum Species Complex Driven by Accessory Chromosomes, MSphere. 3 (2018). https://doi.org/10.1128/mSphere.00231-18.

157. L.-J. Ma, H.C. van der Does, K.A. Borkovich, J.J. Coleman, M.-J. Daboussi, A. Di Pietro, M. Dufresne, M. Freitag, M. Grabherr, B. Henrissat, P.M. Houterman, S. Kang, W.-B. Shim, C. Woloshuk, X. Xie, J.-R. Xu, J. Antoniw, S.E. Baker, B.H. Bluhm, A. Breakspear, D.W. Brown, R.A.E. Butchko, S. Chapman, R. Coulson, P.M. Coutinho, E.G.J. Danchin, A. Diener, L.R. Gale, D.M. Gardiner, S. Goff, K.E. Hammond-Kosack, K. Hilburn, A. Hua-Van, W. Jonkers, K. Kazan, C.D. Kodira, M. Koehrsen, L. Kumar, Y.-H. Lee, L. Li, J.M. Manners, D. Miranda-Saavedra, M. Mukherjee, G. Park, J. Park, S.-Y. Park, R.H. Proctor, A. Regev, M.C. Ruiz-Roldan, D. Sain, S. Sakthikumar, S. Sykes, D.C. Schwartz, B.G. Turgeon, I. Wapinski, O. Yoder, S. Young, Q. Zeng, S. Zhou, J. Galagan, C.A. Cuomo, H.C. Kistler, M. Rep, Comparative genomics reveals mobile pathogenicity chromosomes in Fusarium, Nature. 464 (2010) 367–373. https://doi.org/10.1038/nature08850.

158. L.-J. Ma, T. Shea, S. Young, Q. Zeng, H.C. Kistler, Genome Sequence of Fusarium oxysporum f. sp. melonis Strain NRRL 26406, a Fungus Causing Wilt Disease on Melon, Genome Announc. 2 (2014). https://doi.org/10.1128/genomeA.00730-14.

159. A.H. Williams, M. Sharma, L.F. Thatcher, S. Azam, J.K. Hane, J. Sperschneider, B.N. Kidd, J.P. Anderson, R. Ghosh, G. Garg, J. Lichtenzveig, H.C. Kistler, T. Shea, S. Young, S.-A.G. Buck, L.G. Kamphuis, R. Saxena, S. Pande, L.-J. Ma, R.K. Varshney, K.B. Singh, Comparative genomics and prediction of conditionally dispensable sequences in legume-infecting Fusarium oxysporum formae speciales facilitates identification of candidate effectors, BMC Genomics. 17 (2016) 191. https://doi.org/10.1186/s12864-016-2486-8.

160. D.M. Gardiner, M.C. McDonald, L. Covarelli, P.S. Solomon, A.G. Rusu, M. Marshall, K. Kazan, S. Chakraborty, B.A. McDonald, J.M. Manners, Comparative pathogenomics reveals horizontally acquired novel virulence genes in fungi infecting cereal hosts, PLoS Pathog. 8 (2012) e1002952. https://doi.org/10.1371/journal.ppat.1002952.

161. L.H. Okagaki, C.C. Nunes, J. Sailsbery, B. Clay, D. Brown, T. John, Y. Oh, N. Young, M. Fitzgerald, B.J. Haas, Q. Zeng, S. Young, X. Adiconis, L. Fan, J.Z. Levin, T.K. Mitchell, P.A. Okubara, M.L. Farman, L.M. Kohn, B. Birren, L.-J. Ma, R.A. Dean, Genome Sequences of Three Phytopathogenic Species of the Magnaporthaceae Family of Fungi, G3 Bethesda Md. 5 (2015) 2539–2545. https://doi.org/10.1534/g3.115.020057.

162. L. Chen, Q. Yue, X. Zhang, M. Xiang, C. Wang, S. Li, Y. Che, F.J. Ortiz-López, G.F. Bills, X. Liu, Z. An, Genomics-driven discovery of the pneumocandin biosynthetic gene cluster in the fungus Glarea lozoyensis, BMC Genomics. 14 (2013) 339. https://doi.org/10.1186/1471-2164-14-339.

163. S. DiGuistini, Y. Wang, N.Y. Liao, G. Taylor, P. Tanguay, N. Feau, B. Henrissat, S.K. Chan, U. Hesse-Orce, S.M. Alamouti, C.K.M. Tsui, R.T. Docking, A. Levasseur, S. Haridas, G. Robertson, I. Birol, R.A. Holt, M.A. Marra, R.C. Hamelin, M. Hirst, S.J.M. Jones, J. Bohlmann, C. Breuil, Genome and transcriptome analyses of the mountain pine beetle-fungal symbiont Grosmannia clavigera, a lodgepole pine pathogen, Proc. Natl. Acad. Sci. U. S. A. 108 (2011) 2504–2509. https://doi.org/10.1073/pnas.1011289108.

164. F. Barbi, A. Kohler, K. Barry, P. Baskaran, C. Daum, L. Fauchery, K. Ihrmark, A. Kuo, K. LaButti, A. Lipzen, E. Morin, I.V. Grigoriev, B. Henrissat, B.D. Lindahl, F. Martin, Fungal ecological strategies reflected in gene transcription - a case study of two litter decomposers, Environ. Microbiol. 22 (2020) 1089–1103. https://doi.org/10.1111/1462-2920.14873.

165. A. Olson, A. Aerts, F. Asiegbu, L. Belbahri, O. Bouzid, A. Broberg, B. Canbäck, P.M. Coutinho, D. Cullen, K. Dalman, G. Deflorio, L.T.A. van Diepen, C. Dunand, S. Duplessis, M. Durling, P. Gonthier, J. Grimwood, C.G. Fossdal, D. Hansson, B. Henrissat, A. Hietala, K. Himmelstrand, D. Hoffmeister, N. Högberg, T.Y. James, M. Karlsson, A. Kohler, U. Kües, Y.-H. Lee, Y.-C. Lin, M. Lind, E. Lindquist, V. Lombard, S. Lucas, K. Lundén, E. Morin, C. Murat, J. Park, T. Raffaello, P. Rouzé, A. Salamov, J. Schmutz, H. Solheim, J. Ståhlberg, H. Vélëz, R.P. de Vries, A. Wiebenga, S. Woodward, I. Yakovlev, M. Garbelotto, F. Martin, I.V. Grigoriev, J. Stenlid, Insight into trade-off between wood decay and parasitism from the genome of a fungal forest pathogen, New Phytol. 194 (2012) 1001–1013. https://doi.org/10.1111/j.1469-8137.2012.04128.x.

166. S. Joneson, J.E. Stajich, S.-H. Shiu, E.B. Rosenblum, Genomic transition to pathogenicity in chytrid fungi, PLoS Pathog. 7 (2011) e1002338. https://doi.org/10.1371/journal.ppat.1002338.

167. M. Lenassi, C. Gostinčar, S. Jackman, M. Turk, I. Sadowski, C. Nislow, S. Jones, I. Birol, N.G. Cimerman, A. Plemenitaš, Whole genome duplication and enrichment of metal cation transporters revealed by de novo genome sequencing of extremely halotolerant black yeast Hortaea werneckii, PloS One. 8 (2013) e71328. https://doi.org/10.1371/journal.pone.0071328.

168. H.-L. Liao, G. Bonito, J.A. Rojas, K. Hameed, S. Wu, C.W. Schadt, J. Labbé, G.A. Tuskan, F. Martin, I.V. Grigoriev, R. Vilgalys, Fungal Endophytes of Populus trichocarpa Alter Host Phenotype, Gene Expression, and Rhizobiome Composition, Mol. Plant-Microbe Interact. MPMI. 32 (2019) 853–864. https://doi.org/10.1094/MPMI-05-18-0133-R.

169. J.L. Gordon, D. Armisén, E. Proux-Wéra, S.S. ÓhÉigeartaigh, K.P. Byrne, K.H. Wolfe, Evolutionary erosion of yeast sex chromosomes by mating-type switching accidents, Proc. Natl. Acad. Sci. U. S. A. 108 (2011) 20024–20029. https://doi.org/10.1073/pnas.1112808108.

170. B. Dujon, D. Sherman, G. Fischer, P. Durrens, S. Casaregola, I. Lafontaine, J. De Montigny, C. Marck, C. Neuvéglise, E. Talla, N. Goffard, L. Frangeul, M. Aigle, V. Anthouard, A. Babour, V. Barbe, S. Barnay, S. Blanchin, J.-M. Beckerich, E. Beyne, C. Bleykasten, A. Boisramé, J. Boyer, L. Cattolico, F. Confanioleri, A. De Daruvar, L. Despons, E. Fabre, C. Fairhead, H. Ferry-Dumazet, A. Groppi, F. Hantraye, C. Hennequin, N. Jauniaux, P. Joyet, R. Kachouri, A. Kerrest, R. Koszul, M. Lemaire, I. Lesur, L. Ma, H. Muller, J.-M. Nicaud, M. Nikolski, S. Oztas, O. Ozier-Kalogeropoulos, S. Pellenz, S. Potier, G.-F. Richard, M.-L. Straub, A. Suleau, D. Swennen, F. Tekaia, M. Wésolowski-Louvel, E. Westhof, B. Wirth, M. Zeniou-Meyer, I. Zivanovic, M. Bolotin-Fukuhara, A. Thierry, C. Bouchier, B. Caudron, C. Scarpelli, C. Gaillardin, J. Weissenbach, P. Wincker, J.-L. Souciet, Genome evolution in yeasts, Nature. 430 (2004) 35–44. https://doi.org/10.1038/nature02579.

171. L. Morales, B. Noel, B. Porcel, M. Marcet-Houben, M.-F. Hullo, C. Sacerdot, F. Tekaia, V. Leh-Louis, L. Despons, V. Khanna, J.-M. Aury, V. Barbe, A. Couloux, K. Labadie, E. Pelletier, J.-L. Souciet, T. Boekhout, T. Gabaldon, P. Wincker, B. Dujon, Complete DNA sequence of Kuraishia capsulata illustrates novel genomic features among budding yeasts (Saccharomycotina), Genome Biol. Evol. 5 (2013) 2524–2539. https://doi.org/10.1093/gbe/evt201.

172. F. Martin, A. Aerts, D. Ahrén, A. Brun, E.G.J. Danchin, F. Duchaussoy, J. Gibon, A. Kohler, E. Lindquist, V. Pereda, A. Salamov, H.J. Shapiro, J. Wuyts, D. Blaudez, M. Buée, P. Brokstein, B. Canbäck, D. Cohen, P.E. Courty, P.M. Coutinho, C. Delaruelle, J.C. Detter, A. Deveau, S. DiFazio, S. Duplessis, L. Fraissinet-Tachet, E. Lucic, P. Frey-Klett, C. Fourrey, I. Feussner, G. Gay, J. Grimwood, P.J. Hoegger, P. Jain, S. Kilaru, J. Labbé, Y.C. Lin, V. Legué, F. Le Tacon, R. Marmeisse, D. Melayah, B. Montanini, M. Muratet, U. Nehls, H. Niculita-Hirzel, M.P. Oudot-Le Secq, M. Peter, H. Quesneville, B. Rajashekar, M. Reich, N. Rouhier, J. Schmutz, T. Yin, M. Chalot, B. Henrissat, U. Kües, S. Lucas, Y. Van de Peer, G.K. Podila, A. Polle, P.J. Pukkila, P.M. Richardson, P. Rouzé, I.R. Sanders, J.E. Stajich, A. Tunlid, G. Tuskan, I.V. Grigoriev, The genome of Laccaria bicolor provides insights into mycorrhizal symbiosis, Nature. 452 (2008) 88–92. https://doi.org/10.1038/nature06556.

173. S.-G. Park, S.I. Yoo, D.S. Ryu, H. Lee, Y.J. Ahn, H. Ryu, J. Ko, C.P. Hong, Long-read transcriptome data for improved gene prediction in Lentinula edodes, Data Brief. 15 (2017) 454–458. https://doi.org/10.1016/j.dib.2017.09.052.

174. L. Chen, Y. Gong, Y. Cai, W. Liu, Y. Zhou, Y. Xiao, Z. Xu, Y. Liu, X. Lei, G. Wang, M. Guo, X. Ma, Y. Bian, Genome Sequence of the Edible Cultivated Mushroom Lentinula edodes (Shiitake) Reveals Insights into Lignocellulose Degradation, PloS One. 11 (2016) e0160336. https://doi.org/10.1371/journal.pone.0160336.

175. B. Wu, Z. Xu, A. Knudson, A. Carlson, N. Chen, S. Kovaka, K. LaButti, A. Lipzen, C. Pennachio, R. Riley, W. Schakwitz, K. Umezawa, R.A. Ohm, I.V. Grigoriev, L.G. Nagy, J. Gibbons, D. Hibbett, Genomics and Development of Lentinus tigrinus: A White-Rot Wood-Decaying Mushroom with Dimorphic Fruiting Bodies, Genome Biol. Evol. 10 (2018) 3250–3261. https://doi.org/10.1093/gbe/evy246.

176. T. Rouxel, J. Grandaubert, J.K. Hane, C. Hoede, A.P. van de Wouw, A. Couloux, V. Dominguez, V. Anthouard, P. Bally, S. Bourras, A.J. Cozijnsen, L.M. Ciuffetti, A. Degrave, A. Dilmaghani, L. Duret, I. Fudal, S.B. Goodwin, L. Gout, N. Glaser, J. Linglin, G.H.J. Kema, N. Lapalu, C.B. Lawrence, K. May, M. Meyer, B. Ollivier, J. Poulain, C.L. Schoch, A. Simon, J.W. Spatafora, A. Stachowiak, B.G. Turgeon, B.M. Tyler, D. Vincent, J. Weissenbach, J. Amselem, H. Quesneville, R.P. Oliver, P. Wincker, M.-H. Balesdent, B.J. Howlett, Effector diversification within compartments of the Leptosphaeria maculans genome affected by Repeat-Induced Point mutations, Nat. Commun. 2 (2011) 202. https://doi.org/10.1038/ncomms1189.

177. F.O. Aylward, K.E. Burnum-Johnson, S.G. Tringe, C. Teiling, D.M. Tremmel, J.A. Moeller, J.J. Scott, K.W. Barry, P.D. Piehowski, C.D. Nicora, S.A. Malfatti, M.E. Monroe, S.O. Purvine, L.A. Goodwin, R.D. Smith, G.M. Weinstock, N.M. Gerardo, G. Suen, M.S. Lipton, C.R. Currie, Leucoagaricus gongylophorus produces diverse enzymes for the degradation of recalcitrant plant polymers in leaf-cutter ant fungus gardens, Appl. Environ. Microbiol. 79 (2013) 3770– 3778. https://doi.org/10.1128/AEM.03833-12.

178. S.J. Mondo, R.O. Dannebaum, R.C. Kuo, K.B. Louie, A.J. Bewick, K. LaButti, S. Haridas, A. Kuo, A. Salamov, S.R. Ahrendt, R. Lau, B.P. Bowen, A. Lipzen, W. Sullivan, B.B. Andreopoulos, A. Clum, E. Lindquist, C. Daum, T.R. Northen, G. Kunde-Ramamoorthy, R.J. Schmitz, A. Gryganskyi, D. Culley, J. Magnuson, T.Y. James, M.A. O’Malley, J.E. Stajich, J.W. Spatafora, A. Visel, I.V. Grigoriev, Widespread adenine N6-methylation of active genes in fungi, Nat. Genet. 49 (2017) 964–968. https://doi.org/10.1038/ng.3859.

179. V.U. Schwartze, S. Winter, E. Shelest, M. Marcet-Houben, F. Horn, S. Wehner, J. Linde, V. Valiante, M. Sammeth, K. Riege, M. Nowrousian, K. Kaerger, I.D. Jacobsen, M. Marz, A.A. Brakhage, T. Gabaldón, S. Böcker, K. Voigt, Gene expansion shapes genome architecture in the human pathogen Lichtheimia corymbifera: an evolutionary genomics analysis in the ancient terrestrial mucorales (Mucoromycotina), PLoS Genet. 10 (2014) e1004496. https://doi.org/10.1371/journal.pgen.1004496.

180. M.S. Islam, M.S. Haque, M.M. Islam, E.M. Emdad, A. Halim, Q.M.M. Hossen, M.Z. Hossain, B. Ahmed, S. Rahim, M.S. Rahman, M.M. Alam, S. Hou, X. Wan, J.A. Saito, M. Alam, Tools to kill: genome of one of the most destructive plant pathogenic fungi Macrophomina phaseolina, BMC Genomics. 13 (2012) 493. https://doi.org/10.1186/1471-2164-13-493.

181. R.A. Dean, N.J. Talbot, D.J. Ebbole, M.L. Farman, T.K. Mitchell, M.J. Orbach, M. Thon, R. Kulkarni, J.-R. Xu, H. Pan, N.D. Read, Y.-H. Lee, I. Carbone, D. Brown, Y.Y. Oh, N. Donofrio, J.S. Jeong, D.M. Soanes, S. Djonovic, E. Kolomiets, C. Rehmeyer, W. Li, M. Harding, S. Kim, M.-H. Lebrun, H. Bohnert, S. Coughlan, J. Butler, S. Calvo, L.-J. Ma, R. Nicol, S. Purcell, C. Nusbaum, J.E. Galagan, B.W. Birren, The genome sequence of the rice blast fungus Magnaporthe grisea, Nature. 434 (2005) 980–986. https://doi.org/10.1038/nature03449.

182. J. Xu, C.W. Saunders, P. Hu, R.A. Grant, T. Boekhout, E.E. Kuramae, J.W. Kronstad, Y.M. Deangelis, N.L. Reeder, K.R. Johnstone, M. Leland, A.M. Fieno, W.M. Begley, Y. Sun, M.P. Lacey, T. Chaudhary, T. Keough, L. Chu, R. Sears, B. Yuan, T.L. Dawson, Dandruff-associated Malassezia genomes reveal convergent and divergent virulence traits shared with plant and human fungal pathogens, Proc. Natl. Acad. Sci. U. S. A. 104 (2007) 18730–18735. https://doi.org/10.1073/pnas.0706756104.

183. A. Gioti, B. Nystedt, W. Li, J. Xu, A. Andersson, A.F. Averette, K. Münch, X. Wang, C. Kappauf, J.M. Kingsbury, B. Kraak, L.A. Walker, H.J. Johansson, T. Holm, J. Lehtiö, J.E. Stajich, P. Mieczkowski, R. Kahmann, J.C. Kennell, M.E. Cardenas, J. Lundeberg, C.W. Saunders, T. Boekhout, T.L. Dawson, C.A. Munro, P.W.J. de Groot, G. Butler, J. Heitman, A. Scheynius, Genomic insights into the atopic eczema-associated skin commensal yeast Malassezia sympodialis, MBio. 4 (2013) e00572–00512. https://doi.org/10.1128/mBio.00572-12.

184. S. Zhu, Y.-Z. Cao, C. Jiang, B.-Y. Tan, Z. Wang, S. Feng, L. Zhang, X.-H. Su, B. Brejova, T. Vinar, M. Xu, M.-X. Wang, S.-G. Zhang, M.-R. Huang, R. Wu, Y. Zhou, Sequencing the genome of Marssonina brunnea reveals fungus-poplar co-evolution, BMC Genomics. 13 (2012) 382. https://doi.org/10.1186/1471-2164-13-382.

185. S. Duplessis, C.A. Cuomo, Y.-C. Lin, A. Aerts, E. Tisserant, C. Veneault-Fourrey, D.L. Joly, S. Hacquard, J. Amselem, B.L. Cantarel, R. Chiu, P.M. Coutinho, N. Feau, M. Field, P. Frey, E. Gelhaye, J. Goldberg, M.G. Grabherr, C.D. Kodira, A. Kohler, U. Kües, E.A. Lindquist, S.M. Lucas, R. Mago, E. Mauceli, E. Morin, C. Murat, J.L. Pangilinan, R. Park, M. Pearson, H. Quesneville, N. Rouhier, S. Sakthikumar, A.A. Salamov, J. Schmutz, B. Selles, H. Shapiro, P. Tanguay, G.A. Tuskan, B. Henrissat, Y. Van de Peer, P. Rouzé, J.G. Ellis, P.N. Dodds, J.E. Schein, S. Zhong, R.C. Hamelin, I.V. Grigoriev, L.J. Szabo, F. Martin, Obligate biotrophy features unraveled by the genomic analysis of rust fungi, Proc. Natl. Acad. Sci. U. S. A. 108 (2011) 9166–9171. https://doi.org/10.1073/pnas.1019315108.

186. A. Nemri, D.G.O. Saunders, C. Anderson, N.M. Upadhyaya, J. Win, G.J. Lawrence, D.A. Jones, S. Kamoun, J.G. Ellis, P.N. Dodds, The genome sequence and effector complement of the flax rust pathogen Melampsora lini, Front. Plant Sci. 5 (2014) 98. https://doi.org/10.3389/fpls.2014.00098.

187. Q. Gao, K. Jin, S.-H. Ying, Y. Zhang, G. Xiao, Y. Shang, Z. Duan, X. Hu, X.-Q. Xie, G. Zhou, G. Peng, Z. Luo, W. Huang, B. Wang, W. Fang, S. Wang, Y. Zhong, L.-J. Ma, R.J. St Leger, G.-P. Zhao, Y. Pei, M.-G. Feng, Y. Xia, C. Wang, Genome sequencing and comparative transcriptomics of the model entomopathogenic fungi Metarhizium anisopliae and M. acridum, PLoS Genet. 7 (2011) e1001264. https://doi.org/10.1371/journal.pgen.1001264.

188. X. Hu, G. Xiao, P. Zheng, Y. Shang, Y. Su, X. Zhang, X. Liu, S. Zhan, R.J. St Leger, C. Wang, Trajectory and genomic determinants of fungal-pathogen speciation and host adaptation, Proc. Natl. Acad. Sci. U. S. A. 111 (2014) 16796–16801. https://doi.org/10.1073/pnas.1412662111.

189. S.R. Ahrendt, C.A. Quandt, D. Ciobanu, A. Clum, A. Salamov, B. Andreopoulos, J.-F. Cheng, T. Woyke, A. Pelin, B. Henrissat, N.K. Reynolds, G.L. Benny, M.E. Smith, T.Y. James, I.V. Grigoriev, Leveraging single-cell genomics to expand the fungal tree of life, Nat. Microbiol. 3 (2018) 1417– 1428. https://doi.org/10.1038/s41564-018-0261-0.

190. V. Hershkovitz, N. Sela, L. Taha-Salaime, J. Liu, G. Rafael, C. Kessler, R. Aly, M. Levy, M. Wisniewski, S. Droby, De-novo assembly and characterization of the transcriptome of Metschnikowia fructicola reveals differences in gene expression following interaction with Penicillium digitatum and grapefruit peel, BMC Genomics. 14 (2013) 168. https://doi.org/10.1186/1471-2164-14-168.

191. M.H. Perlin, J. Amselem, E. Fontanillas, S.S. Toh, Z. Chen, J. Goldberg, S. Duplessis, B. Henrissat, S. Young, Q. Zeng, G. Aguileta, E. Petit, H. Badouin, J. Andrews, D. Razeeq, T. Gabaldón, H. Quesneville, T. Giraud, M.E. Hood, D.J. Schultz, C.A. Cuomo, Sex and parasites: genomic and transcriptomic analysis of Microbotryum lychnidis-dioicae, the biotrophic and plant-castrating anther smut fungus, BMC Genomics. 16 (2015) 461. https://doi.org/10.1186/s12864-015-1660-8.

192. A.S. David, S. Haridas, K. LaButti, J. Lim, A. Lipzen, M. Wang, K. Barry, I.V. Grigoriev, J.W. Spatafora, G. May, Draft Genome Sequence of Microdochium bolleyi, a Dark Septate Fungal Endophyte of Beach Grass, Genome Announc. 4 (2016). https://doi.org/10.1128/genomeA.00270-16.

193. A. D.A. Martinez, B.G. Oliver, Y. Gräser, J.M. Goldberg, W. Li, N.M. Martinez-Rossi, M. Monod, E. Shelest, R.C. Barton, E. Birch, A.A. Brakhage, Z. Chen, S.J. Gurr, D. Heiman, J. Heitman, I. Kosti, Rossi, S. Saif, M. Samalova, C.W. Saunders, T. Shea, R.C. Summerbell, J. Xu, S. Young, Q. Zeng, B.W. Birren, C.A. Cuomo, T.C. White, Comparative genome analysis of Trichophyton rubrum and related dermatophytes reveals candidate genes involved in infection, MBio. 3 (2012) e00259–00212. https://doi.org/10.1128/mBio.00259-12.

194. K.L. Haag, T.Y. James, J.-F. Pombert, R. Larsson, T.M.M. Schaer, D. Refardt, D. Ebert, Evolution of a morphological novelty occurred before genome compaction in a lineage of extreme parasites, Proc. Natl. Acad. Sci. U. S. A. 111 (2014) 15480–15485. https://doi.org/10.1073/pnas.1410442111.

195. A. M. Toome, R.A. Ohm, R.W. Riley, T.Y. James, K.L. Lazarus, B. Henrissat, S. Albu, A. Boyd, J. Chow, Clum, G. Heller, A. Lipzen, M. Nolan, L. Sandor, N. Zvenigorodsky, I.V. Grigoriev, J.W. Spatafora, M.C. Aime, Genome sequencing provides insight into the reproductive biology, nutritional mode and ploidy of the fern pathogen Mixia osmundae, New Phytol. 202 (2014) 554–564. https://doi.org/10.1111/nph.12653.

196. S. Lorenz, M. Guenther, C. Grumaz, S. Rupp, S. Zibek, K. Sohn, Genome Sequence of the Basidiomycetous Fungus Pseudozyma aphidis DSM70725, an Efficient Producer of Biosurfactant Mannosylerythritol Lipids, Genome Announc. 2 (2014). https://doi.org/10.1128/genomeA.00053-14.

197. T. Meerupati, K.-M. Andersson, E. Friman, D. Kumar, A. Tunlid, D. Ahrén, Genomic mechanisms accounting for the adaptation to parasitism in nematode-trapping fungi, PLoS Genet. 9 (2013) e1003909. https://doi.org/10.1371/journal.pgen.1003909.

198. J.M.C. Mondego, M.F. Carazzolle, G.G.L. Costa, E.F. Formighieri, L.P. Parizzi, J. Rincones, C. Cotomacci, D.M. Carraro, A.F. Cunha, H. Carrer, R.O. Vidal, R.C. Estrela, O. García, D.P.T. Thomazella, B.V. de Oliveira, A.B. Pires, M.C.S. Rio, M.R.R. Araújo, M.H. de Moraes, L.A.B. Castro, K.P. Gramacho, M.S. Gonçalves, J.P.M. Neto, A.G. Neto, L.V. Barbosa, M.J. Guiltinan, B.A. Bailey, L.W. Meinhardt, J.C. Cascardo, G.A.G. Pereira, A genome survey of Moniliophthora perniciosa gives new insights into Witches’ Broom Disease of cacao, BMC Genomics. 9 (2008) 548. https://doi.org/10.1186/1471-2164-9-548.

199. H. Tan, A. Kohler, R. Miao, T. Liu, Q. Zhang, B. Zhang, L. Jiang, Y. Wang, L. Xie, J. Tang, X. Li, L. Liu, I.V. Grigoriev, C. Daum, K. LaButti, A. Lipzen, A. Kuo, E. Morin, E. Drula, B. Henrissat, B. Wang, Z. Huang, B. Gan, W. Peng, F.M. Martin, Multi-omic analyses of exogenous nutrient bag decomposition by the black morel Morchella importuna reveal sustained carbon acquisition and transferring, Environ. Microbiol. 21 (2019) 3909–3926. https://doi.org/10.1111/1462-2920.14741.

200. J. Uehling, A. Gryganskyi, K. Hameed, T. Tschaplinski, P.K. Misztal, S. Wu, A. Desirò, N. Vande Pol, Z. Du, A. Zienkiewicz, K. Zienkiewicz, E. Morin, E. Tisserant, R. Splivallo, M. Hainaut, B. Henrissat, R. Ohm, A. Kuo, J. Yan, A. Lipzen, M. Nolan, K. LaButti, K. Barry, A.H. Goldstein, J. Labbé, C. Schadt, G. Tuskan, I. Grigoriev, F. Martin, R. Vilgalys, G. Bonito, Comparative genomics of Mortierella elongata and its bacterial endosymbiont Mycoavidus cysteinexigens, Environ. Microbiol. 19 (2017) 2964–2983. https://doi.org/10.1111/1462-2920.13669.

201. A. Lebreton, E. Corre, J.-L. Jany, L. Brillet-Guéguen, C. Pèrez-Arques, V. Garre, M. Monsoor, R. Debuchy, C. Le Meur, E. Coton, G. Barbier, L. Meslet-Cladière, Comparative genomics applied to Mucor species with different lifestyles, BMC Genomics. 21 (2020) 135. https://doi.org/10.1186/s12864-019-6256-2.

202. M.I. Navarro-Mendoza, C. Pérez-Arques, S. Panchal, F.E. Nicolás, S.J. Mondo, P. Ganguly, J. Pangilinan, I.V. Grigoriev, J. Heitman, K. Sanyal, V. Garre, Early Diverging Fungus Mucor circinelloides Lacks Centromeric Histone CENP-A and Displays a Mosaic of Point and Regional Centromeres, Curr. Biol. CB. 29 (2019) 3791–3802.e6. https://doi.org/10.1016/j.cub.2019.09.024.

203. L.M. Corrochano, A. Kuo, M. Marcet-Houben, S. Polaino, A. Salamov, J.M. Villalobos-Escobedo, J. Grimwood, M.I. Álvarez, J. Avalos, D. Bauer, E.P. Benito, I. Benoit, G. Burger, L.P. Camino, D. Cánovas, E. Cerdá-Olmedo, J.-F. Cheng, A. Domínguez, M. Eliáš, A.P. Eslava, F. Glaser, G. Gutiérrez, J. Heitman, B. Henrissat, E.A. Iturriaga, B.F. Lang, J.L. Lavín, S.C. Lee, W. Li, E. Lindquist, S. López-García, E.M. Luque, A.T. Marcos, J. Martin, K. McCluskey, H.R. Medina, A. Miralles-Durán, A. Miyazaki, E. Muñoz-Torres, J.A. Oguiza, R.A. Ohm, M. Olmedo, M. Orejas, L. Ortiz-Castellanos, A.G. Pisabarro, J. Rodríguez-Romero, J. Ruiz-Herrera, R. Ruiz-Vázquez, C. Sanz, W. Schackwitz, M. Shahriari, E. Shelest, F. Silva-Franco, D. Soanes, K. Syed, V.G. Tagua, N.J. Talbot, M.R. Thon, H. Tice, R.P. de Vries, A. Wiebenga, J.S. Yadav, E.L. Braun, S.E. Baker, V. Garre, J. Schmutz, B.A. Horwitz, S. Torres-Martínez, A. Idnurm, A. Herrera-Estrella, T. Gabaldón, I.V. Grigoriev, Expansion of Signal Transduction Pathways in Fungi by Extensive Genome Duplication, Curr. Biol. CB. 26 (2016) 1577–1584. https://doi.org/10.1016/j.cub.2016.04.038.

204. R.M. Berka, I.V. Grigoriev, R. Otillar, A. Salamov, J. Grimwood, I. Reid, N. Ishmael, T. John, C. Darmond, M.-C. Moisan, B. Henrissat, P.M. Coutinho, V. Lombard, D.O. Natvig, E. Lindquist, J. Schmutz, S. Lucas, P. Harris, J. Powlowski, A. Bellemare, D. Taylor, G. Butler, R.P. de Vries, I.E. Allijn, J. van den Brink, S. Ushinsky, R. Storms, A.J. Powell, I.T. Paulsen, L.D.H. Elbourne, S.E. Baker, J. Magnuson, S. Laboissiere, A.J. Clutterbuck, D. Martinez, M. Wogulis, A.L. de Leon, M.W. Rey, A. Tsang, Comparative genomic analysis of the thermophilic biomass-degrading fungi Myceliophthora thermophila and Thielavia terrestris, Nat. Biotechnol. 29 (2011) 922–927. https://doi.org/10.1038/nbt.1976.

205. S.B. Goodwin, S.B. M’barek, B. Dhillon, A.H.J. Wittenberg, C.F. Crane, J.K. Hane, A.J. Foster, T.A.J. Van der Lee, J. Grimwood, A. Aerts, J. Antoniw, A. Bailey, B. Bluhm, J. Bowler, J. Bristow, A. van der Burgt, B. Canto-Canché, A.C.L. Churchill, L. Conde-Ferràez, H.J. Cools, P.M. Coutinho, M. Csukai, P. Dehal, P. De Wit, B. Donzelli, H.C. van de Geest, R.C.H.J. van Ham, K.E. Hammond-Kosack, B. Henrissat, A. Kilian, A.K. Kobayashi, E. Koopmann, Y. Kourmpetis, A. Kuzniar, E. Lindquist, V. Lombard, C. Maliepaard, N. Martins, R. Mehrabi, J.P.H. Nap, A. Ponomarenko, J.J. Rudd, A. Salamov, J. Schmutz, H.J. Schouten, H. Shapiro, I. Stergiopoulos, S.F.F. Torriani, H. Tu, R.P. de Vries, C. Waalwijk, S.B. Ware, A. Wiebenga, L.-H. Zwiers, R.P. Oliver, I.V. Grigoriev, G.H.J. Kema, Finished genome of the fungal wheat pathogen Mycosphaerella graminicola reveals dispensome structure, chromosome plasticity, and stealth pathogenesis, PLoS Genet. 7 (2011) e1002070. https://doi.org/10.1371/journal.pgen.1002070.

206. T. Gabaldón, T. Martin, M. Marcet-Houben, P. Durrens, M. Bolotin-Fukuhara, O. Lespinet, S. Arnaise, S. Boisnard, G. Aguileta, R. Atanasova, C. Bouchier, A. Couloux, S. Creno, J. Almeida Cruz, H. Devillers, A. Enache-Angoulvant, J. Guitard, L. Jaouen, L. Ma, C. Marck, C. Neuvéglise, E. Pelletier, A. Pinard, J. Poulain, J. Recoquillay, E. Westhof, P. Wincker, B. Dujon, C. Hennequin, C. Fairhead, Comparative genomics of emerging pathogens in the Candida glabrata clade, BMC Genomics. 14 (2013) 623. https://doi.org/10.1186/1471-2164-14-623.

207. J.J. Coleman, S.D. Rounsley, M. Rodriguez-Carres, A. Kuo, C.C. Wasmann, J. Grimwood, J. Schmutz, M. Taga, G.J. White, S. Zhou, D.C. Schwartz, M. Freitag, L.-J. Ma, E.G.J. Danchin, B. Henrissat, P.M. Coutinho, D.R. Nelson, D. Straney, C.A. Napoli, B.M. Barker, M. Gribskov, M. Rep, S. Kroken, I. Molnár, C. Rensing, J.C. Kennell, J. Zamora, M.L. Farman, E.U. Selker, A. Salamov, H. Shapiro, J. Pangilinan, E. Lindquist, C. Lamers, I.V. Grigoriev, D.M. Geiser, S.F. Covert, E. Temporini, H.D. Vanetten, The genome of Nectria haematococca: contribution of supernumerary chromosomes to gene expansion, PLoS Genet. 5 (2009) e1000618. https://doi.org/10.1371/journal.pgen.1000618.

208. C.A. Cuomo, C.A. Desjardins, M.A. Bakowski, J. Goldberg, A.T. Ma, J.J. Becnel, E.S. Didier, L. Fan, D.I. Heiman, J.Z. Levin, S. Young, Q. Zeng, E.R. Troemel, Microsporidian genome analysis reveals evolutionary strategies for obligate intracellular growth, Genome Res. 22 (2012) 2478–2488. https://doi.org/10.1101/gr.142802.112.

209. T.A. Nguyen, O.H. Cissé, J. Yun Wong, P. Zheng, D. Hewitt, M. Nowrousian, J.E. Stajich, G. Jedd, Innovation and constraint leading to complex multicellularity in the Ascomycota, Nat. Commun. 8 (2017) 14444. https://doi.org/10.1038/ncomms14444.

210. A. Gómez-Cortecero, R.J. Harrison, A.D. Armitage, Draft Genome Sequence of a European Isolate of the Apple Canker Pathogen Neonectria ditissima, Genome Announc. 3 (2015). https://doi.org/10.1128/genomeA.01243-15.

211. S.E. Baker, W. Schackwitz, A. Lipzen, J. Martin, S. Haridas, K. LaButti, I.V. Grigoriev, B.A. Simmons, K. McCluskey, Draft Genome Sequence of Neurospora crassa Strain FGSC 73, Genome Announc. 3 (2015). https://doi.org/10.1128/genomeA.00074-15.

212. J.E. Galagan, S.E. Calvo, K.A. Borkovich, E.U. Selker, N.D. Read, D. Jaffe, W. FitzHugh, L.-J. Ma, S. Smirnov, S. Purcell, B. Rehman, T. Elkins, R. Engels, S. Wang, C.B. Nielsen, J. Butler, M. Endrizzi, D. Qui, P. Ianakiev, D. Bell-Pedersen, M.A. Nelson, M. Werner-Washburne, C.P. Selitrennikoff, J.A. Kinsey, E.L. Braun, A. Zelter, U. Schulte, G.O. Kothe, G. Jedd, W. Mewes, C. Staben, E. Marcotte, D. Greenberg, A. Roy, K. Foley, J. Naylor, N. Stange-Thomann, R. Barrett, S. Gnerre, M. Kamal, M. Kamvysselis, E. Mauceli, C. Bielke, S. Rudd, D. Frishman, S. Krystofova, C. Rasmussen, R.L. Metzenberg, D.D. Perkins, S. Kroken, C. Cogoni, G. Macino, D. Catcheside, W. Li, R.J. Pratt, S.A. Osmani, C.P.C. DeSouza, L. Glass, M.J. Orbach, J.A. Berglund, R. Voelker, O. Yarden, M. Plamann, S. Seiler, J. Dunlap, A. Radford, R. Aramayo, D.O. Natvig, L.A. Alex, G. Mannhaupt, D.J. Ebbole, M. Freitag, I. Paulsen, M.S. Sachs, E.S. Lander, C. Nusbaum, B. Birren, The genome sequence of the filamentous fungus Neurospora crassa, Nature. 422 (2003) 859– 868. https://doi.org/10.1038/nature01554.

213. C.E. Ellison, J.E. Stajich, D.J. Jacobson, D.O. Natvig, A. Lapidus, B. Foster, A. Aerts, R. Riley, E.A. Lindquist, I.V. Grigoriev, J.W. Taylor, Massive changes in genome architecture accompany the transition to self-fertility in the filamentous fungus Neurospora tetrasperma, Genetics. 189 (2011) 55–69. https://doi.org/10.1534/genetics.111.130690.

214. R.S. Cornman, Y.P. Chen, M.C. Schatz, C. Street, Y. Zhao, B. Desany, M. Egholm, S. Hutchison, J.S. Pettis, W.I. Lipkin, J.D. Evans, Genomic analyses of the microsporidian Nosema ceranae, an emergent pathogen of honey bees, PLoS Pathog. 5 (2009) e1000466. https://doi.org/10.1371/journal.ppat.1000466.

215. O. Miettinen, R. Riley, K. Barry, D. Cullen, R.P. de Vries, M. Hainaut, A. Hatakka, B. Henrissat, K. Hildén, R. Kuo, K. LaButti, A. Lipzen, M.R. Mäkelä, L. Sandor, J.W. Spatafora, I.V. Grigoriev, D.S. Hibbett, Draft Genome Sequence of the White-Rot Fungus Obba rivulosa 3A-2, Genome Announc. 4 (2016). https://doi.org/10.1128/genomeA.00976-16.

216. G.T. Wawrzyn, M.B. Quin, S. Choudhary, F. López-Gallego, C. Schmidt-Dannert, Draft genome of Omphalotus olearius provides a predictive framework for sesquiterpenoid natural product biosynthesis in Basidiomycota, Chem. Biol. 19 (2012) 772–783. https://doi.org/10.1016/j.chembiol.2012.05.012.

217. V. Forgetta, G. Leveque, J. Dias, D. Grove, R. Lyons, S. Genik, C. Wright, S. Singh, N. Peterson, M. Zianni, J. Kieleczawa, R. Steen, A. Perera, D. Bintzler, S. Adams, W. Hintz, V. Jacobi, L. Bernier, R. Levesque, K. Dewar, Sequencing of the Dutch elm disease fungus genome using the Roche/454 GS-FLX Titanium System in a comparison of multiple genomics core facilities, J. Biomol. Tech. JBT. 24 (2013) 39–49. https://doi.org/10.7171/jbt.12-2401-005.

218. S. Haridas, Y. Wang, L. Lim, S. Massoumi Alamouti, S. Jackman, R. Docking, G. Robertson, I. Birol, J. Bohlmann, C. Breuil, The genome and transcriptome of the pine saprophyte Ophiostoma piceae, and a comparison with the bark beetle-associated pine pathogen Grosmannia clavigera, BMC Genomics. 14 (2013) 373. https://doi.org/10.1186/1471-2164-14-373.

219. N.H. Youssef, M.B. Couger, C.G. Struchtemeyer, A.S. Liggenstoffer, R.A. Prade, F.Z. Najar, H.K. Atiyeh, M.R. Wilkins, M.S. Elshahed, The genome of the anaerobic fungus Orpinomyces sp. strain C1A reveals the unique evolutionary history of a remarkable plant biomass degrader, Appl. Environ. Microbiol. 79 (2013) 4620–4634. https://doi.org/10.1128/AEM.00821-13.

220. M.N. Biango-Daniels, T.W. Wang, K.T. Hodge, Draft Genome Sequence of the Patulin-Producing Fungus Paecilomyces niveus Strain CO7, Genome Announc. 6 (2018). https://doi.org/10.1128/genomeA.00556-18.

221. A.S. Urquhart, S.J. Mondo, M.R. Mäkelä, J.K. Hane, A. Wiebenga, G. He, S. Mihaltcheva, J. Pangilinan, A. Lipzen, K. Barry, R.P. de Vries, I.V. Grigoriev, A. Idnurm, Genomic and Genetic Insights Into a Cosmopolitan Fungus, Paecilomyces variotii (Eurotiales), Front. Microbiol. 9 (2018) 3058. https://doi.org/10.3389/fmicb.2018.03058.

222. C.A. Desjardins, M.D. Champion, J.W. Holder, A. Muszewska, J. Goldberg, A.M. Bailão, M.M. Brigido, M.E. da S. Ferreira, A.M. Garcia, M. Grynberg, S. Gujja, D.I. Heiman, M.R. Henn, C.D. Kodira, H. León-Narváez, L.V.G. Longo, L.-J. Ma, I. Malavazi, A.L. Matsuo, F.V. Morais, M. Pereira, S. Rodríguez-Brito, S. Sakthikumar, S.M. Salem-Izacc, S.M. Sykes, M.M. Teixeira, M.C. Vallejo, M.E.M.T. Walter, C. Yandava, S. Young, Q. Zeng, J. Zucker, M.S. Felipe, G.H. Goldman, B.J. Haas, J.G. McEwen, G. Nino-Vega, R. Puccia, G. San-Blas, C.M. de A. Soares, B.W. Birren, C.A. Cuomo, Comparative genomic analysis of human fungal pathogens causing paracoccidioidomycosis, PLoS Genet. 7 (2011) e1002345. https://doi.org/10.1371/journal.pgen.1002345.

223. C.A. Zeiner, S.O. Purvine, E.M. Zink, L. Paša-Tolić, D.L. Chaput, S. Haridas, S. Wu, K. LaButti, I.V. Grigoriev, B. Henrissat, C.M. Santelli, C.M. Hansel, Comparative Analysis of Secretome Profiles of Manganese(II)-Oxidizing Ascomycete Fungi, PloS One. 11 (2016) e0157844. https://doi.org/10.1371/journal.pone.0157844.

224. J.C. Nielsen, S. Grijseels, S. Prigent, B. Ji, J. Dainat, K.F. Nielsen, J.C. Frisvad, M. Workman, J. Nielsen, Global analysis of biosynthetic gene clusters reveals vast potential of secondary metabolite production in Penicillium species, Nat. Microbiol. 2 (2017) 17044. https://doi.org/10.1038/nmicrobiol.2017.44.

225. M.A. van den Berg, R. Albang, K. Albermann, J.H. Badger, J.-M. Daran, A.J.M. Driessen, C. Garcia-Estrada, N.D. Fedorova, D.M. Harris, W.H.M. Heijne, V. Joardar, J.A.K.W. Kiel, A. Kovalchuk, J.F. Martín, W.C. Nierman, J.G. Nijland, J.T. Pronk, J.A. Roubos, I.J. van der Klei, N.N.M.E. van Peij, M. Veenhuis, H. von Döhren, C. Wagner, J. Wortman, R.A.L. Bovenberg, Genome sequencing and analysis of the filamentous fungus Penicillium chrysogenum, Nat. Biotechnol. 26 (2008) 1161–1168. https://doi.org/10.1038/nbt.1498.

226. M. Marcet-Houben, A.-R. Ballester, B. de la Fuente, E. Harries, J.F. Marcos, L. González-Candelas, T. Gabaldón, Genome sequence of the necrotrophic fungus Penicillium digitatum, the main postharvest pathogen of citrus, BMC Genomics. 13 (2012) 646. https://doi.org/10.1186/1471-2164-13-646.

227. A.-R. Ballester, M. Marcet-Houben, E. Levin, N. Sela, C. Selma-Lázaro, L. Carmona, M. Wisniewski, S. Droby, L. González-Candelas, T. Gabaldón, Genome, Transcriptome, and Functional Analyses of Penicillium expansum Provide New Insights Into Secondary Metabolism and Pathogenicity, Mol. Plant-Microbe Interact. MPMI. 28 (2015) 232–248. https://doi.org/10.1094/MPMI-09-14-0261-FI.

228. H. Banani, M. Marcet-Houben, A.-R. Ballester, P. Abbruscato, L. González-Candelas, T. Gabaldón, D. Spadaro, Genome sequencing and secondary metabolism of the postharvest pathogen Penicillium griseofulvum, BMC Genomics. 17 (2016) 19. https://doi.org/10.1186/s12864-015-2347-x.

229. B.D. Wingfield, I. Barnes, Z. Wilhelm de Beer, L. De Vos, T.A. Duong, A.M. Kanzi, K. Naidoo, H.D.T. Nguyen, Q.C. Santana, M. Sayari, K.A. Seifert, E.T. Steenkamp, C. Trollip, N.A. van der Merwe, M.A. van der Nest, P. Markus Wilken, M.J. Wingfield, IMA Genome-F 5: Draft genome sequences of Ceratocystis eucalypticola, Chrysoporthe cubensis, C. deuterocubensis, Davidsoniella virescens, Fusarium temperatum,Graphilbum fragrans, Penicillium nordicum, and Thielaviopsis musarum, IMA Fungus. 6 (2015) 493–506. https://doi.org/10.5598/imafungus.2015.06.02.13.

230. G. Liu, L. Zhang, X. Wei, G. Zou, Y. Qin, L. Ma, J. Li, H. Zheng, S. Wang, C. Wang, L. Xun, G.-P. Zhao, Z. Zhou, Y. Qu, Genomic and secretomic analyses reveal unique features of the lignocellulolytic enzyme system of Penicillium decumbens, PloS One. 8 (2013) e55185. https://doi.org/10.1371/journal.pone.0055185.

231. M. Peng, A. Dilokpimol, M.R. Mäkelä, K. Hildén, S. Bervoets, R. Riley, I.V. Grigoriev, M. Hainaut, B. Henrissat, R.P. de Vries, Z. Granchi, The draft genome sequence of the ascomycete fungus Penicillium subrubescens reveals a highly enriched content of plant biomass related CAZymes compared to related fungi, J. Biotechnol. 246 (2017) 1–3. https://doi.org/10.1016/j.jbiotec.2017.02.012.

232. H.D.T. Nguyen, D.R. McMullin, E. Ponomareva, R. Riley, K.R. Pomraning, S.E. Baker, K.A. Seifert, Ochratoxin A production by Penicillium thymicola, Fungal Biol. 120 (2016) 1041–1049. https://doi.org/10.1016/j.funbio.2016.04.002.

233. D.G. Knapp, J.B. Németh, K. Barry, M. Hainaut, B. Henrissat, J. Johnson, A. Kuo, J.H.P. Lim, A. Lipzen, M. Nolan, R.A. Ohm, L. Tamás, I.V. Grigoriev, J.W. Spatafora, L.G. Nagy, G.M. Kovács, Comparative genomics provides insights into the lifestyle and reveals functional heterogeneity of dark septate endophytic fungi, Sci. Rep. 8 (2018) 6321. https://doi.org/10.1038/s41598-018-24686-4.

234. A. Morales-Cruz, K.C.H. Amrine, B. Blanco-Ulate, D.P. Lawrence, R. Travadon, P.E. Rolshausen, K. Baumgartner, D. Cantu, Distinctive expansion of gene families associated with plant cell wall degradation, secondary metabolism, and nutrient uptake in the genomes of grapevine trunk pathogens, BMC Genomics. 16 (2015) 469. https://doi.org/10.1186/s12864-015-1624-z.

235. H. Suzuki, J. MacDonald, K. Syed, A. Salamov, C. Hori, A. Aerts, B. Henrissat, A. Wiebenga, P.A. VanKuyk, K. Barry, E. Lindquist, K. LaButti, A. Lapidus, S. Lucas, P. Coutinho, Y. Gong, M. Samejima, R. Mahadevan, M. Abou-Zaid, R.P. de Vries, K. Igarashi, J.S. Yadav, I.V. Grigoriev, E.R. Master, Comparative genomics of the white-rot fungi, Phanerochaete carnosa and P. chrysosporium, to elucidate the genetic basis of the distinct wood types they colonize, BMC Genomics. 13 (2012) 444. https://doi.org/10.1186/1471-2164-13-444.

236. R.A. Ohm, R. Riley, A. Salamov, B. Min, I.-G. Choi, I.V. Grigoriev, Genomics of wood-degrading fungi, Fungal Genet. Biol. FG B. 72 (2014) 82–90. https://doi.org/10.1016/j.fgb.2014.05.001.

237. A.K. Walker, S.L. Frasz, K.A. Seifert, J.D. Miller, S.J. Mondo, K. LaButti, A. Lipzen, R.B. Dockter, M.C. Kennedy, I.V. Grigoriev, J.W. Spatafora, Full Genome of Phialocephala scopiformis DAOMC 229536, a Fungal Endophyte of Spruce Producing the Potent Anti-Insectan Compound Rugulosin, Genome Announc. 4 (2016). https://doi.org/10.1128/genomeA.01768-15.

238. L.F. Moreno, J.B. Stielow, M. de Vries, V.A. Weiss, V.A. Vicente, S. de Hoog, Draft Genome Sequence of the Ant-Associated Fungus Phialophora attae (CBS 131958), Genome Announc. 3 (2015). https://doi.org/10.1128/genomeA.01099-15.

239. M.R. Mäkelä, M. Peng, Z. Granchi, T. Chin-A-Woeng, R. Hegi, S.I. van Pelt, S. Ahrendt, R. Riley, M. Hainaut, B. Henrissat, I.V. Grigoriev, R.P. de Vries, K.S. Hildén, Draft Genome Sequence of the Basidiomycete White-Rot Fungus Phlebia centrifuga, Genome Announc. 6 (2018). https://doi.org/10.1128/genomeA.01414-17.

240. J. Kuuskeri, M. Häkkinen, P. Laine, O.-P. Smolander, F. Tamene, S. Miettinen, P. Nousiainen, M. Kemell, P. Auvinen, T. Lundell, Time-scale dynamics of proteome and transcriptome of the white-rot fungus Phlebia radiata: growth on spruce wood and decay effect on lignocellulose, Biotechnol. Biofuels. 9 (2016) 192. https://doi.org/10.1186/s13068-016-0608-9.

241. C. Hori, T. Ishida, K. Igarashi, M. Samejima, H. Suzuki, E. Master, P. Ferreira, F.J. Ruiz-Dueñas, B. Held, P. Canessa, L.F. Larrondo, M. Schmoll, I.S. Druzhinina, C.P. Kubicek, J.A. Gaskell, P. Kersten, F. St John, J. Glasner, G. Sabat, S. Splinter BonDurant, K. Syed, J. Yadav, A.C. Mgbeahuruike, A. Kovalchuk, F.O. Asiegbu, G. Lackner, D. Hoffmeister, J. Rencoret, A. Gutiérrez, H. Sun, E. Lindquist, K. Barry, R. Riley, I.V. Grigoriev, B. Henrissat, U. Kües, R.M. Berka, A.T. Martínez, S.F. Covert, R.A. Blanchette, D. Cullen, Analysis of the Phlebiopsis gigantea genome, transcriptome and secretome provides insight into its pioneer colonization strategies of wood, PLoS Genet. 10 (2014) e1004759. https://doi.org/10.1371/journal.pgen.1004759.

242. V. Guarnaccia, T. Gehrmann, G.J. Silva-Junior, P.H. Fourie, S. Haridas, D. Vu, J. Spatafora, F.M. Martin, V. Robert, I.V. Grigoriev, J.Z. Groenewald, P.W. Crous, Phyllosticta citricarpa and sister species of global importance to Citrus, Mol. Plant Pathol. 20 (2019) 1619–1635. https://doi.org/10.1111/mpp.12861.

243. A.P. Douglass, B. Offei, S. Braun-Galleani, A.Y. Coughlan, A.A.R. Martos, R.A. Ortiz-Merino, K.P. Byrne, K.H. Wolfe, Population genomics shows no distinction between pathogenic Candida krusei and environmental Pichia kudriavzevii: One species, four names, PLoS Pathog. 14 (2018) e1007138. https://doi.org/10.1371/journal.ppat.1007138.

244. K. De Schutter, Y.-C. Lin, P. Tiels, A. Van Hecke, S. Glinka, J. Weber-Lehmann, P. Rouzé, Y. Van de Peer, N. Callewaert, Genome sequence of the recombinant protein production host Pichia pastoris, Nat. Biotechnol. 27 (2009) 561–566. https://doi.org/10.1038/nbt.1544.

245. A. Zuccaro, U. Lahrmann, U. Güldener, G. Langen, S. Pfiffi, D. Biedenkopf, P. Wong, B. Samans, C. Grimm, M. Basiewicz, C. Murat, F. Martin, K.-H. Kogel, Endophytic life strategies decoded by genome and transcriptome analyses of the mutualistic root symbiont Piriformospora indica, PLoS Pathog. 7 (2011) e1002290. https://doi.org/10.1371/journal.ppat.1002290.

246. M. Alfaro, R. Castanera, J.L. Lavín, I.V. Grigoriev, J.A. Oguiza, L. Ramírez, A.G. Pisabarro, Comparative and transcriptional analysis of the predicted secretome in the lignocellulose-degrading basidiomycete fungus Pleurotus ostreatus, Environ. Microbiol. 18 (2016) 4710–4726. https://doi.org/10.1111/1462-2920.13360.

247. O.H. Cissé, M. Pagni, P.M. Hauser, De novo assembly of the Pneumocystis jirovecii genome from a single bronchoalveolar lavage fluid specimen from a patient, MBio. 4 (2012) e00428–00412. https://doi.org/10.1128/mBio.00428-12.

248. G. Wang, Z. Liu, R. Lin, E. Li, Z. Mao, J. Ling, Y. Yang, W.-B. Yin, B. Xie, Biosynthesis of Antibiotic Leucinostatins in Bio-control Fungus Purpureocillium lilacinum and Their Inhibition on Phytophthora Revealed by Genome Mining, PLoS Pathog. 12 (2016) e1005685. https://doi.org/10.1371/journal.ppat.1005685.

249. E. Espagne, O. Lespinet, F. Malagnac, C. Da Silva, O. Jaillon, B.M. Porcel, A. Couloux, J.-M. Aury, B. Ségurens, J. Poulain, V. Anthouard, S. Grossetete, H. Khalili, E. Coppin, M. Déquard-Chablat, M. Picard, V. Contamine, S. Arnaise, A. Bourdais, V. Berteaux-Lecellier, D. Gautheret, R.P. de Vries, E. Battaglia, P.M. Coutinho, E.G. Danchin, B. Henrissat, R.E. Khoury, A. Sainsard-Chanet, A. Boivin, B. Pinan-Lucarré, C.H. Sellem, R. Debuchy, P. Wincker, J. Weissenbach, P. Silar, The genome sequence of the model ascomycete fungus Podospora anserina, Genome Biol. 9 (2008) R77. https://doi.org/10.1186/gb-2008-9-5-r77.

250. S. Miyauchi, A. Rancon, E. Drula, H. Hage, D. Chaduli, A. Favel, S. Grisel, B. Henrissat, I. Herpoël-Gimbert, F.J. Ruiz-Dueñas, D. Chevret, M. Hainaut, J. Lin, M. Wang, J. Pangilinan, A. Lipzen, L. Lesage-Meessen, D. Navarro, R. Riley, I.V. Grigoriev, S. Zhou, S. Raouche, M.-N. Rosso, Integrative visual omics of the white-rot fungus Polyporus brumalis exposes the biotechnological potential of its oxidative enzymes for delignifying raw plant biomass, Biotechnol. Biofuels. 11 (2018) 201. https://doi.org/10.1186/s13068-018-1198-5.

251. D. Martinez, J. Challacombe, I. Morgenstern, D. Hibbett, M. Schmoll, C.P. Kubicek, P. Ferreira, F.J. Ruiz-Duenas, A.T. Martinez, P. Kersten, K.E. Hammel, A. Vanden Wymelenberg, J. Gaskell, E. Lindquist, G. Sabat, S.S. Bondurant, L.F. Larrondo, P. Canessa, R. Vicuna, J. Yadav, H. Doddapaneni, V. Subramanian, A.G. Pisabarro, J.L. Lavín, J.A. Oguiza, E. Master, B. Henrissat, P.M. Coutinho, P. Harris, J.K. Magnuson, S.E. Baker, K. Bruno, W. Kenealy, P.J. Hoegger, U. Kües, P. Ramaiya, S. Lucas, A. Salamov, H. Shapiro, H. Tu, C.L. Chee, M. Misra, G. Xie, S. Teter, D. Yaver, T. James, M. Mokrejs, M. Pospisek, I.V. Grigoriev, T. Brettin, D. Rokhsar, R. Berka, D. Cullen, Genome, transcriptome, and secretome analysis of wood decay fungus Postia placenta supports unique mechanisms of lignocellulose conversion, Proc. Natl. Acad. Sci. U. S. A. 106 (2009) 1954–1959. https://doi.org/10.1073/pnas.0809575106.

252. J. Gaskell, P. Kersten, L.F. Larrondo, P. Canessa, D. Martinez, D. Hibbett, M. Schmoll, C.P. Kubicek, A.T. Martinez, J. Yadav, E. Master, J.K. Magnuson, D. Yaver, R. Berka, K. Lail, C. Chen, K. LaButti, M. Nolan, A. Lipzen, A. Aerts, R. Riley, K. Barry, B. Henrissat, R. Blanchette, I.V. Grigoriev, D. Cullen, Draft genome sequence of a monokaryotic model brown-rot fungus Postia (Rhodonia) placenta SB12, Genomics Data. 14 (2017) 21–23. https://doi.org/10.1016/j.gdata.2017.08.003.

253. R.E. Arango Isaza, C. Diaz-Trujillo, B. Dhillon, A. Aerts, J. Carlier, C.F. Crane, T. V de Jong, I. de Vries, R. Dietrich, A.D. Farmer, C. Fortes Fereira, S. Garcia, M. Guzman, R.C. Hamelin, E.A. Lindquist, R. Mehrabi, O. Quiros, J. Schmutz, H. Shapiro, E. Reynolds, G. Scalliet, M. Souza, I. Stergiopoulos, T.A.J. Van der Lee, P.J.G.M. De Wit, M.-F. Zapater, L.-H. Zwiers, I.V. Grigoriev, S.B. Goodwin, G.H.J. Kema, Combating a Global Threat to a Clonal Crop: Banana Black Sigatoka Pathogen Pseudocercospora fijiensis (Synonym Mycosphaerella fijiensis) Genomes Reveal Clues for Disease Control, PLoS Genet. 12 (2016) e1005876. https://doi.org/10.1371/journal.pgen.1005876.

254. K.P. Drees, J.M. Palmer, R. Sebra, J.M. Lorch, C. Chen, C.-C. Wu, J.W. Bok, N.P. Keller, D.S. Blehert, C.A. Cuomo, D.L. Lindner, J.T. Foster, Use of Multiple Sequencing Technologies To Produce a High-Quality Genome of the Fungus Pseudogymnoascus destructans, the Causative Agent of Bat White-Nose Syndrome, Genome Announc. 4 (2016). https://doi.org/10.1128/genomeA.00445-16.

255. T. Morita, H. Koike, Y. Koyama, H. Hagiwara, E. Ito, T. Fukuoka, T. Imura, M. Machida, D. Kitamoto, Genome Sequence of the Basidiomycetous Yeast Pseudozyma antarctica T-34, a Producer of the Glycolipid Biosurfactants Mannosylerythritol Lipids, Genome Announc. 1 (2013) e0006413. https://doi.org/10.1128/genomeA.00064-13.

256. M. Konishi, Y. Hatada, J.-I. Horiuchi, Draft Genome Sequence of the Basidiomycetous Yeast-Like Fungus Pseudozyma hubeiensis SY62, Which Produces an Abundant Amount of the Biosurfactant Mannosylerythritol Lipids, Genome Announc. 1 (2013). https://doi.org/10.1128/genomeA.00409-13.

257. J. Fricke, F. Blei, D. Hoffmeister, Enzymatic Synthesis of Psilocybin, Angew. Chem. Int. Ed Engl. 56 (2017) 12352–12355. https://doi.org/10.1002/anie.201705489.

258. E.S. Nazareno, F. Li, M. Smith, R.F. Park, S.F. Kianian, M. Figueroa, Puccinia coronata f. sp. avenae: a threat to global oat production, Mol. Plant Pathol. 19 (2018) 1047–1060. https://doi.org/10.1111/mpp.12608.

259. F. Li, N.M. Upadhyaya, J. Sperschneider, O. Matny, H. Nguyen-Phuc, R. Mago, C. Raley, M.E. Miller, K.A.T. Silverstein, E. Henningsen, C.D. Hirsch, B. Visser, Z.A. Pretorius, B.J. Steffenson, B. Schwessinger, P.N. Dodds, M. Figueroa, Emergence of the Ug99 lineage of the wheat stem rust pathogen through somatic hybridisation, Nat. Commun. 10 (2019) 5068. https://doi.org/10.1038/s41467-019-12927-7.

260. B. Schwessinger, J. Sperschneider, W.S. Cuddy, D.P. Garnica, M.E. Miller, J.M. Taylor, P.N. Dodds, M. Figueroa, R.F. Park, J.P. Rathjen, A Near-Complete Haplotype-Phased Genome of the Dikaryotic Wheat Stripe Rust Fungus Puccinia striiformis f. sp. tritici Reveals High Interhaplotype Diversity, MBio. 9 (2018). https://doi.org/10.1128/mBio.02275-17.

261. D. Cantu, M. Govindarajulu, A. Kozik, M. Wang, X. Chen, K.K. Kojima, J. Jurka, R.W. Michelmore, J. Dubcovsky, Next generation sequencing provides rapid access to the genome of Puccinia striiformis f. sp. tritici, the causal agent of wheat stripe rust, PloS One. 6 (2011) e24230. https://doi.org/10.1371/journal.pone.0024230.

262. C.A. Cuomo, G. Bakkeren, H.B. Khalil, V. Panwar, D. Joly, R. Linning, S. Sakthikumar, X. Song, X. Adiconis, L. Fan, J.M. Goldberg, J.Z. Levin, S. Young, Q. Zeng, Y. Anikster, M. Bruce, M. Wang, C. Yin, B. McCallum, L.J. Szabo, S. Hulbert, X. Chen, J.P. Fellers, Comparative Analysis Highlights Variable Genome Content of Wheat Rusts and Divergence of the Mating Loci, G3 Bethesda Md. 7 (2017) 361–376. https://doi.org/10.1534/g3.116.032797.

263. A. Levasseur, A. Lomascolo, O. Chabrol, F.J. Ruiz-Dueñas, E. Boukhris-Uzan, F. Piumi, U. Kües, A.F.J. Ram, C. Murat, M. Haon, I. Benoit, Y. Arfi, D. Chevret, E. Drula, M.J. Kwon, P. Gouret, L. Lesage-Meessen, V. Lombard, J. Mariette, C. Noirot, J. Park, A. Patyshakuliyeva, J.C. Sigoillot, A. Wiebenga, H.A.B. Wösten, F. Martin, P.M. Coutinho, R.P. de Vries, A.T. Martínez, C. Klopp, P. Pontarotti, B. Henrissat, E. Record, The genome of the white-rot fungus Pycnoporus cinnabarinus: a basidiomycete model with a versatile arsenal for lignocellulosic biomass breakdown, BMC Genomics. 15 (2014) 486. https://doi.org/10.1186/1471-2164-15-486.

264. S. Miyauchi, H. Hage, E. Drula, L. Lesage-Meessen, J.-G. Berrin, D. Navarro, A. Favel, D. Chaduli, S. Grisel, M. Haon, F. Piumi, A. Levasseur, A. Lomascolo, S. Ahrendt, K. Barry, K.M. LaButti, D. Chevret, C. Daum, J. Mariette, C. Klopp, D. Cullen, R.P. de Vries, A.C. Gathman, M. Hainaut, B. Henrissat, K.S. Hildén, U. Kües, W. Lilly, A. Lipzen, M.R. Mäkelä, A.T. Martinez, M. Morel-Rouhier, E. Morin, J. Pangilinan, A.F.J. Ram, H.A.B. Wösten, F.J. Ruiz-Dueñas, R. Riley, E. Record, I.V. Grigoriev, M.-N. Rosso, Conserved white-rot enzymatic mechanism for wood decay in the Basidiomycota genus Pycnoporus, DNA Res. Int. J. Rapid Publ. Rep. Genes Genomes. 27 (2020). https://doi.org/10.1093/dnares/dsaa011.

265. S.R. Ellwood, Z. Liu, R.A. Syme, Z. Lai, J.K. Hane, F. Keiper, C.S. Moffat, R.P. Oliver, T.L. Friesen, A first genome assembly of the barley fungal pathogen Pyrenophora teres f. teres, Genome Biol. 11 (2010) R109. https://doi.org/10.1186/gb-2010-11-11-r109.

266. V.A. Manning, I. Pandelova, B. Dhillon, L.J. Wilhelm, S.B. Goodwin, A.M. Berlin, M. Figueroa, M. Freitag, J.K. Hane, B. Henrissat, W.H. Holman, C.D. Kodira, J. Martin, R.P. Oliver, B. Robbertse, W. Schackwitz, D.C. Schwartz, J.W. Spatafora, B.G. Turgeon, C. Yandava, S. Young, S. Zhou, Q. Zeng, I.V. Grigoriev, L.-J. Ma, L.M. Ciuffetti, Comparative genomics of a plant-pathogenic fungus, Pyrenophora tritici-repentis, reveals transduplication and the impact of repeat elements on pathogenicity and population divergence, G3 Bethesda Md. 3 (2013) 41–63. https://doi.org/10.1534/g3.112.004044.

267. S. Traeger, F. Altegoer, M. Freitag, T. Gabaldon, F. Kempken, A. Kumar, M. Marcet-Houben, S. Pöggeler, J.E. Stajich, M. Nowrousian, The genome and development-dependent transcriptomes of Pyronema confluens: a window into fungal evolution, PLoS Genet. 9 (2013) e1003820. https://doi.org/10.1371/journal.pgen.1003820.

268. D. Wibberg, L. Jelonek, O. Rupp, M. Hennig, F. Eikmeyer, A. Goesmann, A. Hartmann, R. Borriss, R. Grosch, A. Pühler, A. Schlüter, Establishment and interpretation of the genome sequence of the phytopathogenic fungus Rhizoctonia solani AG1-IB isolate 7/3/14, J. Biotechnol. 167 (2013) 142–155. https://doi.org/10.1016/j.jbiotec.2012.12.010.

269. E.C.H. Chen, E. Morin, D. Beaudet, J. Noel, G. Yildirir, S. Ndikumana, P. Charron, C. St-Onge, J. Giorgi, M. Krüger, T. Marton, J. Ropars, I.V. Grigoriev, M. Hainaut, B. Henrissat, C. Roux, F. Martin, N. Corradi, High intraspecific genome diversity in the model arbuscular mycorrhizal symbiont Rhizophagus irregularis, New Phytol. 220 (2018) 1161–1171. https://doi.org/10.1111/nph.14989.

270. E. Tisserant, M. Malbreil, A. Kuo, A. Kohler, A. Symeonidi, R. Balestrini, P. Charron, N. Duensing, N. Frei dit Frey, V. Gianinazzi-Pearson, L.B. Gilbert, Y. Handa, J.R. Herr, M. Hijri, R. Koul, M. Kawaguchi, F. Krajinski, P.J. Lammers, F.G. Masclaux, C. Murat, E. Morin, S. Ndikumana, M. Pagni, D. Petitpierre, N. Requena, P. Rosikiewicz, R. Riley, K. Saito, H. San Clemente, H. Shapiro, D. van Tuinen, G. Bécard, P. Bonfante, U. Paszkowski, Y.Y. Shachar-Hill, G.A. Tuskan, J.P.W. Young, P.W. Young, I.R. Sanders, B. Henrissat, S.A. Rensing, I.V. Grigoriev, N. Corradi, C. Roux, F. Martin, Genome of an arbuscular mycorrhizal fungus provides insight into the oldest plant symbiosis, Proc. Natl. Acad. Sci. U. S. A. 110 (2013) 20117–20122. https://doi.org/10.1073/pnas.1313452110.

271. A.B. Mujic, A. Kuo, A. Tritt, A. Lipzen, C. Chen, J. Johnson, A. Sharma, K. Barry, I.V. Grigoriev, J.W. Spatafora, Comparative Genomics of the Ectomycorrhizal Sister Species Rhizopogon vinicolor and Rhizopogon vesiculosus (Basidiomycota: Boletales) Reveals a Divergence of the Mating Type B Locus, G3 Bethesda Md. 7 (2017) 1775–1789. https://doi.org/10.1534/g3.117.039396.

272. L.-J. Ma, A.S. Ibrahim, C. Skory, M.G. Grabherr, G. Burger, M. Butler, M. Elias, A. Idnurm, B.F. Lang, T. Sone, A. Abe, S.E. Calvo, L.M. Corrochano, R. Engels, J. Fu, W. Hansberg, J.-M. Kim, C.D. Kodira, M.J. Koehrsen, B. Liu, D. Miranda-Saavedra, S. O’Leary, L. Ortiz-Castellanos, R. Poulter, J. Rodriguez-Romero, J. Ruiz-Herrera, Y.-Q. Shen, Q. Zeng, J. Galagan, B.W. Birren, C.A. Cuomo, B.L. Wickes, Genomic analysis of the basal lineage fungus Rhizopus oryzae reveals a whole-genome duplication, PLoS Genet. 5 (2009) e1000549. https://doi.org/10.1371/journal.pgen.1000549.

273. O.A. Lastovetsky, M.L. Gaspar, S.J. Mondo, K.M. LaButti, L. Sandor, I.V. Grigoriev, S.A. Henry, T.E. Pawlowska, Lipid metabolic changes in an early divergent fungus govern the establishment of a mutualistic symbiosis with endobacteria, Proc. Natl. Acad. Sci. U. S. A. 113 (2016) 15102– 15107. https://doi.org/10.1073/pnas.1615148113.

274. D. Wang, R. Wu, Y. Xu, M. Li, Draft Genome Sequence of Rhizopus chinensis CCTCCM201021, Used for Brewing Traditional Chinese Alcoholic Beverages, Genome Announc. 1 (2013) e0019512. https://doi.org/10.1128/genomeA.00195-12.

275. S. Zhang, J.M. Skerker, C.D. Rutter, M.J. Maurer, A.P. Arkin, C.V. Rao, Engineering Rhodosporidium toruloides for increased lipid production, Biotechnol. Bioeng. 113 (2016) 1056–1066. https://doi.org/10.1002/bit.25864.

276. S.T. Coradetti, D. Pinel, G.M. Geiselman, M. Ito, S.J. Mondo, M.C. Reilly, Y.-F. Cheng, S. Bauer, I.V. Grigoriev, J.M. Gladden, B.A. Simmons, R.B. Brem, A.P. Arkin, J.M. Skerker, Functional genomics of lipid metabolism in the oleaginous yeast Rhodosporidium toruloides, ELife. 7 (2018). https://doi.org/10.7554/eLife.32110.

277. Z. Zhu, S. Zhang, H. Liu, H. Shen, X. Lin, F. Yang, Y.J. Zhou, G. Jin, M. Ye, H. Zou, H. Zou, Z.K. Zhao, A multi-omic map of the lipid-producing yeast Rhodosporidium toruloides, Nat. Commun. 3 (2012) 1112. https://doi.org/10.1038/ncomms2112.

278. A. Firrincieli, R. Otillar, A. Salamov, J. Schmutz, Z. Khan, R.S. Redman, N.D. Fleck, E. Lindquist, I.V. Grigoriev, S.L. Doty, Genome sequence of the plant growth promoting endophytic yeast Rhodotorula graminis WP1, Front. Microbiol. 6 (2015) 978. https://doi.org/10.3389/fmicb.2015.00978.

279. J. Goordial, I. Raymond-Bouchard, R. Riley, J. Ronholm, N. Shapiro, T. Woyke, K.M. LaButti, H. Tice, M. Amirebrahimi, I.V. Grigoriev, C. Greer, C. Bakermans, L. Whyte, Improved High-Quality Draft Genome Sequence of the Eurypsychrophile Rhodotorula sp. JG1b, Isolated from Permafrost in the Hyperarid Upper-Elevation McMurdo Dry Valleys, Antarctica, Genome Announc. 4 (2016). https://doi.org/10.1128/genomeA.00069-16.

280. H.B. Korotkin, R.A. Swenie, O. Miettinen, J.M. Budke, K.-H. Chen, F. Lutzoni, M.E. Smith, P.B. Matheny, Stable isotope analyses reveal previously unknown trophic mode diversity in the Hymenochaetales, Am. J. Bot. 105 (2018) 1869–1887. https://doi.org/10.1002/ajb2.1183.

281. K. Krizsán, É. Almási, Z. Merényi, N. Sahu, M. Virágh, T. Kószó, S. Mondo, B. Kiss, B. Bálint, U. Kües, K. Barry, J. Cseklye, B. Hegedüs, B. Henrissat, J. Johnson, A. Lipzen, R.A. Ohm, I. Nagy, J. Pangilinan, J. Yan, Y. Xiong, I.V. Grigoriev, D.S. Hibbett, L.G. Nagy, Transcriptomic atlas of mushroom development reveals conserved genes behind complex multicellularity in fungi, Proc. Natl. Acad. Sci. U. S. A. 116 (2019) 7409–7418. https://doi.org/10.1073/pnas.1817822116.

282. A. A.O. Oghenekaro, A. Kovalchuk, T. Raffaello, S. Camarero, M. Gressler, B. Henrissat, J. Lee, M. Liu, A.T. Martínez, O. Miettinen, S. Mihaltcheva, J. Pangilinan, F. Ren, R. Riley, F.J. Ruiz-Dueñas, Serrano, M.R. Thon, Z. Wen, Z. Zeng, K. Barry, I.V. Grigoriev, F. Martin, F.O. Asiegbu, Genome sequencing of Rigidoporus microporus provides insights on genes important for wood decay, latex tolerance and interspecific fungal interactions, Sci. Rep. 10 (2020) 5250. https://doi.org/10.1038/s41598-020-62150-4.

283. T.Y. James, A. Pelin, L. Bonen, S. Ahrendt, D. Sain, N. Corradi, J.E. Stajich, Shared signatures of parasitism and phylogenomics unite Cryptomycota and microsporidia, Curr. Biol. CB. 23 (2013) 1548–1553. https://doi.org/10.1016/j.cub.2013.06.057.

284. G. Liti, A.N. Nguyen Ba, M. Blythe, C.A. Müller, A. Bergström, F.A. Cubillos, F. Dafhnis-Calas, S. Khoshraftar, S. Malla, N. Mehta, C.C. Siow, J. Warringer, A.M. Moses, E.J. Louis, C.A. Nieduszynski, High quality de novo sequencing and assembly of the Saccharomyces arboricolus genome, BMC Genomics. 14 (2013) 69. https://doi.org/10.1186/1471-2164-14-69.

285. A. A. Burmester, E. Shelest, G. Glöckner, C. Heddergott, S. Schindler, P. Staib, A. Heidel, M. Felder, Petzold, K. Szafranski, M. Feuermann, I. Pedruzzi, S. Priebe, M. Groth, R. Winkler, W. Li, O. Kniemeyer, V. Schroeckh, C. Hertweck, B. Hube, T.C. White, M. Platzer, R. Guthke, J. Heitman, J. Wöstemeyer, P.F. Zipfel, M. Monod, A.A. Brakhage, Comparative and functional genomics provide insights into the pathogenicity of dermatophytic fungi, Genome Biol. 12 (2011) R7. https://doi.org/10.1186/gb-2011-12-1-r7.

286. A. Goffeau, B.G. Barrell, H. Bussey, R.W. Davis, B. Dujon, H. Feldmann, F. Galibert, J.D. Hoheisel, C. Jacq, M. Johnston, E.J. Louis, H.W. Mewes, Y. Murakami, P. Philippsen, H. Tettelin, S.G. Oliver, Life with 6000 genes, Science. 274 (1996) 546, 563–567 https://doi.org/10.1126/science.274.5287.546.

287. M.C. Chibucos, S. Soliman, T. Gebremariam, H. Lee, S. Daugherty, J. Orvis, A.C. Shetty, J. Crabtree, T.H. Hazen, K.A. Etienne, P. Kumari, T.D. O’Connor, D.A. Rasko, S.G. Filler, C.M. Fraser, S.R. Lockhart, C.D. Skory, A.S. Ibrahim, V.M. Bruno, An integrated genomic and transcriptomic survey of mucormycosis-causing fungi, Nat. Commun. 7 (2016) 12218. https://doi.org/10.1038/ncomms12218.

288. T.W. Jeffries, I.V. Grigoriev, J. Grimwood, J.M. Laplaza, A. Aerts, A. Salamov, J. Schmutz, E. Lindquist, P. Dehal, H. Shapiro, Y.-S. Jin, V. Passoth, P.M. Richardson, Genome sequence of the lignocellulose-bioconverting and xylose-fermenting yeast Pichia stipitis, Nat. Biotechnol. 25 (2007) 319–326. https://doi.org/10.1038/nbt1290.

289. R.A. Ohm, J.F. de Jong, L.G. Lugones, A. Aerts, E. Kothe, J.E. Stajich, R.P. de Vries, E. Record, A. Levasseur, S.E. Baker, K.A. Bartholomew, P.M. Coutinho, S. Erdmann, T.J. Fowler, A.C. Gathman, V. Lombard, B. Henrissat, N. Knabe, U. Kües, W.W. Lilly, E. Lindquist, S. Lucas, J.K. Magnuson, F. Piumi, M. Raudaskoski, A. Salamov, J. Schmutz, F.W.M.R. Schwarze, P.A. vanKuyk, J.S. Horton, I.V. Grigoriev, H.A.B. Wösten, Genome sequence of the model mushroom Schizophyllum commune, Nat. Biotechnol. 28 (2010) 957–963. https://doi.org/10.1038/nbt.1643.

290. B. Min, H. Park, Y. Jang, J.-J. Kim, K.H. Kim, J. Pangilinan, A. Lipzen, R. Riley, I.V. Grigoriev, J.W. Spatafora, I.-G. Choi, Genome sequence of a white rot fungus Schizopora paradoxa KUC8140 for wood decay and mycoremediation, J. Biotechnol. 211 (2015) 42–43. https://doi.org/10.1016/j.jbiotec.2015.06.426.

291. N. Rhind, Z. Chen, M. Yassour, D.A. Thompson, B.J. Haas, N. Habib, I. Wapinski, S. Roy, M.F. Lin, D.I. Heiman, S.K. Young, K. Furuya, Y. Guo, A. Pidoux, H.M. Chen, B. Robbertse, J.M. Goldberg, K. Aoki, E.H. Bayne, A.M. Berlin, C.A. Desjardins, E. Dobbs, L. Dukaj, L. Fan, M.G. FitzGerald, C. French, S. Gujja, K. Hansen, D. Keifenheim, J.Z. Levin, R.A. Mosher, C.A. Müller, J. Pfiffner, M. Priest, C. Russ, A. Smialowska, P. Swoboda, S.M. Sykes, M. Vaughn, S. Vengrova, R. Yoder, Q. Zeng, R. Allshire, D. Baulcombe, B.W. Birren, W. Brown, K. Ekwall, M. Kellis, J. Leatherwood, H. Levin, H. Margalit, R. Martienssen, C.A. Nieduszynski, J.W. Spatafora, N. Friedman, J.Z. Dalgaard, P. Baumann, H. Niki, A. Regev, C. Nusbaum, Comparative functional genomics of the fission yeasts, Science. 332 (2011) 930–936. https://doi.org/10.1126/science.1203357.

292. V. Wood, R. Gwilliam, M.-A. Rajandream, M. Lyne, R. Lyne, A. Stewart, J. Sgouros, N. Peat, J. Hayles, S. Baker, D. Basham, S. Bowman, K. Brooks, D. Brown, S. Brown, T. Chillingworth, C. Churcher, M. Collins, R. Connor, A. Cronin, P. Davis, T. Feltwell, A. Fraser, S. Gentles, A. Goble, N. Hamlin, D. Harris, J. Hidalgo, G. Hodgson, S. Holroyd, T. Hornsby, S. Howarth, E.J. Huckle, S. Hunt, K. Jagels, K. James, L. Jones, M. Jones, S. Leather, S. McDonald, J. McLean, P. Mooney, S. Moule, K. Mungall, L. Murphy, D. Niblett, C. Odell, K. Oliver, S. O’Neil, D. Pearson, M.A. Quail, E. Rabbinowitsch, K. Rutherford, S. Rutter, D. Saunders, K. Seeger, S. Sharp, J. Skelton, M. Simmonds, R. Squares, S. Squares, K. Stevens, K. Taylor, R.G. Taylor, A. Tivey, S. Walsh, T. Warren, S. Whitehead, J. Woodward, G. Volckaert, R. Aert, J. Robben, B. Grymonprez, I. Weltjens, E. Vanstreels, M. Rieger, M. Schäfer, S. Müller-Auer, C. Gabel, M. Fuchs, A. Düsterhöft, C. Fritzc, E. Holzer, D. Moestl, H. Hilbert, K. Borzym, I. Langer, A. Beck, H. Lehrach, R. Reinhardt, T.M. Pohl, P. Eger, W. Zimmermann, H. Wedler, R. Wambutt, B. Purnelle, A. Goffeau, E. Cadieu, S. Dréano, S. Gloux, V. Lelaure, S. Mottier, F. Galibert, S.J. Aves, Z. Xiang, C. Hunt, K. Moore, S.M. Hurst, M. Lucas, M. Rochet, C. Gaillardin, V.A. Tallada, A. Garzon, G. Thode, R.R. Daga, L. Cruzado, J. Jimenez, M. Sánchez, F. del Rey, J. Benito, A. Domínguez, J.L. Revuelta, S. Moreno, J. Armstrong, S.L. Forsburg, L. Cerutti, T. Lowe, W.R. McCombie, I. Paulsen, J. Potashkin, G.V. Shpakovski, D. Ussery, B.G. Barrell, P. Nurse, L. Cerrutti, The genome sequence of Schizosaccharomyces pombe, Nature. 415 (2002) 871–880. https://doi.org/10.1038/nature724.

293. J. Amselem, C.A. Cuomo, J.A.L. van Kan, M. Viaud, E.P. Benito, A. Couloux, P.M. Coutinho, R.P. de Vries, P.S. Dyer, S. Fillinger, E. Fournier, L. Gout, M. Hahn, L. Kohn, N. Lapalu, K.M. Plummer, J.-M. Pradier, E. Quévillon, A. Sharon, A. Simon, A. ten Have, B. Tudzynski, P. Tudzynski, P. Wincker, M. Andrew, V. Anthouard, R.E. Beever, R. Beffa, I. Benoit, O. Bouzid, B. Brault, Z. Chen, M. Choquer, J. Collémare, P. Cotton, E.G. Danchin, C. Da Silva, A. Gautier, C. Giraud, T. Giraud, C. Gonzalez, S. Grossetete, U. Güldener, B. Henrissat, B.J. Howlett, C. Kodira, M. Kretschmer, A. Lappartient, M. Leroch, C. Levis, E. Mauceli, C. Neuvéglise, B. Oeser, M. Pearson, J. Poulain, N. Poussereau, H. Quesneville, C. Rascle, J. Schumacher, B. Ségurens, A. Sexton, E. Silva, C. Sirven, D.M. Soanes, N.J. Talbot, M. Templeton, C. Yandava, O. Yarden, Q. Zeng, J.A. Rollins, M.-H. Lebrun, M. Dickman, Genomic analysis of the necrotrophic fungal pathogens Sclerotinia sclerotiorum and Botrytis cinerea, PLoS Genet. 7 (2011) e1002230. https://doi.org/10.1371/journal.pgen.1002230.

294. S.V. Balasundaram, J. Hess, M.B. Durling, S.C. Moody, L. Thorbek, C. Progida, K. LaButti, A. Aerts, K. Barry, I.V. Grigoriev, L. Boddy, N. Högberg, H. Kauserud, D.C. Eastwood, I. Skrede, The fungus that came in from the cold: dry rot’s pre-adapted ability to invade buildings, ISME J. 12 (2018) 791–801. https://doi.org/10.1038/s41396-017-0006-8.

295. D.C. Eastwood, D. Floudas, M. Binder, A. Majcherczyk, P. Schneider, A. Aerts, F.O. Asiegbu, S.E. Baker, K. Barry, M. Bendiksby, M. Blumentritt, P.M. Coutinho, D. Cullen, R.P. de Vries, A. Gathman, B. Goodell, B. Henrissat, K. Ihrmark, H. Kauserud, A. Kohler, K. LaButti, A. Lapidus, J.L. Lavin, Y.-H. Lee, E. Lindquist, W. Lilly, S. Lucas, E. Morin, C. Murat, J.A. Oguiza, J. Park, A.G. Pisabarro, R. Riley, A. Rosling, A. Salamov, O. Schmidt, J. Schmutz, I. Skrede, J. Stenlid, A. Wiebenga, X. Xie, U. Kües, D.S. Hibbett, D. Hoffmeister, N. Högberg, F. Martin, I.V. Grigoriev, S.C. Watkinson, The plant cell wall-decomposing machinery underlies the functional diversity of forest fungi, Science. 333 (2011) 762–765. https://doi.org/10.1126/science.1205411.

296. Y. Wang, M.M. White, S. Kvist, J.-M. Moncalvo, Genome-Wide Survey of Gut Fungi (Harpellales) Reveals the First Horizontally Transferred Ubiquitin Gene from a Mosquito Host, Mol. Biol. Evol. 33 (2016) 2544–2554. https://doi.org/10.1093/molbev/msw126.

297. A. A.A. Grum-Grzhimaylo, D.L. Falkoski, J. van den Heuvel, C.A. Valero-Jiménez, B. Min, I.-G. Choi, Lipzen, C.G. Daum, D.K. Aanen, A. Tsang, B. Henrissat, E.N. Bilanenko, R.P. de Vries, J.A.L. van Kan, I.V. Grigoriev, A.J.M. Debets, The obligate alkalophilic soda-lake fungus Sodiomyces alkalinus has shifted to a protein diet, Mol. Ecol. 27 (2018) 4808–4819. https://doi.org/10.1111/mec.14912.

298. G.M.N. Benucci, S. Haridas, K. Labutti, G. Marozzi, L. Antonielli, S. Sanchez, P. Marco, X. Wang, K. Barry, A. Lipzen, M. Chovatia, H. Hundley, L. Baciarelli Falini, C. Murat, F. Martin, E. Albertini, D. Donnini, I.V. Grigoriev, G. Bonito, Draft Genome Sequence of the Ectomycorrhizal Ascomycete Sphaerosporella brunnea, Microbiol. Resour. Announc. 8 (2019). https://doi.org/10.1128/MRA.00857-19.

299. C. Russ, B.F. Lang, Z. Chen, S. Gujja, T. Shea, Q. Zeng, S. Young, C.A. Cuomo, C. Nusbaum, Genome Sequence of Spizellomyces punctatus, Genome Announc. 4 (2016). https://doi.org/10.1128/genomeA.00849-16.

300. J. Schirawski, G. Mannhaupt, K. Münch, T. Brefort, K. Schipper, G. Doehlemann, M. Di Stasio, N. Rössel, A. Mendoza-Mendoza, D. Pester, O. Müller, B. Winterberg, E. Meyer, H. Ghareeb, T. Wollenberg, M. Münsterkötter, P. Wong, M. Walter, E. Stukenbrock, U. Güldener, R. Kahmann, Pathogenicity determinants in smut fungi revealed by genome comparison, Science. 330 (2010) 1546–1548. https://doi.org/10.1126/science.1195330.

301. J.K. Hane, R.G.T. Lowe, P.S. Solomon, K.-C. Tan, C.L. Schoch, J.W. Spatafora, P.W. Crous, C. Kodira, B.W. Birren, J.E. Galagan, S.F.F. Torriani, B.A. McDonald, R.P. Oliver, Dothideomycete plant interactions illuminated by genome sequencing and EST analysis of the wheat pathogen Stagonospora nodorum, Plant Cell. 19 (2007) 3347–3368. https://doi.org/10.1105/tpc.107.052829.

302. M.E.E. Franco, S. López, R. Medina, M.C.N. Saparrat, P. Balatti, Draft Genome Sequence and Gene Annotation of Stemphylium lycopersici Strain CIDEFI-216, Genome Announc. 3 (2015). https://doi.org/10.1128/genomeA.01069-15.

303. S. Branco, P. Gladieux, C.E. Ellison, A. Kuo, K. LaButti, A. Lipzen, I.V. Grigoriev, H.-L. Liao, R. Vilgalys, K.G. Peay, J.W. Taylor, T.D. Bruns, Genetic isolation between two recently diverged populations of a symbiotic fungus, Mol. Ecol. 24 (2015) 2747–2758. https://doi.org/10.1111/mec.13132.

304. S. Varriale, J. Houbraken, Z. Granchi, O. Pepe, G. Cerullo, V. Ventorino, T. Chin-A-Woeng, M. Meijer, R. Riley, I.V. Grigoriev, B. Henrissat, R.P. de Vries, V. Faraco, Talaromyces borbonicus, sp. nov., a novel fungus from biodegraded Arundo donax with potential abilities in lignocellulose conversion, Mycologia. 110 (2018) 316–324. https://doi.org/10.1080/00275514.2018.1456835.

305. W.C. Nierman, N.D. Fedorova-Abrams, A. Andrianopoulos, Genome Sequence of the AIDS-Associated Pathogen Penicillium marneffei (ATCC18224) and Its Near Taxonomic Relative Talaromyces stipitatus (ATCC10500), Genome Announc. 3 (2015). https://doi.org/10.1128/genomeA.01559-14.

306. O.H. Cissé, J.M.G.C.F. Almeida, A. Fonseca, A.A. Kumar, J. Salojärvi, K. Overmyer, P.M. Hauser, M. Pagni, Genome sequencing of the plant pathogen Taphrina deformans, the causal agent of peach leaf curl, MBio. 4 (2013) e00055–00013. https://doi.org/10.1128/mBio.00055-13.

307. N.P. McHunu, K. Permaul, A.Y. Abdul Rahman, J.A. Saito, S. Singh, M. Alam, Xylanase Superproducer: Genome Sequence of a Compost-Loving Thermophilic Fungus, Thermomyces lanuginosus Strain SSBP, Genome Announc. 1 (2013). https://doi.org/10.1128/genomeA.00388-13.

308. K.E. Bushley, R. Raja, P. Jaiswal, J.S. Cumbie, M. Nonogaki, A.E. Boyd, C.A. Owensby, B.J. Knaus, J. Elser, D. Miller, Y. Di, K.L. McPhail, J.W. Spatafora, The genome of tolypocladium inflatum: evolution, organization, and expression of the cyclosporin biosynthetic gene cluster, PLoS Genet. 9 (2013) e1003496. https://doi.org/10.1371/journal.pgen.1003496.

309. Z. Granchi, M. Peng, T. Chi-A-Woeng, R.P. de Vries, K. Hildén, M.R. Mäkelä, Genome Sequence of the Basidiomycete White-Rot Fungus Trametes pubescens FBCC735, Genome Announc. 5 (2017). https://doi.org/10.1128/genomeA.01643-16.

310. R.H. Proctor, S.P. McCormick, H.-S. Kim, R.E. Cardoza, A.M. Stanley, L. Lindo, A. Kelly, D.W. Brown, T. Lee, M.M. Vaughan, N.J. Alexander, M. Busman, S. Gutiérrez, Evolution of structural diversity of trichothecenes, a family of toxins produced by plant pathogenic and entomopathogenic fungi, PLoS Pathog. 14 (2018) e1006946. https://doi.org/10.1371/journal.ppat.1006946.

311. I.S. Druzhinina, K. Chenthamara, J. Zhang, L. Atanasova, D. Yang, Y. Miao, M.J. Rahimi, M. Grujic, F. Cai, S. Pourmehdi, K.A. Salim, C. Pretzer, A.G. Kopchinskiy, B. Henrissat, A. Kuo, H. Hundley, M. Wang, A. Aerts, A. Salamov, A. Lipzen, K. LaButti, K. Barry, I.V. Grigoriev, Q. Shen, C.P. Kubicek, Massive lateral transfer of genes encoding plant cell wall-degrading enzymes to the mycoparasitic fungus Trichoderma from its plant-associated hosts, PLoS Genet. 14 (2018) e1007322. https://doi.org/10.1371/journal.pgen.1007322.

312. C.P. Kubicek, A.S. Steindorff, K. Chenthamara, G. Manganiello, B. Henrissat, J. Zhang, F. Cai, A.G. Kopchinskiy, E.M. Kubicek, A. Kuo, R. Baroncelli, S. Sarrocco, E.F. Noronha, G. Vannacci, Q. Shen, I.V. Grigoriev, I.S. Druzhinina, Evolution and comparative genomics of the most common Trichoderma species, BMC Genomics. 20 (2019) 485. https://doi.org/10.1186/s12864-019-5680-7.

313. F. Fanelli, V.C. Liuzzi, A.F. Logrieco, C. Altomare, Genomic characterization of Trichoderma atrobrunneum (T. harzianum species complex) ITEM 908: insight into the genetic endowment of a multi-target biocontrol strain, BMC Genomics. 19 (2018) 662. https://doi.org/10.1186/s12864-018-5049-3.

314. A. C.P. Kubicek, A. Herrera-Estrella, V. Seidl-Seiboth, D.A. Martinez, I.S. Druzhinina, M. Thon, S. Zeilinger, S. Casas-Flores, B.A. Horwitz, P.K. Mukherjee, M. Mukherjee, L. Kredics, L.D. Alcaraz, Aerts, Z. Antal, L. Atanasova, M.G. Cervantes-Badillo, J. Challacombe, O. Chertkov, K. McCluskey, F. Coulpier, N. Deshpande, H. von Döhren, D.J. Ebbole, E.U. Esquivel-Naranjo, E. Fekete, M. Flipphi, F. Glaser, E.Y. Gómez-Rodríguez, S. Gruber, C. Han, B. Henrissat, R. Hermosa, M. Hernández-Oñate, L. Karaffa, I. Kosti, S. Le Crom, E. Lindquist, S. Lucas, M. Lübeck, P.S. Lübeck, A. Margeot, B. Metz, M. Misra, H. Nevalainen, M. Omann, N. Packer, G. Perrone, E.E. Uresti-Rivera, A. Salamov, M. Schmoll, B. Seiboth, H. Shapiro, S. Sukno, J.A. Tamayo-Ramos, D. Tisch, A. Wiest, H.H. Wilkinson, M. Zhang, P.M. Coutinho, C.M. Kenerley, E. Monte, S.E. Baker, I.V. Grigoriev, Comparative genome sequence analysis underscores mycoparasitism as the ancestral life style of Trichoderma, Genome Biol. 12 (2011) R40. https://doi.org/10.1186/gb-2011-12-4-r40.

315. D.J. Studholme, B. Harris, K. Le Cocq, R. Winsbury, V. Perera, L. Ryder, J.L. Ward, M.H. Beale, C.R. Thornton, M. Grant, Investigating the beneficial traits of Trichoderma hamatum GD12 for sustainable agriculture-insights from genomics, Front. Plant Sci. 4 (2013) 258. https://doi.org/10.3389/fpls.2013.00258.

316. D. Yang, K. Pomraning, A. Kopchinskiy, R. Karimi Aghcheh, L. Atanasova, K. Chenthamara, S.E. Baker, R. Zhang, Q. Shen, M. Freitag, C.P. Kubicek, I.S. Druzhinina, Genome Sequence and Annotation of Trichoderma parareesei, the Ancestor of the Cellulase Producer Trichoderma reesei, Genome Announc. 3 (2015). https://doi.org/10.1128/genomeA.00885-15.

317. T. Marik, P. Urbán, C. Tyagi, A. Szekeres, B. Leitgeb, M. Vágvölgyi, L. Manczinger, I.S. Druzhinina, C. Vágvölgyi, L. Kredics, Diversity Profile and Dynamics of Peptaibols Produced by Green Mould Trichoderma Species in Interactions with Their Hosts Agaricus bisporus and Pleurotus ostreatus, Chem. Biodivers. 14 (2017). https://doi.org/10.1002/cbdv.201700033.

318. W.-C. Li, C.-H. Huang, C.-L. Chen, Y.-C. Chuang, S.-Y. Tung, T.-F. Wang, Trichoderma reesei complete genome sequence, repeat-induced point mutation, and partitioning of CAZyme gene clusters, Biotechnol. Biofuels. 10 (2017) 170. https://doi.org/10.1186/s13068-017-0825-x.

319. E. Jourdier, L. Baudry, D. Poggi-Parodi, Y. Vicq, R. Koszul, A. Margeot, M. Marbouty, F. Bidard, Proximity ligation scaffolding and comparison of two Trichoderma reesei strains genomes, Biotechnol. Biofuels. 10 (2017) 151. https://doi.org/10.1186/s13068-017-0837-6.

320. D. Martinez, R.M. Berka, B. Henrissat, M. Saloheimo, M. Arvas, S.E. Baker, J. Chapman, O. Chertkov, P.M. Coutinho, D. Cullen, E.G.J. Danchin, I.V. Grigoriev, P. Harris, M. Jackson, C.P. Kubicek, C.S. Han, I. Ho, L.F. Larrondo, A.L. de Leon, J.K. Magnuson, S. Merino, M. Misra, B. Nelson, N. Putnam, B. Robbertse, A.A. Salamov, M. Schmoll, A. Terry, N. Thayer, A. Westerholm-Parvinen, C.L. Schoch, J. Yao, R. Barabote, R. Barbote, M.A. Nelson, C. Detter, D. Bruce, C.R. Kuske, G. Xie, P. Richardson, D.S. Rokhsar, S.M. Lucas, E.M. Rubin, N. Dunn-Coleman, M. Ward, T.S. Brettin, Genome sequencing and analysis of the biomass-degrading fungus Trichoderma reesei (syn. Hypocrea jecorina), Nat. Biotechnol. 26 (2008) 553–560. https://doi.org/10.1038/nbt1403.

321. R. Yang, J. Ao, W. Wang, K. Song, R. Li, D. Wang, Disseminated trichosporonosis in China, Mycoses. 46 (2003) 519–523. https://doi.org/10.1046/j.1439-0507.2003.00920.x.

322. R.Y. Yang, H.T. Li, H. Zhu, G.P. Zhou, M. Wang, L. Wang, Genome sequence of the Trichosporon asahii environmental strain CBS 8904, Eukaryot. Cell. 11 (2012) 1586–1587. https://doi.org/10.1128/EC.00264-12.

323. R. Kourist, F. Bracharz, J. Lorenzen, O.N. Kracht, M. Chovatia, C. Daum, S. Deshpande, A. Lipzen, M. Nolan, R.A. Ohm, I.V. Grigoriev, S. Sun, J. Heitman, T. Brück, M. Nowrousian, Genomics and Transcriptomics Analyses of the Oil-Accumulating Basidiomycete Yeast Trichosporon oleaginosus: Insights into Substrate Utilization and Alternative Evolutionary Trajectories of Fungal Mating Systems, MBio. 6 (2015) e00918. https://doi.org/10.1128/mBio.00918-15.

324. F. Martin, A. Kohler, C. Murat, R. Balestrini, P.M. Coutinho, O. Jaillon, B. Montanini, E. Morin, B. Noel, R. Percudani, B. Porcel, A. Rubini, A. Amicucci, J. Amselem, V. Anthouard, S. Arcioni, F. Artiguenave, J.-M. Aury, P. Ballario, A. Bolchi, A. Brenna, A. Brun, M. Buée, B. Cantarel, G. Chevalier, A. Couloux, C. Da Silva, F. Denoeud, S. Duplessis, S. Ghignone, B. Hilselberger, M. Iotti, B. Marçais, A. Mello, M. Miranda, G. Pacioni, H. Quesneville, C. Riccioni, R. Ruotolo, R. Splivallo, V. Stocchi, E. Tisserant, A.R. Viscomi, A. Zambonelli, E. Zampieri, B. Henrissat, M.-H. Lebrun, F. Paolocci, P. Bonfante, S. Ottonello, P. Wincker, Périgord black truffle genome uncovers evolutionary origins and mechanisms of symbiosis, Nature. 464 (2010) 1033–1038. https://doi.org/10.1038/nature08867.

325. T. Kumagai, T. Ishii, G. Terai, M. Umemura, M. Machida, K. Asai, Genome Sequence of Ustilaginoidea virens IPU010, a Rice Pathogenic Fungus Causing False Smut, Genome Announc. 4 (2016). https://doi.org/10.1128/genomeA.00306-16.

326. J.D. Laurie, S. Ali, R. Linning, G. Mannhaupt, P. Wong, U. Güldener, M. Münsterkötter, R. Moore, R. Kahmann, G. Bakkeren, J. Schirawski, Genome comparison of barley and maize smut fungi reveals targeted loss of RNA silencing components and species-specific presence of transposable elements, Plant Cell. 24 (2012) 1733–1745. https://doi.org/10.1105/tpc.112.097261.

327. J. Kämper, R. Kahmann, M. Bölker, L.-J. Ma, T. Brefort, B.J. Saville, F. Banuett, J.W. Kronstad, S.E. Gold, O. Müller, M.H. Perlin, H.A.B. Wösten, R. de Vries, J. Ruiz-Herrera, C.G. Reynaga-Peña, K. Snetselaar, M. McCann, J. Pérez-Martín, M. Feldbrügge, C.W. Basse, G. Steinberg, J.I. Ibeas, W. Holloman, P. Guzman, M. Farman, J.E. Stajich, R. Sentandreu, J.M. González-Prieto, J.C. Kennell, L. Molina, J. Schirawski, A. Mendoza-Mendoza, D. Greilinger, K. Münch, N. Rössel, M. Scherer, M. Vranes, O. Ladendorf, V. Vincon, U. Fuchs, B. Sandrock, S. Meng, E.C.H. Ho, M.J. Cahill, K.J. Boyce, J. Klose, S.J. Klosterman, H.J. Deelstra, L. Ortiz-Castellanos, W. Li, P. Sanchez-Alonso, P.H. Schreier, I. Häuser-Hahn, M. Vaupel, E. Koopmann, G. Friedrich, H. Voss, T. Schlüter, J. Margolis, D. Platt, C. Swimmer, A. Gnirke, F. Chen, V. Vysotskaia, G. Mannhaupt, U. Güldener, M. Münsterkötter, D. Haase, M. Oesterheld, H.-W. Mewes, E.W. Mauceli, D. DeCaprio, C.M. Wade, J. Butler, S. Young, D.B. Jaffe, S. Calvo, C. Nusbaum, J. Galagan, B.W. Birren, Insights from the genome of the biotrophic fungal plant pathogen Ustilago maydis, Nature. 444 (2006) 97–101. https://doi.org/10.1038/nature05248.

328. C.H. Deng, K.M. Plummer, D.A.B. Jones, C.H. Mesarich, J. Shiller, A.P. Taranto, A.J. Robinson, P. Kastner, N.E. Hall, M.D. Templeton, J.K. Bowen, Comparative analysis of the predicted secretomes of Rosaceae scab pathogens Venturia inaequalis and V. pirina reveals expanded effector families and putative determinants of host range, BMC Genomics. 18 (2017) 339. https://doi.org/10.1186/s12864-017-3699-1.

329. S.J. Klosterman, K.V. Subbarao, S. Kang, P. Veronese, S.E. Gold, B.P.H.J. Thomma, Z. Chen, B. Henrissat, Y.-H. Lee, J. Park, M.D. Garcia-Pedrajas, D.J. Barbara, A. Anchieta, R. de Jonge, P. Santhanam, K. Maruthachalam, Z. Atallah, S.G. Amyotte, Z. Paz, P. Inderbitzin, R.J. Hayes, D.I. Heiman, S. Young, Q. Zeng, R. Engels, J. Galagan, C.A. Cuomo, K.F. Dobinson, L.-J. Ma, Comparative genomics yields insights into niche adaptation of plant vascular wilt pathogens, PLoS Pathog. 7 (2011) e1002137. https://doi.org/10.1371/journal.ppat.1002137.

330. D. Bao, M. Gong, H. Zheng, M. Chen, L. Zhang, H. Wang, J. Jiang, L. Wu, Y. Zhu, G. Zhu, Y. Zhou, C. Li, S. Wang, Y. Zhao, G. Zhao, Q. Tan, Sequencing and comparative analysis of the straw mushroom (Volvariella volvacea) genome, PloS One. 8 (2013) e58294. https://doi.org/10.1371/journal.pone.0058294.

331. J. Zajc, Y. Liu, W. Dai, Z. Yang, J. Hu, C. Gostinčar, N. Gunde-Cimerman, Genome and transcriptome sequencing of the halophilic fungus Wallemia ichthyophaga: haloadaptations present and absent, BMC Genomics. 14 (2013) 617. https://doi.org/10.1186/1471-2164-14-617.

332. M. Padamsee, T.K.A. Kumar, R. Riley, M. Binder, A. Boyd, A.M. Calvo, K. Furukawa, C. Hesse, S. Hohmann, T.Y. James, K. LaButti, A. Lapidus, E. Lindquist, S. Lucas, K. Miller, S. Shantappa, I.V. Grigoriev, D.S. Hibbett, D.J. McLaughlin, J.W. Spatafora, M.C. Aime, The genome of the xerotolerant mold Wallemia sebi reveals adaptations to osmotic stress and suggests cryptic sexual reproduction, Fungal Genet. Biol. FG B. 49 (2012) 217–226. https://doi.org/10.1016/j.fgb.2012.01.007.

333. R. Gazis, A. Kuo, R. Riley, K. LaButti, A. Lipzen, J. Lin, M. Amirebrahimi, C.N. Hesse, J.W. Spatafora, B. Henrissat, M. Hainaut, I.V. Grigoriev, D.S. Hibbett, The genome of Xylona heveae provides a window into fungal endophytism, Fungal Biol. 120 (2016) 26–42. https://doi.org/10.1016/j.funbio.2015.10.002.

334. C. Magnan, J. Yu, I. Chang, E. Jahn, Y. Kanomata, J. Wu, M. Zeller, M. Oakes, P. Baldi, S. Sandmeyer, Sequence Assembly of Yarrowia lipolytica Strain W29/CLIB89 Shows Transposable Element Diversity, PloS One. 11 (2016) e0162363. https://doi.org/10.1371/journal.pone.0162363.

335. K.R. Pomraning, E.L. Bredeweg, E.J. Kerkhoven, K. Barry, S. Haridas, H. Hundley, K. LaButti, A. Lipzen, M. Yan, J.K. Magnuson, B.A. Simmons, I.V. Grigoriev, J. Nielsen, S.E. Baker, Regulation of Yeast-to-Hyphae Transition in Yarrowia lipolytica, MSphere. 3 (2018). https://doi.org/10.1128/mSphere.00541-18.

336. C. Walker, S. Ryu, S. Haridas, H. Na, M. Zane, K. LaButti, K. Barry, I.V. Grigoriev, C.T. Trinh, Draft Genome Assemblies of Ionic Liquid-Resistant Yarrowia lipolytica PO1f and Its Superior Evolved Strain, YlCW001, Microbiol. Resour. Announc. 9 (2020). https://doi.org/10.1128/MRA.01356-19.

337. C. Walker, S. Ryu, H. Na, M. Zane, K. LaButti, A. Lipzen, S. Haridas, K. Barry, I.V. Grigoriev, J. Quarterman, P. Slininger, B. Dien, C.T. Trinh, Draft Genome Assemblies of Five Robust Yarrowia lipolytica Strains Exhibiting High Lipid Production, Pentose Sugar Utilization, and Sugar Alcohol Secretion from Undetoxified Lignocellulosic Biomass Hydrolysates, Microbiol. Resour. Announc. 7 (2018). https://doi.org/10.1128/MRA.01040-18.

338. Génolevures Consortium, J.-L. Souciet, B. Dujon, C. Gaillardin, M. Johnston, P.V. Baret, P. Cliften, D.J. Sherman, J. Weissenbach, E. Westhof, P. Wincker, C. Jubin, J. Poulain, V. Barbe, B. Ségurens, F. Artiguenave, V. Anthouard, B. Vacherie, M.-E. Val, R.S. Fulton, P. Minx, R. Wilson, P. Durrens, G. Jean, C. Marck, T. Martin, M. Nikolski, T. Rolland, M.-L. Seret, S. Casarégola, L. Despons, C. Fairhead, G. Fischer, I. Lafontaine, V. Leh, M. Lemaire, J. de Montigny, C. Neuvéglise, A. Thierry, I. Blanc-Lenfle, C. Bleykasten, J. Diffels, E. Fritsch, L. Frangeul, A. Goëffon, N. Jauniaux, R. Kachouri-Lafond, C. Payen, S. Potier, L. Pribylova, C. Ozanne, G.-F. Richard, C. Sacerdot, M.-L. Straub, E. Talla, Comparative genomics of protoploid Saccharomycetaceae, Genome Res. 19 (2009) 1696–1709. https://doi.org/10.1101/gr.091546.109.

339. E.H. Stukenbrock, F.B. Christiansen, T.T. Hansen, J.Y. Dutheil, M.H. Schierup, Fusion of two divergent fungal individuals led to the recent emergence of a unique widespread pathogen species, Proc. Natl. Acad. Sci. U. S. A. 109 (2012) 10954–10959. https://doi.org/10.1073/pnas.1201403109.

340. J. Grandaubert, A. Bhattacharyya, E.H. Stukenbrock, RNA-seq-Based Gene Annotation and Comparative Genomics of Four Fungal Grass Pathogens in the Genus Zymoseptoria Identify Novel Orphan Genes and Species-Specific Invasions of Transposable Elements, G3 Bethesda Md. 5 (2015) 1323–1333. https://doi.org/10.1534/g3.115.017731.

341. F. Sievers, A. Wilm, D. Dineen, T.J. Gibson, K. Karplus, W. Li, R. Lopez, H. McWilliam, M. Remmert, J. Söding, J.D. Thompson, D.G. Higgins, Fast, scalable generation of high-quality protein multiple sequence alignments using Clustal Omega, Mol. Syst. Biol. 7 (2011) 539. https://doi.org/10.1038/msb.2011.75.

342. J.J.A. Armenteros, M. Salvatore, O. Emanuelsson, O. Winther, G. von Heijne, A. Elofsson, H. Nielsen, Detecting sequence signals in targeting peptides using deep learning, Life Sci. Alliance. 2 (2019). https://doi.org/10.26508/lsa.201900429.

343. J.J. Almagro Armenteros, K.D. Tsirigos, C.K. Sønderby, T.N. Petersen, O. Winther, S. Brunak, G. von Heijne, H. Nielsen, SignalP 5.0 improves signal peptide predictions using deep neural networks, Nat. Biotechnol. 37 (2019) 420–423. https://doi.org/10.1038/s41587-019-0036-z.

344. P. Horton, K.-J. Park, T. Obayashi, N. Fujita, H. Harada, C.J. Adams-Collier, K. Nakai, WoLF PSORT: protein localization predictor, Nucleic Acids Res. 35 (2007) W585–W587. https://doi.org/10.1093/nar/gkm259.

345. M. Bhasin, G.P.S. Raghava, ESLpred: SVM-based method for subcellular localization of eukaryotic proteins using dipeptide composition and PSI-BLAST, Nucleic Acids Res. 32 (2004) W414–W419. https://doi.org/10.1093/nar/gkh350.

346. K.D. Yamada, K. Tomii, K. Katoh, Application of the MAFFT sequence alignment program to large data—reexamination of the usefulness of chained guide trees, Bioinformatics. 32 (2016) 3246–3251. https://doi.org/10.1093/bioinformatics/btw412.

347. S. Capella-Gutiérrez, J.M. Silla-Martínez, T. Gabaldón, trimAl: a tool for automated alignment trimming in large-scale phylogenetic analyses, Bioinformatics. 25 (2009) 1972–1973. https://doi.org/10.1093/bioinformatics/btp348.

348. A. Stamatakis, RAxML version 8: a tool for phylogenetic analysis and post-analysis of large phylogenies, Bioinformatics. 30 (2014) 1312–1313. https://doi.org/10.1093/bioinformatics/btu033.

349. I. Letunic, P. Bork, Interactive Tree Of Life (iTOL) v4: recent updates and new developments, Nucleic Acids Res. 47 (2019) W256–W259. https://doi.org/10.1093/nar/gkz239.

350. T.Y. James, J.E. Stajich, C.T. Hittinger, A. Rokas, Toward a Fully Resolved Fungal Tree of Life, Annu. Rev. Microbiol. 74 (2020) 291–313. https://doi.org/10.1146/annurev-micro-022020-051835.

351. T. Varga, K. Krizsán, C. Földi, B. Dima, M. Sánchez-García, S. Sánchez-Ramírez, G.J. Szöllősi, J.G. Szarkándi, V. Papp, L. Albert, W. Andreopoulos, C. Angelini, V. Antonín, K.W. Barry, N.L. Bougher, P. Buchanan, B. Buyck, V. Bense, P. Catcheside, M. Chovatia, J. Cooper, W. Dämon, D. Desjardin, P. Finy, J. Geml, S. Haridas, K. Hughes, A. Justo, D. Karasiński, I. Kautmanova, B. Kiss, S. Kocsubé, H. Kotiranta, K.M. LaButti, B.E. Lechner, K. Liimatainen, A. Lipzen, Z. Lukács, S. Mihaltcheva, L.N. Morgado, T. Niskanen, M.E. Noordeloos, R.A. Ohm, B. Ortiz-Santana, C. Ovrebo, N. Rácz, R. Riley, A. Savchenko, A. Shiryaev, K. Soop, V. Spirin, C. Szebenyi, M. Tomšovský, R.E. Tulloss, J. Uehling, I.V. Grigoriev, C. Vágvölgyi, T. Papp, F.M. Martin, O. Miettinen, D.S. Hibbett, L.G. Nagy, Megaphylogeny resolves global patterns of mushroom evolution, Nat. Ecol. Evol. 3 (2019) 668–678. https://doi.org/10.1038/s41559-019-0834-1.

352. A. Kohler, A. Kuo, L.G. Nagy, E. Morin, K.W. Barry, F. Buscot, B. Canbäck, C. Choi, N. Cichocki, A. Clum, J. Colpaert, A. Copeland, M.D. Costa, J. Doré, D. Floudas, G. Gay, M. Girlanda, B. Henrissat, S. Herrmann, J. Hess, N. Högberg, T. Johansson, H.-R. Khouja, K. LaButti, U. Lahrmann, A. Levasseur, E.A. Lindquist, A. Lipzen, R. Marmeisse, E. Martino, C. Murat, C.Y. Ngan, U. Nehls, J.M. Plett, A. Pringle, R.A. Ohm, S. Perotto, M. Peter, R. Riley, F. Rineau, J. Ruytinx, A. Salamov, F. Shah, H. Sun, M. Tarkka, A. Tritt, C. Veneault-Fourrey, A. Zuccaro, Mycorrhizal Genomics Initiative Consortium, A. Tunlid, I.V. Grigoriev, D.S. Hibbett, F. Martin, Convergent losses of decay mechanisms and rapid turnover of symbiosis genes in mycorrhizal mutualists, Nat. Genet. 47 (2015) 410–415. https://doi.org/10.1038/ng.3223.

353. S. Miyauchi, E. Kiss, A. Kuo, E. Drula, A. Kohler, M. Sánchez-García, E. Morin, B. Andreopoulos, K.W. Barry, G. Bonito, M. Buée, A. Carver, C. Chen, N. Cichocki, A. Clum, D. Culley, P.W. Crous, L. Fauchery, M. Girlanda, R.D. Hayes, Z. Kéri, K. LaButti, A. Lipzen, V. Lombard, J. Magnuson, F. Maillard, C. Murat, M. Nolan, R.A. Ohm, J. Pangilinan, M. de F. Pereira, S. Perotto, M. Peter, S. Pfister, R. Riley, Y. Sitrit, J.B. Stielow, G. Szöllősi, L. Žifčáková, M. Štursová, J.W. Spatafora, L. Tedersoo, L.-M. Vaario, A. Yamada, M. Yan, P. Wang, J. Xu, T. Bruns, P. Baldrian, R. Vilgalys, C. Dunand, B. Henrissat, I.V. Grigoriev, D. Hibbett, L.G. Nagy, F.M. Martin, Large-scale genome sequencing of mycorrhizal fungi provides insights into the early evolution of symbiotic traits, Nat. Commun. 11 (2020) 5125. https://doi.org/10.1038/s41467-020-18795-w.

354. A.P. Douglass, B. Offei, S. Braun-Galleani, A.Y. Coughlan, A.A.R. Martos, R.A. Ortiz-Merino, K.P. Byrne, K.H. Wolfe, Population genomics shows no distinction between pathogenic Candida krusei and environmental Pichia kudriavzevii: One species, four names, PLoS Pathog. 14 (2018) e1007138. https://doi.org/10.1371/journal.ppat.1007138.

355. B. Dujon, D. Sherman, G. Fischer, P. Durrens, S. Casaregola, I. Lafontaine, J. De Montigny, C. Marck, C. Neuvéglise, E. Talla, N. Goffard, L. Frangeul, M. Aigle, V. Anthouard, A. Babour, V. Barbe, S. Barnay, S. Blanchin, J.-M. Beckerich, E. Beyne, C. Bleykasten, A. Boisramé, J. Boyer, L. Cattolico, F. Confanioleri, A. De Daruvar, L. Despons, E. Fabre, C. Fairhead, H. Ferry-Dumazet, A. Groppi, F. Hantraye, C. Hennequin, N. Jauniaux, P. Joyet, R. Kachouri, A. Kerrest, R. Koszul, M. Lemaire, I. Lesur, L. Ma, H. Muller, J.-M. Nicaud, M. Nikolski, S. Oztas, O. Ozier-Kalogeropoulos, S. Pellenz, S. Potier, G.-F. Richard, M.-L. Straub, A. Suleau, D. Swennen, F. Tekaia, M. Wésolowski-Louvel, E. Westhof, B. Wirth, M. Zeniou-Meyer, I. Zivanovic, M. Bolotin-Fukuhara, A. Thierry, C. Bouchier, B. Caudron, C. Scarpelli, C. Gaillardin, J. Weissenbach, P. Wincker, J.-L. Souciet, Genome evolution in yeasts, Nature. 430 (2004) 35–44. https://doi.org/10.1038/nature02579.

356. C. Magnan, J. Yu, I. Chang, E. Jahn, Y. Kanomata, J. Wu, M. Zeller, M. Oakes, P. Baldi, S. Sandmeyer, Sequence Assembly of Yarrowia lipolytica Strain W29/CLIB89 Shows Transposable Element Diversity, PloS One. 11 (2016) e0162363. https://doi.org/10.1371/journal.pone.0162363.

357. K.R. Pomraning, E.L. Bredeweg, E.J. Kerkhoven, K. Barry, S. Haridas, H. Hundley, K. LaButti, A. Lipzen, M. Yan, J.K. Magnuson, B.A. Simmons, I.V. Grigoriev, J. Nielsen, S.E. Baker, Regulation of Yeast-to-Hyphae Transition in Yarrowia lipolytica, MSphere. 3 (2018). https://doi.org/10.1128/mSphere.00541-18.

358. C. Walker, S. Ryu, S. Haridas, H. Na, M. Zane, K. LaButti, K. Barry, I.V. Grigoriev, C.T. Trinh, Draft Genome Assemblies of Ionic Liquid-Resistant Yarrowia lipolytica PO1f and Its Superior Evolved Strain, YlCW001, Microbiol. Resour. Announc. 9 (2020). https://doi.org/10.1128/MRA.01356-19.

359. C. Walker, S. Ryu, H. Na, M. Zane, K. LaButti, A. Lipzen, S. Haridas, K. Barry, I.V. Grigoriev, J. Quarterman, P. Slininger, B. Dien, C.T. Trinh, Draft Genome Assemblies of Five Robust Yarrowia lipolytica Strains Exhibiting High Lipid Production, Pentose Sugar Utilization, and Sugar Alcohol Secretion from Undetoxified Lignocellulosic Biomass Hydrolysates, Microbiol. Resour. Announc. 7 (2018). https://doi.org/10.1128/MRA.01040-18.

360. S.J. Mondo, R.O. Dannebaum, R.C. Kuo, K.B. Louie, A.J. Bewick, K. LaButti, S. Haridas, A. Kuo, A. Salamov, S.R. Ahrendt, R. Lau, B.P. Bowen, A. Lipzen, W. Sullivan, B.B. Andreopoulos, A. Clum, E. Lindquist, C. Daum, T.R. Northen, G. Kunde-Ramamoorthy, R.J. Schmitz, A. Gryganskyi, D. Culley, J. Magnuson, T.Y. James, M.A. O’Malley, J.E. Stajich, J.W. Spatafora, A. Visel, I.V. Grigoriev, Widespread adenine N6-methylation of active genes in fungi, Nat. Genet. 49 (2017) 964–968. https://doi.org/10.1038/ng.3859.

361. Y. Wang, M.M. White, S. Kvist, J.-M. Moncalvo, Genome-Wide Survey of Gut Fungi (Harpellales) Reveals the First Horizontally Transferred Ubiquitin Gene from a Mosquito Host, Mol. Biol. Evol. 33 (2016) 2544–2554. https://doi.org/10.1093/molbev/msw126.

362. Y. Chang, A. Desirò, H. Na, L. Sandor, A. Lipzen, A. Clum, K. Barry, I.V. Grigoriev, F.M. Martin, J.E. Stajich, M.E. Smith, G. Bonito, J.W. Spatafora, Phylogenomics of Endogonaceae and evolution of mycorrhizas within Mucoromycota, New Phytol. 222 (2019) 511–525. https://doi.org/10.1111/nph.15613.

363. J.H. Skalski, T.J. Kottom, A.H. Limper, Pathobiology of Pneumocystis pneumonia: life cycle, cell wall and cell signal transduction, FEMS Yeast Res. 15 (2015). https://doi.org/10.1093/femsyr/fov046.

364. N. Corradi, D.E. Akiyoshi, H.G. Morrison, X. Feng, L.M. Weiss, S. Tzipori, P.J. Keeling, Patterns of genome evolution among the microsporidian parasites Encephalitozoon cuniculi, Antonospora locustae and Enterocytozoon bieneusi, PloS One. 2 (2007) e1277. https://doi.org/10.1371/journal.pone.0001277.

365. E. Peyretaillade, O. Gonçalves, S. Terrat, E. Dugat-Bony, P. Wincker, R.S. Cornman, J.D. Evans, F. Delbac, P. Peyret, Identification of transcriptional signals in Encephalitozoon cuniculi widespread among Microsporidia phylum: support for accurate structural genome annotation, BMC Genomics. 10 (2009) 607. https://doi.org/10.1186/1471-2164-10-607.

366. J.-F. Pombert, M. Selman, F. Burki, F.T. Bardell, L. Farinelli, L.F. Solter, D.W. Whitman, L.M. Weiss, N. Corradi, P.J. Keeling, Gain and loss of multiple functionally related, horizontally transferred genes in the reduced genomes of two microsporidian parasites, Proc. Natl. Acad. Sci. U. S. A. 109 (2012) 12638–12643. https://doi.org/10.1073/pnas.1205020109.

367. N. Corradi, J.-F. Pombert, L. Farinelli, E.S. Didier, P.J. Keeling, The complete sequence of the smallest known nuclear genome from the microsporidian Encephalitozoon intestinalis, Nat. Commun. 1 (2010) 77. https://doi.org/10.1038/ncomms1082.

368. D.E. Akiyoshi, H.G. Morrison, S. Lei, X. Feng, Q. Zhang, N. Corradi, H. Mayanja, J.K. Tumwine, P.J. Keeling, L.M. Weiss, S. Tzipori, Genomic survey of the non-cultivatable opportunistic human pathogen, Enterocytozoon bieneusi, PLoS Pathog. 5 (2009) e1000261. https://doi.org/10.1371/journal.ppat.1000261.

369. K.L. Haag, T.Y. James, J.-F. Pombert, R. Larsson, T.M.M. Schaer, D. Refardt, D. Ebert, Evolution of a morphological novelty occurred before genome compaction in a lineage of extreme parasites, Proc. Natl. Acad. Sci. U. S. A. 111 (2014) 15480–15485. https://doi.org/10.1073/pnas.1410442111.

370. C.A. Cuomo, C.A. Desjardins, M.A. Bakowski, J. Goldberg, A.T. Ma, J.J. Becnel, E.S. Didier, L. Fan, D.I. Heiman, J.Z. Levin, S. Young, Q. Zeng, E.R. Troemel, Microsporidian genome analysis reveals evolutionary strategies for obligate intracellular growth, Genome Res. 22 (2012) 2478–2488. https://doi.org/10.1101/gr.142802.112.

371. R.S. Cornman, Y.P. Chen, M.C. Schatz, C. Street, Y. Zhao, B. Desany, M. Egholm, S. Hutchison, J.S. Pettis, W.I. Lipkin, J.D. Evans, Genomic analyses of the microsporidian Nosema ceranae, an emergent pathogen of honey bees, PLoS Pathog. 5 (2009) e1000466. https://doi.org/10.1371/journal.ppat.1000466.

372. C.H. Haitjema, S.P. Gilmore, J.K. Henske, K.V. Solomon, R. de Groot, A. Kuo, S.J. Mondo, A.A. Salamov, K. LaButti, Z. Zhao, J. Chiniquy, K. Barry, H.M. Brewer, S.O. Purvine, A.T. Wright, M. Hainaut, B. Boxma, T. van Alen, J.H.P. Hackstein, B. Henrissat, S.E. Baker, I.V. Grigoriev, M.A. O’Malley, A parts list for fungal cellulosomes revealed by comparative genomics, Nat. Microbiol. 2 (2017) 17087. https://doi.org/10.1038/nmicrobiol.2017.87.

373. N.H. Youssef, M.B. Couger, C.G. Struchtemeyer, A.S. Liggenstoffer, R.A. Prade, F.Z. Najar, H.K. Atiyeh, M.R. Wilkins, M.S. Elshahed, The genome of the anaerobic fungus Orpinomyces sp. strain C1A reveals the unique evolutionary history of a remarkable plant biomass degrader, Appl. Environ. Microbiol. 79 (2013) 4620–4634. https://doi.org/10.1128/AEM.00821-13.

374. O.H. Cissé, M. Pagni, P.M. Hauser, De novo assembly of the Pneumocystis jirovecii genome from a single bronchoalveolar lavage fluid specimen from a patient, MBio. 4 (2012) e00428–00412. https://doi.org/10.1128/mBio.00428-12.

375. T.A. Nguyen, O.H. Cissé, J. Yun Wong, P. Zheng, D. Hewitt, M. Nowrousian, J.E. Stajich, G. Jedd, Innovation and constraint leading to complex multicellularity in the Ascomycota, Nat. Commun. 8 (2017) 14444. https://doi.org/10.1038/ncomms14444.

376. N. Rhind, Z. Chen, M. Yassour, D.A. Thompson, B.J. Haas, N. Habib, I. Wapinski, S. Roy, M.F. Lin, D.I. Heiman, S.K. Young, K. Furuya, Y. Guo, A. Pidoux, H.M. Chen, B. Robbertse, J.M. Goldberg, K. Aoki, E.H. Bayne, A.M. Berlin, C.A. Desjardins, E. Dobbs, L. Dukaj, L. Fan, M.G. FitzGerald, C. French, S. Gujja, K. Hansen, D. Keifenheim, J.Z. Levin, R.A. Mosher, C.A. Müller, J. Pfiffner, M. Priest, C. Russ, A. Smialowska, P. Swoboda, S.M. Sykes, M. Vaughn, S. Vengrova, R. Yoder, Q. Zeng, R. Allshire, D. Baulcombe, B.W. Birren, W. Brown, K. Ekwall, M. Kellis, J. Leatherwood, H. Levin, H. Margalit, R. Martienssen, C.A. Nieduszynski, J.W. Spatafora, N. Friedman, J.Z. Dalgaard, P. Baumann, H. Niki, A. Regev, C. Nusbaum, Comparative functional genomics of the fission yeasts, Science. 332 (2011) 930–936. https://doi.org/10.1126/science.1203357.

377. V. Wood, R. Gwilliam, M.-A. Rajandream, M. Lyne, R. Lyne, A. Stewart, J. Sgouros, N. Peat, J. Hayles, S. Baker, D. Basham, S. Bowman, K. Brooks, D. Brown, S. Brown, T. Chillingworth, C. Churcher, M. Collins, R. Connor, A. Cronin, P. Davis, T. Feltwell, A. Fraser, S. Gentles, A. Goble, N. Hamlin, D. Harris, J. Hidalgo, G. Hodgson, S. Holroyd, T. Hornsby, S. Howarth, E.J. Huckle, S. Hunt, K. Jagels, K. James, L. Jones, M. Jones, S. Leather, S. McDonald, J. McLean, P. Mooney, S. Moule, K. Mungall, L. Murphy, D. Niblett, C. Odell, K. Oliver, S. O’Neil, D. Pearson, M.A. Quail, E. Rabbinowitsch, K. Rutherford, S. Rutter, D. Saunders, K. Seeger, S. Sharp, J. Skelton, M. Simmonds, R. Squares, S. Squares, K. Stevens, K. Taylor, R.G. Taylor, A. Tivey, S. Walsh, T. Warren, S. Whitehead, J. Woodward, G. Volckaert, R. Aert, J. Robben, B. Grymonprez, I. Weltjens, E. Vanstreels, M. Rieger, M. Schäfer, S. Müller-Auer, C. Gabel, M. Fuchs, A. Düsterhöft, C. Fritzc, E. Holzer, D. Moestl, H. Hilbert, K. Borzym, I. Langer, A. Beck, H. Lehrach, R. Reinhardt, T.M. Pohl, P. Eger, W. Zimmermann, H. Wedler, R. Wambutt, B. Purnelle, A. Goffeau, E. Cadieu, S. Dréano, S. Gloux, V. Lelaure, S. Mottier, F. Galibert, S.J. Aves, Z. Xiang, C. Hunt, K. Moore, S.M. Hurst, M. Lucas, M. Rochet, C. Gaillardin, V.A. Tallada, A. Garzon, G. Thode, R.R. Daga, L. Cruzado, J. Jimenez, M. Sánchez, F. del Rey, J. Benito, A. Domínguez, J.L. Revuelta, S. Moreno, J. Armstrong, S.L. Forsburg, L. Cerutti, T. Lowe, W.R. McCombie, I. Paulsen, J. Potashkin, G.V. Shpakovski, D. Ussery, B.G. Barrell, P. Nurse, L. Cerrutti, The genome sequence of Schizosaccharomyces pombe, Nature. 415 (2002) 871–880. https://doi.org/10.1038/nature724.

378. J.M.C. Mondego, M.F. Carazzolle, G.G.L. Costa, E.F. Formighieri, L.P. Parizzi, J. Rincones, C. Cotomacci, D.M. Carraro, A.F. Cunha, H. Carrer, R.O. Vidal, R.C. Estrela, O. García, D.P.T. Thomazella, B.V. de Oliveira, A.B. Pires, M.C.S. Rio, M.R.R. Araújo, M.H. de Moraes, L.A.B. Castro, K.P. Gramacho, M.S. Gonçalves, J.P.M. Neto, A.G. Neto, L.V. Barbosa, M.J. Guiltinan, B.A. Bailey, L.W. Meinhardt, J.C. Cascardo, G.A.G. Pereira, A genome survey of Moniliophthora perniciosa gives new insights into Witches’ Broom Disease of cacao, BMC Genomics. 9 (2008) 548. https://doi.org/10.1186/1471-2164-9-548.

379. J.C. Nielsen, S. Grijseels, S. Prigent, B. Ji, J. Dainat, K.F. Nielsen, J.C. Frisvad, M. Workman, J. Nielsen, Global analysis of biosynthetic gene clusters reveals vast potential of secondary metabolite production in Penicillium species, Nat. Microbiol. 2 (2017) 17044. https://doi.org/10.1038/nmicrobiol.2017.44.

380. G.G. Moore, B.M. Mack, S.B. Beltz, M.K. Gilbert, Draft Genome Sequence of an Aflatoxigenic Aspergillus Species, A. bombycis, Genome Biol. Evol. 8 (2016) 3297–3300. https://doi.org/10.1093/gbe/evw238.

381. L.H. Okagaki, C.C. Nunes, J. Sailsbery, B. Clay, D. Brown, T. John, Y. Oh, N. Young, M. Fitzgerald, B.J. Haas, Q. Zeng, S. Young, X. Adiconis, L. Fan, J.Z. Levin, T.K. Mitchell, P.A. Okubara, M.L. Farman, L.M. Kohn, B. Birren, L.-J. Ma, R.A. Dean, Genome Sequences of Three Phytopathogenic Species of the Magnaporthaceae Family of Fungi, G3 Bethesda Md. 5 (2015) 2539–2545 https://doi.org/10.1534/g3.115.020057.

382. G. Wang, Z. Liu, R. Lin, E. Li, Z. Mao, J. Ling, Y. Yang, W.-B. Yin, B. Xie, Biosynthesis of Antibiotic Leucinostatins in Bio-control Fungus Purpureocillium lilacinum and Their Inhibition on Phytophthora Revealed by Genome Mining, PLoS Pathog. 12 (2016) e1005685. https://doi.org/10.1371/journal.ppat.1005685.

383. T. Krassowski, A.Y. Coughlan, X.-X. Shen, X. Zhou, J. Kominek, D.A. Opulente, R. Riley, I.V. Grigoriev, N. Maheshwari, D.C. Shields, C.P. Kurtzman, C.T. Hittinger, A. Rokas, K.H. Wolfe, Evolutionary instability of CUG-Leu in the genetic code of budding yeasts, Nat. Commun. 9 (2018) 1887. https://doi.org/10.1038/s41467-018-04374-7.

384. R. Yang, J. Ao, W. Wang, K. Song, R. Li, D. Wang, Disseminated trichosporonosis in China, Mycoses. 46 (2003) 519–523. https://doi.org/10.1046/j.1439-0507.2003.00920.x.

385. R.Y. Yang, H.T. Li, H. Zhu, G.P. Zhou, M. Wang, L. Wang, Genome sequence of the Trichosporon asahii environmental strain CBS 8904, Eukaryot. Cell. 11 (2012) 1586–1587. https://doi.org/10.1128/EC.00264-12.

386. D. Armaleo, O. Müller, F. Lutzoni, Ó.S. Andrésson, G. Blanc, H.B. Bode, F.R. Collart, F. Dal Grande, F. Dietrich, I.V. Grigoriev, S. Joneson, A. Kuo, P.E. Larsen, J.M. Logsdon, D. Lopez, F. Martin, S.P. May, T.R. McDonald, S.S. Merchant, V. Miao, E. Morin, R. Oono, M. Pellegrini, N. Rubinstein, M.V. Sanchez-Puerta, E. Savelkoul, I. Schmitt, J.C. Slot, D. Soanes, P. Szövényi, N.J. Talbot, C. Veneault-Fourrey, B.B. Xavier, The lichen symbiosis re-viewed through the genomes of Cladonia grayi and its algal partner Asterochloris glomerata, BMC Genomics. 20 (2019) 605. https://doi.org/10.1186/s12864-019-5629-x.

387. S.R. Ahrendt, C.A. Quandt, D. Ciobanu, A. Clum, A. Salamov, B. Andreopoulos, J.-F. Cheng, T. Woyke, A. Pelin, B. Henrissat, N.K. Reynolds, G.L. Benny, M.E. Smith, T.Y. James, I.V. Grigoriev, Leveraging single-cell genomics to expand the fungal tree of life, Nat. Microbiol. 3 (2018) 1417– 1428. https://doi.org/10.1038/s41564-018-0261-0.

388. S. Duplessis, C.A. Cuomo, Y.-C. Lin, A. Aerts, E. Tisserant, C. Veneault-Fourrey, D.L. Joly, S. Hacquard, J. Amselem, B.L. Cantarel, R. Chiu, P.M. Coutinho, N. Feau, M. Field, P. Frey, E. Gelhaye, J. Goldberg, M.G. Grabherr, C.D. Kodira, A. Kohler, U. Kües, E.A. Lindquist, S.M. Lucas, R. Mago, E. Mauceli, E. Morin, C. Murat, J.L. Pangilinan, R. Park, M. Pearson, H. Quesneville, N. Rouhier, S. Sakthikumar, A.A. Salamov, J. Schmutz, B. Selles, H. Shapiro, P. Tanguay, G.A. Tuskan, B. Henrissat, Y. Van de Peer, P. Rouzé, J.G. Ellis, P.N. Dodds, J.E. Schein, S. Zhong, R.C. Hamelin, I.V. Grigoriev, L.J. Szabo, F. Martin, Obligate biotrophy features unraveled by the genomic analysis of rust fungi, Proc. Natl. Acad. Sci. U. S. A. 108 (2011) 9166–9171. https://doi.org/10.1073/pnas.1019315108.

389. A. Nemri, D.G.O. Saunders, C. Anderson, N.M. Upadhyaya, J. Win, G.J. Lawrence, D.A. Jones, S. Kamoun, J.G. Ellis, P.N. Dodds, The genome sequence and effector complement of the flax rust pathogen Melampsora lini, Front. Plant Sci. 5 (2014) 98. https://doi.org/10.3389/fpls.2014.00098.

390. E.S. Nazareno, F. Li, M. Smith, R.F. Park, S.F. Kianian, M. Figueroa, Puccinia coronata f. sp. avenae: a threat to global oat production, Mol. Plant Pathol. 19 (2018) 1047–1060. https://doi.org/10.1111/mpp.12608.

391. B. Schwessinger, J. Sperschneider, W.S. Cuddy, D.P. Garnica, M.E. Miller, J.M. Taylor, P.N. Dodds, M. Figueroa, R.F. Park, J.P. Rathjen, A Near-Complete Haplotype-Phased Genome of the Dikaryotic Wheat Stripe Rust Fungus Puccinia striiformis f. sp. tritici Reveals High Interhaplotype Diversity, MBio. 9 (2018). https://doi.org/10.1128/mBio.02275-17.

392. D. Cantu, M. Govindarajulu, A. Kozik, M. Wang, X. Chen, K.K. Kojima, J. Jurka, R.W. Michelmore, J. Dubcovsky, Next generation sequencing provides rapid access to the genome of Puccinia striiformis f. sp. tritici, the causal agent of wheat stripe rust, PloS One. 6 (2011) e24230. https://doi.org/10.1371/journal.pone.0024230.

393. C.A. Cuomo, G. Bakkeren, H.B. Khalil, V. Panwar, D. Joly, R. Linning, S. Sakthikumar, X. Song, X. Adiconis, L. Fan, J.M. Goldberg, J.Z. Levin, S. Young, Q. Zeng, Y. Anikster, M. Bruce, M. Wang, C. Yin, B. McCallum, L.J. Szabo, S. Hulbert, X. Chen, J.P. Fellers, Comparative Analysis Highlights Variable Genome Content of Wheat Rusts and Divergence of the Mating Loci, G3 Bethesda Md. 7 (2017) 361–376. https://doi.org/10.1534/g3.116.032797.

394. L. Frantzeskakis, B. Kracher, S. Kusch, M. Yoshikawa-Maekawa, S. Bauer, C. Pedersen, P.D. Spanu, T. Maekawa, P. Schulze-Lefert, R. Panstruga, Signatures of host specialization and a recent transposable element burst in the dynamic one-speed genome of the fungal barley powdery mildew pathogen, BMC Genomics. 19 (2018) 381. https://doi.org/10.1186/s12864-018-4750-6.

395. T. Wicker, S. Oberhaensli, F. Parlange, J.P. Buchmann, M. Shatalina, S. Roffler, R. Ben-David, J. Doležel, H. Šimková, P. Schulze-Lefert, P.D. Spanu, R. Bruggmann, J. Amselem, H. Quesneville, E. Ver Loren van Themaat, T. Paape, K.K. Shimizu, B. Keller, The wheat powdery mildew genome shows the unique evolution of an obligate biotroph, Nat. Genet. 45 (2013) 1092–1096. https://doi.org/10.1038/ng.2704.

396. E. Morin, S. Miyauchi, H. San Clemente, E.C.H. Chen, A. Pelin, I. de la Providencia, S. Ndikumana, D. Beaudet, M. Hainaut, E. Drula, A. Kuo, N. Tang, S. Roy, J. Viala, B. Henrissat, I.V. Grigoriev, N. Corradi, C. Roux, F.M. Martin, Comparative genomics of Rhizophagus irregularis, R. cerebriforme, R. diaphanus and Gigaspora rosea highlights specific genetic features in Glomeromycotina, New Phytol. 222 (2019) 1584–1598. https://doi.org/10.1111/nph.15687.

397. E.C.H. Chen, E. Morin, D. Beaudet, J. Noel, G. Yildirir, S. Ndikumana, P. Charron, C. St-Onge, J. Giorgi, M. Krüger, T. Marton, J. Ropars, I.V. Grigoriev, M. Hainaut, B. Henrissat, C. Roux, F. Martin, N. Corradi, High intraspecific genome diversity in the model arbuscular mycorrhizal symbiont Rhizophagus irregularis, New Phytol. 220 (2018) 1161–1171. https://doi.org/10.1111/nph.14989.

398. V.U. Schwartze, S. Winter, E. Shelest, M. Marcet-Houben, F. Horn, S. Wehner, J. Linde, V. Valiante, M. Sammeth, K. Riege, M. Nowrousian, K. Kaerger, I.D. Jacobsen, M. Marz, A.A. Brakhage, T. Gabaldón, S. Böcker, K. Voigt, Gene expansion shapes genome architecture in the human pathogen Lichtheimia corymbifera: an evolutionary genomics analysis in the ancient terrestrial mucorales (Mucoromycotina), PLoS Genet. 10 (2014) e1004496. https://doi.org/10.1371/journal.pgen.1004496.

399. M. Antoine, A. Gand, S. Boschi-Muller, G. Branlant, Characterization of the Amino Acids from Neisseria meningitidis MsrA Involved in the Chemical Catalysis of the Methionine Sulfoxide Reduction Step, J. Biol. Chem. 281 (2006) 39062–39070. https://doi.org/10.1074/jbc.M608844200.

400. H.-Y. Kim, D.E. Fomenko, Y.-E. Yoon, V.N. Gladyshev, Catalytic advantages provided by selenocysteine in methionine-S-sulfoxide reductases, Biochemistry. 45 (2006) 13697–13704. https://doi.org/10.1021/bi0611614.

401. X.-X. Ma, P.-C. Guo, W.-W. Shi, M. Luo, X.-F. Tan, Y. Chen, C.-Z. Zhou, Structural Plasticity of the Thioredoxin Recognition Site of Yeast Methionine S-Sulfoxide Reductase Mxr1, J. Biol. Chem. 286 (2011) 13430–13437. https://doi.org/10.1074/jbc.M110.205161.

402. M. Mariotti, G. Salinas, T. Gabaldón, V.N. Gladyshev, Utilization of selenocysteine in early-branching fungal phyla, Nat. Microbiol. 4 (2019) 759–765. https://doi.org/10.1038/s41564-018-0354-9.

403. E.H. Lee, K. Lee, G.-H. Kwak, Y.S. Park, K.-J. Lee, K.Y. Hwang, H.-Y. Kim, Evidence for the dimerization-mediated catalysis of methionine sulfoxide reductase A from Clostridium oremlandii, PloS One. 10 (2015) e0131523. https://doi.org/10.1371/journal.pone.0131523.

404. H.-Y. Kim, V.N. Gladyshev, Different catalytic mechanisms in mammalian selenocysteine- and cysteine-containing methionine-R-sulfoxide reductases, PLoS Biol. 3 (2005) e375. https://doi.org/10.1371/journal.pbio.0030375.

405. F. Neiers, S. Sonkaria, A. Olry, S. Boschi-Muller, G. Branlant, Characterization of the amino acids from Neisseria meningitidis methionine sulfoxide reductase B involved in the chemical catalysis and substrate specificity of the reductase step, J. Biol. Chem. 282 (2007) 32397–32405. https://doi.org/10.1074/jbc.M704730200.

406. A. Olry, S. Boschi-Muller, M. Marraud, S. Sanglier-Cianferani, A. Van Dorsselear, G. Branlant, Characterization of the methionine sulfoxide reductase activities of PILB, a probable virulence factor from Neisseria meningitidis, J. Biol. Chem. 277 (2002) 12016–12022. https://doi.org/10.1074/jbc.M112350200.

407. F.M. Ranaivoson, F. Neiers, B. Kauffmann, S. Boschi-Muller, G. Branlant, F. Favier, Methionine sulfoxide reductase B displays a high level of flexibility, J. Mol. Biol. 394 (2009) 83–93. https://doi.org/10.1016/j.jmb.2009.08.073.

408. Z. Cao, L. Mitchell, O. Hsia, M. Scarpa, S.T. Caldwell, A.D. Alfred, A. Gennaris, J.-F. Collet, R.C. Hartley, N.J. Bulleid, Methionine sulfoxide reductase B3 requires resolving cysteine residues for full activity and can act as a stereospecific methionine oxidase, Biochem. J. 475 (2018) 827–838. https://doi.org/10.1042/BCJ20170929.

409. E. Shumilina, O. Dobrovolska, A. Dikiy, Evolution of Structural and Coordination Features Within the Methionine Sulfoxide Reductase B Family, in: M.F. Hohmann-Marriott (Ed.), Struct. Basis Biol. Energy Gener., Springer Netherlands, Dordrecht, 2014: pp. 199–215. https://doi.org/10.1007/978-94-017-8742-0_11.

410. E. Fisher, A.M. Dawson, G. Polshyna, J. Lisak, B. Crable, E. Perera, M. Ranganathan, M. Thangavelu, P. Basu, J.F. Stolz, Transformation of Inorganic and Organic Arsenic byAlkaliphilus oremlandiisp. nov. Strain OhILAs, Ann. N. Y. Acad. Sci. 1125 (2008) 230–241. https://doi.org/10.1196/annals.1419.006.

411. M.S. Fuller, R.P. Clay, Observations of Gonapodya in Pure Culture: Growth, Development and Cell Wall Characterization, Mycologia. 85 (1993) 38–45. https://doi.org/10.2307/3760475.

412. I. Letunic, P. Bork, Interactive Tree Of Life (iTOL) v4: recent updates and new developments, Nucleic Acids Res. 47 (2019) W256–W259. https://doi.org/10.1093/nar/gkz239.

413. C.L. Murphy, N.H. Youssef, R.A. Hanafy, M.B. Couger, J.E. Stajich, Y. Wang, K. Baker, S.S. Dagar, G.W. Griffith, I.F. Farag, T.M. Callaghan, M.S. Elshahed, Horizontal Gene Transfer as an Indispensable Driver for Evolution of Neocallimastigomycota into a Distinct Gut-Dwelling Fungal Lineage, Appl. Environ. Microbiol. 85 (2019). https://doi.org/10.1128/AEM.00988-19.

414. M. Nishiyama, K. Senoo, H. Wada, S. Matsumoto, Identification of soil micro-habitats for growth, death and survival of a bacterium, γ-1,2,3,4,5,6-hexachlorocyclohexane-assimilating Sphingomonas paucimobilis, by fractionation of soil, FEMS Microbiol. Lett. 101 (1992) 145–150. https://doi.org/10.1016/0378-1097(92)90810-B.

415. M. Ravenhall, N. Škunca, F. Lassalle, C. Dessimoz, Inferring Horizontal Gene Transfer, PLOS Comput. Biol. 11 (2015) e1004095. https://doi.org/10.1371/journal.pcbi.1004095.

416. B.C. Lee, D.T. Le, V.N. Gladyshev, Mammals reduce methionine-S-sulfoxide with MsrA and are unable to reduce methionine-R-sulfoxide, and this function can be restored with a yeast reductase, J. Biol. Chem. 283 (2008) 28361–28369. https://doi.org/10.1074/jbc.M805059200.

417. D.T. Le, L. Tarrago, Y. Watanabe, A. Kaya, B.C. Lee, U. Tran, R. Nishiyama, D.E. Fomenko, V.N. Gladyshev, L.-S.P. Tran, Diversity of plant methionine sulfoxide reductases B and evolution of a form specific for free methionine sulfoxide, PloS One. 8 (2013) e65637. https://doi.org/10.1371/journal.pone.0065637.

418. Y. Zhang, H. Romero, G. Salinas, V.N. Gladyshev, Dynamic evolution of selenocysteine utilization in bacteria: a balance between selenoprotein loss and evolution of selenocysteine from redox active cysteine residues, Genome Biol. 7 (2006) R94. https://doi.org/10.1186/gb-2006-7-10-r94.

419. L. Tarrago, E. Laugier, P. Rey, Protein-repairing methionine sulfoxide reductases in photosynthetic organisms: gene organization, reduction mechanisms, and physiological roles, Mol. Plant. 2 (2009) 202–217. https://doi.org/10.1093/mp/ssn067.

420. V. Tarallo, K. Sudarshan, V. Nosek, J. Míšek, Development of a simple high-throughput assay for directed evolution of enantioselective sulfoxide reductases, Chem. Commun. 56 (2020) 5386– 5388. https://doi.org/10.1039/D0CC01660H.

421. I. Pirkov, J. Norbeck, L. Gustafsson, E. Albers, A complete inventory of all enzymes in the eukaryotic methionine salvage pathway, FEBS J. 275 (2008) 4111–4120. https://doi.org/10.1111/j.1742-4658.2008.06552.x.

