## Supplementary figures and table for "Distribution of methionine sulfoxide reductases in fungi and conservation of the free-methionine-*R*-sulfoxide reductase in multicellular eukaryotes"

### SUPPLEMENTARY DATA

| | |
|---|---|
| **Figure S1.** Phylogenetic analysis of fungal MsrAs. | Page 2 |
| **Figure S2.** Phylogenetic analysis of fungal MsrBs. | Page 3 |
| **Figure S3.** Phylogenetic analysis of fungal fRMsrs. | Page 4 |
| **Figure S4.** *Gonapodya prolifera* selenocysteine-containing MsrA gene properties. | Pages 5-7 |
| **Table S1.** List of multicellular eukaryotic species possessing a *fRmsr* gene. | Page 8 |

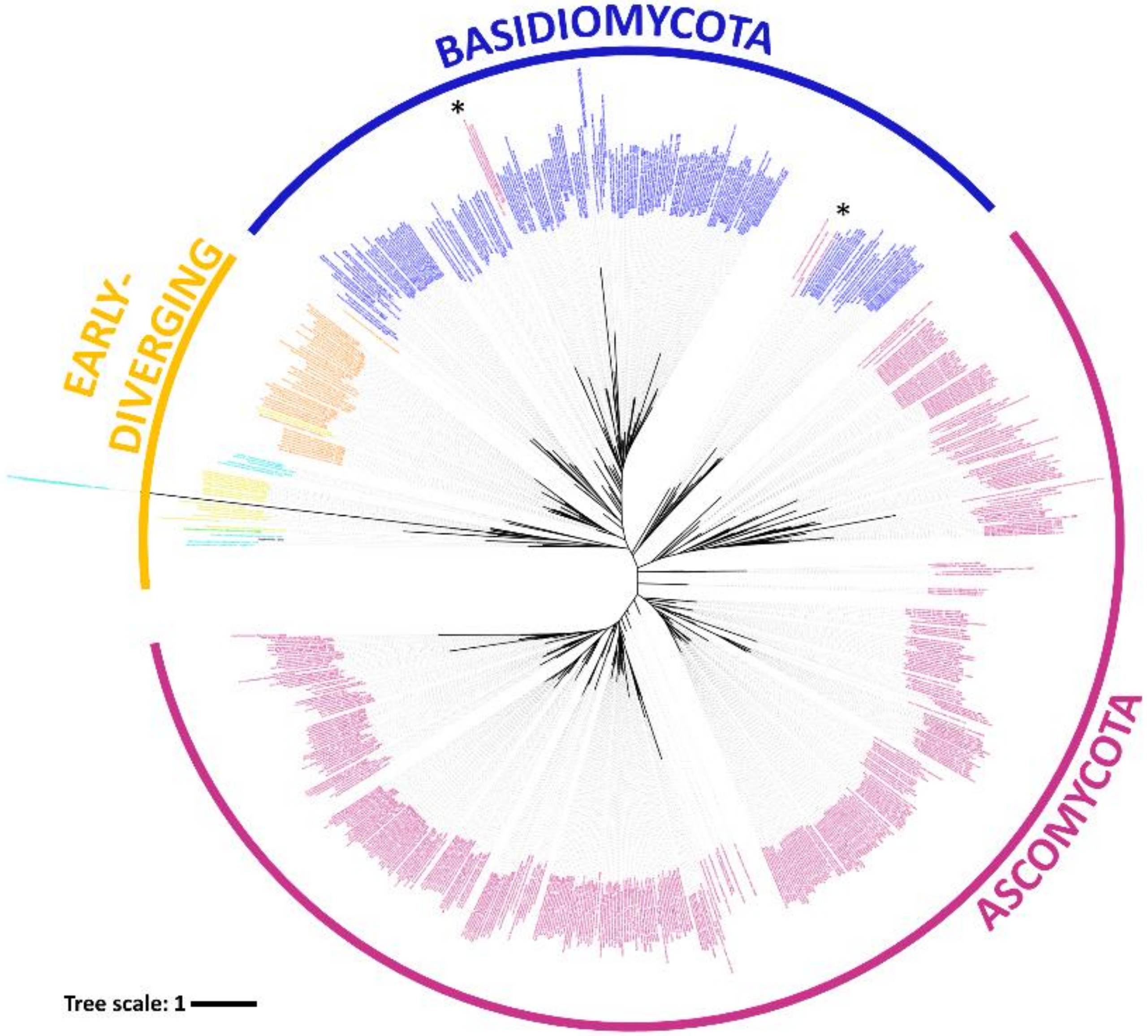


**Figure S1. Phylogenetic analysis of fungal MsrAs.** Ascomycota and Basidiomycota species labels are in *pink* and *blue*, respectively. Early-diverging fungi labels are in *black*, *light blue*, *green*, *yellow* and *orange*, for Opisthosporidia, Chytridiomycota, Blastocladiomycota, Zoopagomycota and Mucoromycota, respectively. Asterisks highlight protein sequence positions that are not congruent with the species phylogeny. The phylogenetic tree was built with 713 protein sequences using RAxML v8.2 [348] and represented using iTOL (https://itol.embl.de/) [413].

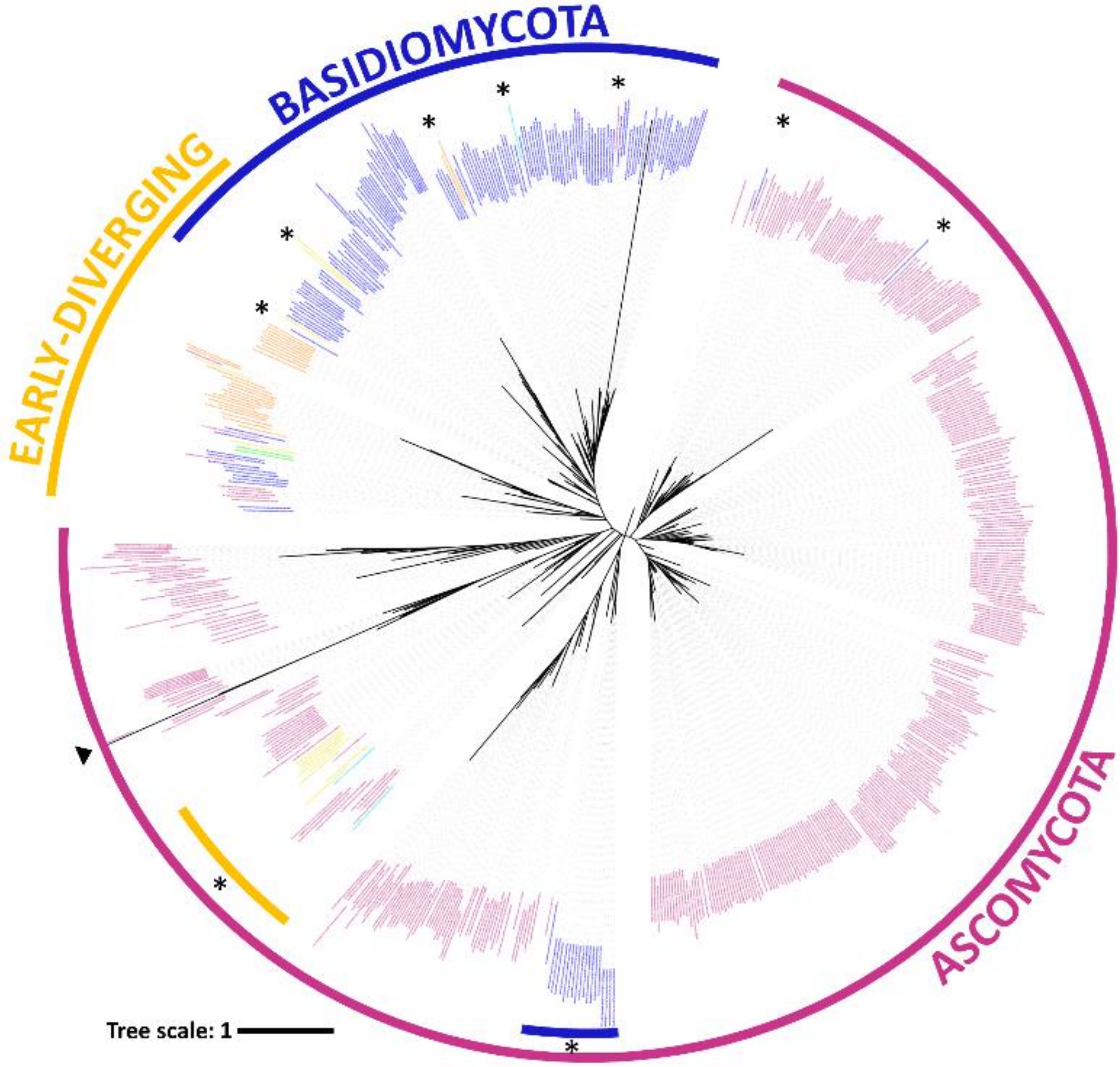


**Figure S2. Phylogenetic analysis of fungal MsrBs.** Ascomycota and Basidiomycota species labels are in *pink* and *blue*, respectively. Early-diverging fungi labels are in *black*, *light blue*, *green*, *yellow* and *orange*, for Opisthosporidia, Chytridiomycota, Blastocladiomycota, Zoopagomycota and Mucoromycota, respectively. Asterisks highlight protein sequences that are not congruent with the species phylogeny. The branch indicated by a *black* arrow was truncated for clarity. The phylogenetic tree was built with 685 protein sequences using RAxML v8.2 [348] and represented using iTOL (https://itol.embl.de/) [413].

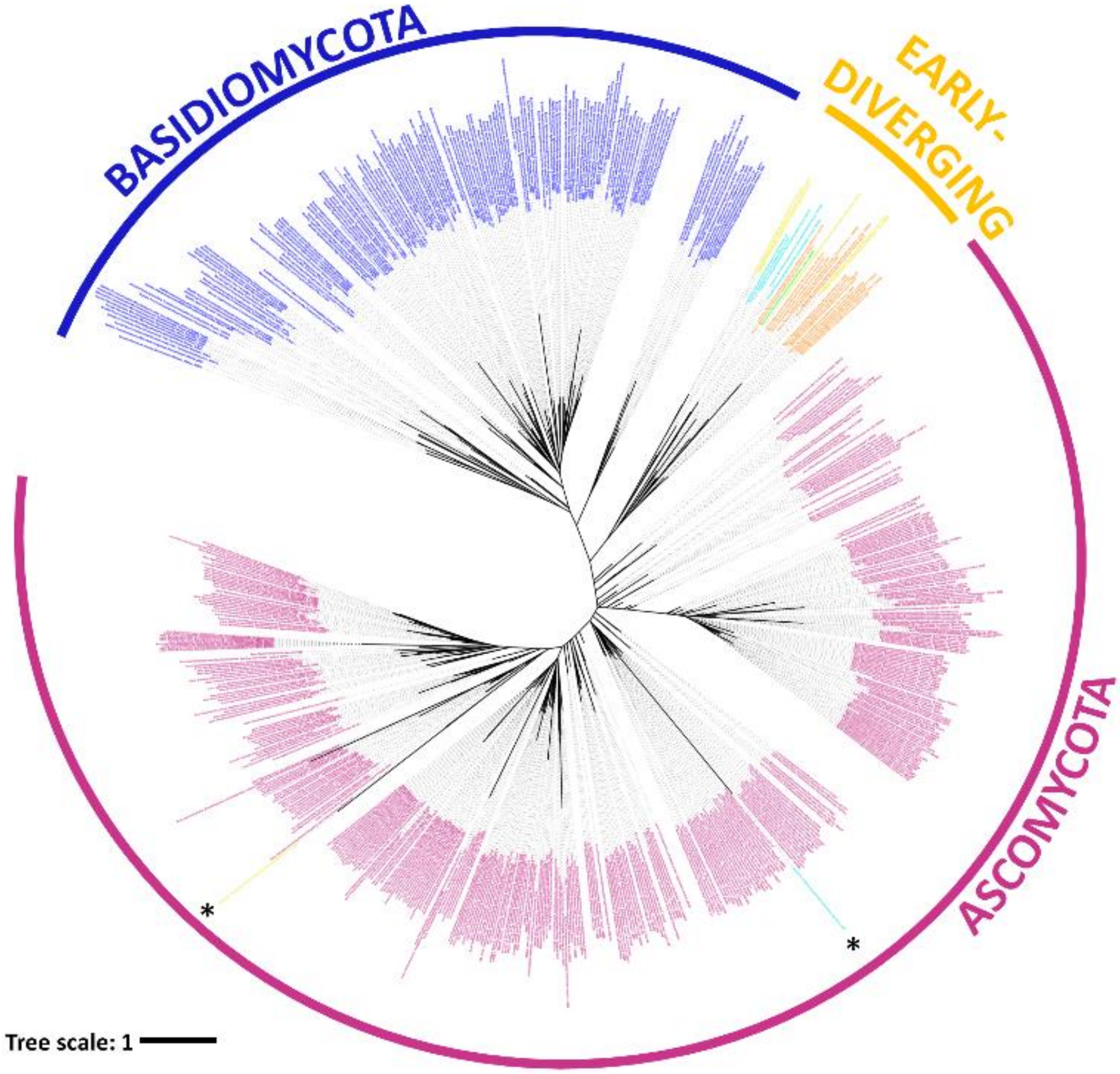


**Figure S3. Phylogenetic analysis of fungal fRMsrs.** Ascomycota and Basidiomycota species labels are in *pink* and *blue*, respectively. Early-diverging fungi labels are in *black*, *light blue*, *green*, *yellow* and *orange*, for Opisthosporidia, Chytridiomycota, Blastocladiomycota, Zoopagomycota and Mucoromycota, respectively. Asterisks highlight protein sequences that are not congruent with the species phylogeny. The phylogenetic tree was built with 589 protein sequences using RAxML v8.2 [348] and represented using iTOL (https://itol.embl.de/) [413].

#### A) MsrA #135492

**tga Sec-codon**

#### Scaffold_36:64,800-65,710 DNA

```
atgagcgggaacgcacacagcgcaacgatggagaaggcggtgttttgtatgggatgattc
tggtcgccggagccggtcttcggctcgttggtgcgttgattcccttttatcgggcaactt
gtcgacgtaaatctcacccggaacccccaccatctctccaggatggcgtcgctgctacga
gggtaggatacgctggcggaagaacacccaacccaacatgtatggtgcgaatcgtgttgt
gctcgctcccttacttgcgctcacaattccaccagattacgaccttgccgaccacaccga
atccgtcgaggtcacattcgacccatccattgtatcctacgccggcctcctccgtgtccg
tcgccaaccacccggggtattcctacacgccccatacgcctcccgtattttcgttctcac
cgacaaacaagaggcagaagcgattgccgaggtggatgcgaaccagaaaagcaaccccga
ggtggcgaccgagatgtggagaatcgccgagacgcatcatcacgatcctgcgttggcatt
tgagactggtcaggcgttagaggctgtagaagcattcacgagattgggtgtcacatcatc
aggagaattcatgtga
```

**Exon 1**

**Exon 2**

**Exon 3**

#### Scaffold_36:64,800-65,578 cDNA with translation

```
atgagcgggaacgcacacagcgcaacgatggagaaggcggtgttttgtatgggatgattc
 M  S  G  N  A  H  S  A  T  M  E  K  A  V  F  C  M  G  U  F
tggtcgccggagccggtcttcggctcgttggatggcgtcgctgctacgagggtaggatac
 W  S  P  E  P  V  F  G  S  L  D  G  V  A  A  T  R  V  G  Y
gctggcggaagaacacccaacccaacatattacgaccttgccgaccacaccgaatccgtc
 A  G  G  R  T  P  N  P  T  Y  Y  D  L  A  D  H  T  E  S  V
gaggtcacattcgacccatccattgtatcctacgccggcctcctccgtgtccgtcgccaa
 E  V  T  F  D  P  S  I  V  S  Y  A  G  L  L  R  V  R  R  Q
ccacccggggtattcctacacgccccatacgcctcccgtattttcgttctcaccgacaaa
 P  P  G  V  F  L  H  A  P  Y  A  S  R  I  F  V  L  T  D  K
caagaggcagaagcgattgccgaggtggatgcgaaccagaaaagcaaccccgaggtggcg
 Q  E  A  E  A  I  A  E  V  D  A  N  Q  K  S  N  P  E  V  A
accgagatgtggagaatcgccgagacgcatcatcacgatcctgcgttggcatttgagact
 T  E  M  W  R  I  A  E  T  H  H  H  D  P  A  L  A  F  E  T
ggtcaggcgttagaggctgtagaagcattcacgagattgggtgtcacatcatcaggagaa
 G  Q  A  L  E  A  V  E  A  F  T  R  L  G  V  T  S  S  G  E
ttcatgtga
 F  M  -
```

#### MycoCosm screen copy justifying the 3 exons structure.

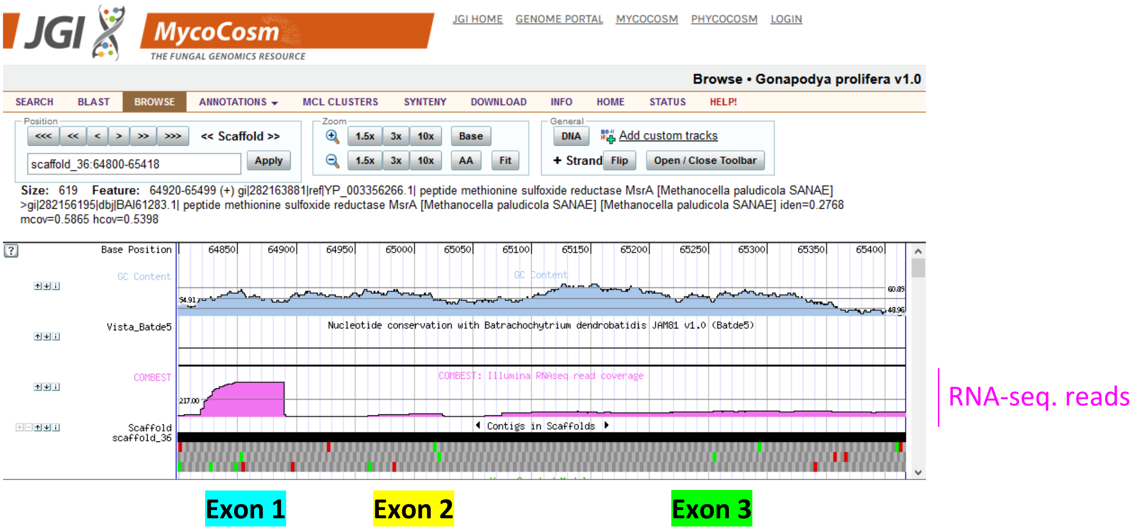

**B) MsrA #159800/159692**

tga Sec-codon

**Scaffold_105:47,377-48,491 DNA**

Exon 1, Exon 2, Exon 3, Exon 4, SECIS

```
atgaacggaaacgcaccaggcacaacgatggagaaggcagtcttcggcatgggatgattc
tggtcgcctgagccggtcttcggctcgttggtgagttgcttcccttttactgggcacctt
ctcgacgtcaatctcatcccgcacaccgatacaccccctgcagaatggcgtcgttgctacg
cgtgtaggctacgctggcggacgaacgcccaacccgacctgtacgtttcaaatcgagtgg
agctcggatccgtgcttacactcacccccgccagactacgacctcgcggaccacaccgaa
tccgtcgaagtcacatttgacccttccattgtgtcatacgccgacctcctccgtgtcttc
tttaccaaccaccacccggggtactcatacacgccccaatatgcctctcgtattttcgct
cttaccgacaaacaagaggcagaggcgattgctcaagtcgatgcgaaacagaaaaccaat
cccgaggtggcgaccgaggtgtggagaattgccgggatgcgccaggccgaattcgtgagt
tccatctcacttctcccacactctgagcttgagctcgacgtcacctcactctcgcattag
taccaccagaaattctacctccaggggcactcctcgctctcccgtctcttccccacgatc
gaatccacgcctgacaagtccacgccctccgatgacccgctcgtgagcacgacggccgcc
acgaagatgaacgcgcttgcgcatggtgtggtggaggacgtgggggagttcaggcgagcc
ctcgacgactggttcgaacaggcacaggcaaaacggaaagcttcagtgaggtgggaagag
gcgggcaaggatgcggtggagagtgcactaaagcagattgaggttcggaaccgacgcaag
gcggcagtggcgagtggaaatgggtccgagtacttcttcaccgaggacagcggcggtgga
tcgtgtggatggcgccgttgaaattgccaattggacgacacgggtgtggaagcgtttctg
agtacaactcctgaacagcacatgaagccacgccaccgaaacgcaagtgctctttgtcag
caggcatggtggcgggactgattcgcgtacata
```

**Scaffold_105:47,377-48,357 cDNA with translation**

```
atgaacggaaacgcaccaggcacaacgatggagaaggcagtcttcggcatgggatgattc
 M  N  G  N  A  P  G  T  T  M  E  K  A  V  F  G  M  G  U  F
tggtcgcctgagccggtcttcggctcgttgaatggcgtcgttgctacgcgtgtaggctac
 W  S  P  E  P  V  F  G  S  L  N  G  V  V  A  T  R  V  G  Y
gctggcggacgaacgcccaacccgacctactacgacctcgcggaccacaccgaatccgtc
 A  G  G  R  T  P  N  P  T  Y  Y  D  L  A  D  H  T  E  S  V
gaagtcacatttgacccttccattgtgtcatacgccgacctcctccgtgtcttctttacc
 E  V  T  F  D  P  S  I  V  S  Y  A  D  L  L  R  V  F  F  T
aaccaccacccggggtactcatacacgccccaatatgcctctcgtattttcgctcttacc
 N  H  H  P  G  Y  S  Y  T  P  Q  Y  A  S  R  I  F  A  L  T
gacaaacaagaggcagaggcgattgctcaagtcgatgcgaaacagaaaaccaatcccgag
 D  K  Q  E  A  E  A  I  A  Q  V  D  A  K  Q  K  T  N  P  E
gtggcgaccgaggtgtggagaattgccgggatgcgccaggccgaattctaccaccagaaa
 V  A  T  E  V  W  R  I  A  G  M  R  Q  A  E  F  Y  H  Q  K
ttctacctccaggggcactcctcgctctcccgtctcttccccacgatcgaatccacgcct
 F  Y  L  Q  G  H  S  S  L  S  R  L  F  P  T  I  E  S  T  P
gacaagtccacgccctccgatgacccgctcgtgagcacgacggccgccacgaagatgaac
 D  K  S  T  P  S  D  D  P  L  V  S  T  T  A  A  T  K  M  N
gcgcttgcgcatggtgtggtggaggacgtgggggagttcaggcgagccctcgacgactgg
 A  L  A  H  G  V  V  E  D  V  G  E  F  R  R  A  L  D  D  W
ttcgaacaggcacaggcaaaacggaaagcttcagtgaggtgggaagaggcgggcaaggat
 F  E  Q  A  Q  A  K  R  K  A  S  V  R  W  E  E  A  G  K  D
gcggtggagagtgcactaaagcagattgaggttcggaaccgacgcaaggcggcagtggcg
 A  V  E  S  A  L  K  Q  I  E  V  R  N  R  R  K  A  A  V  A
agtggaaatgggtccgagtacttcttcaccgaggacagcggcggtggatcgtgtggatgg
 S  G  N  G  S  E  Y  F  F  T  E  D  S  G  G  G  S  C  G  W
```

cgccgttga
 R  R  -

**MycoCosm screen copy justifying the 4 exons structure.**

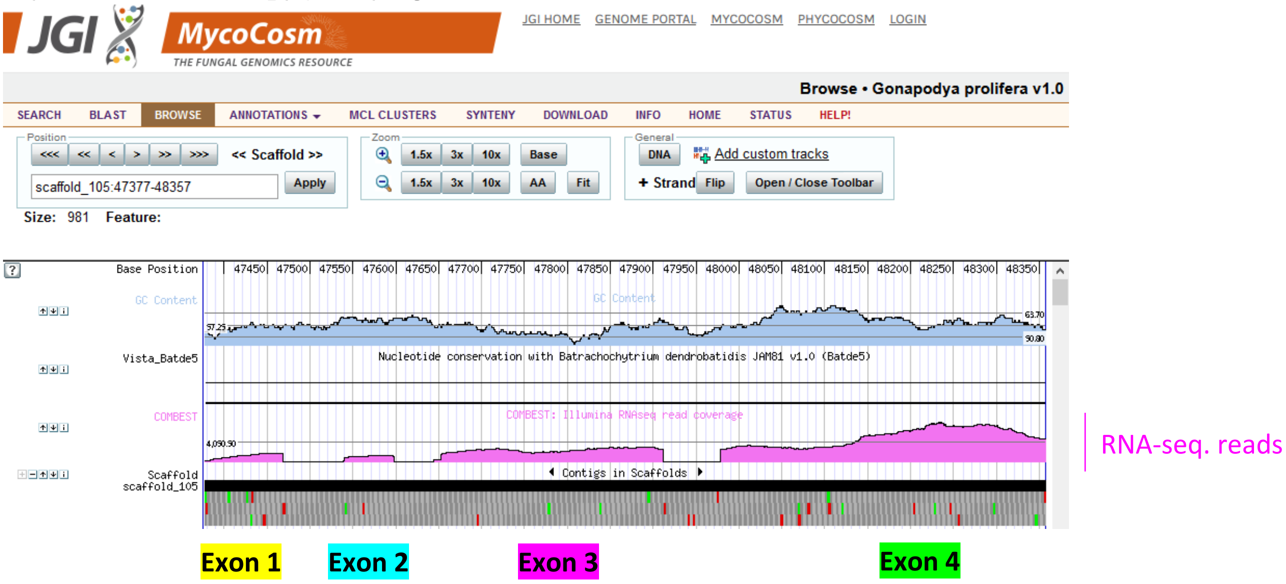


**Figure S4. *Gonapodya prolifera* selenocysteine-containing MsrA gene properties.** Gene structure for MsrA #135492 (**A**) and MsrA #159800/159692 (**B**) were based on the description made by Mariotti *et al.*, [402] and manual reconstruction using sequences and data from MycoCosm. Translations were made using Expasy Translate tool (https://web.expasy.org/translate/).

**Table S1.** List of multicellular eukaryotic species possessing a *fRmsr* gene.

| Accession | Description | Scientific Name (Common Name) | Kingdom | NCBI Taxid |
|---|---|---|---|---|
| XP_023901710.1 | free methionine-*R*-sulfoxide reductase-like | *Quercus suber* (cork oak) | Viridiplantae | 58331 |
| XP_021955388.1 | free methionine-*R*-sulfoxide reductase | *Folsomia candida* (springtail specie) | Animalia | 158441 |
| KNC29678.1 | hypothetical protein FF38_11329 | *Lucilia cuprina* (Australian sheep blowfly) | | 7375 |
| CBY16434.1 | unnamed protein product | *Oikopleura dioica* (tunicate specie) | | 34765 |
| CBY13678.1 | unnamed protein product | *Oikopleura dioica* (tunicate specie) | | 34765 |
| RWS02800.1 | diguanylate cyclase-like protein | *Dinothrombium tinctorium* (Red velvet mite) | | 1965070 |
| RWS29218.1 | diguanylate cyclase-like protein | *Leptotrombidium deliense* (mite) | | 299467 |
| ODN00976.1 | Free methionine-*R*-sulfoxide reductase | *Orchesella cincta* (springtail specie) | | 48709 |
| CAD7235481.1 | unnamed protein product | *Cyprideis torosa* (ostracod specie) | | 163714 |
| KMQ81343.1 | gaf domain nucleotide-binding protein | *Lasius niger* (black garden ant) | | 67767 |
